# Single-nucleus cross-tissue molecular reference maps to decipher disease gene function

**DOI:** 10.1101/2021.07.19.452954

**Authors:** Gokcen Eraslan, Eugene Drokhlyansky, Shankara Anand, Ayshwarya Subramanian, Evgenij Fiskin, Michal Slyper, Jiali Wang, Nicholas Van Wittenberghe, John M. Rouhana, Julia Waldman, Orr Ashenberg, Danielle Dionne, Thet Su Win, Michael S. Cuoco, Olena Kuksenko, Philip A. Branton, Jamie L. Marshall, Anna Greka, Gad Getz, Ayellet V. Segrè, François Aguet, Orit Rozenblatt-Rosen, Kristin G. Ardlie, Aviv Regev

## Abstract

Understanding the function of genes and their regulation in tissue homeostasis and disease requires knowing the cellular context in which genes are expressed in tissues across the body. Single cell genomics allows the generation of detailed cellular atlases in human tissues, but most efforts are focused on single tissue types. Here, we establish a framework for profiling multiple tissues across the human body at single-cell resolution using single nucleus RNA-Seq (snRNA-seq), and apply it to 8 diverse, archived, frozen tissue types (three donors per tissue). We apply four snRNA-seq methods to each of 25 samples from 16 donors, generating a cross-tissue atlas of 209,126 nuclei profiles, and benchmark them *vs*. scRNA-seq of comparable fresh tissues. We use a conditional variational autoencoder (cVAE) to integrate an atlas across tissues, donors, and laboratory methods. We highlight shared and tissue-specific features of tissue-resident immune cells, identifying tissue-restricted and non-restricted resident myeloid populations. These include a cross-tissue conserved dichotomy between *LYVE1-* and *HLA* class II-expressing macrophages, and the broad presence of LAM-like macrophages across healthy tissues that is also observed in disease. For rare, monogenic muscle diseases, we identify cell types that likely underlie the neuromuscular, metabolic, and immune components of these diseases, and biological processes involved in their pathology. For common complex diseases and traits analyzed by GWAS, we identify the cell types and gene modules that potentially underlie disease mechanisms. The experimental and analytical frameworks we describe will enable the generation of large-scale studies of how cellular and molecular processes vary across individuals and populations.

## INTRODUCTION

Tissue homeostasis and pathology arise from an intricate interplay between many different cell types, and disease risk is impacted by variation in genes that affect the functions and interactions of these diverse cells. Advances in human genetic studies have been instrumental in mapping tens of thousands of loci either underlying rare monogenic disease or associated with complex polygenic disease risk (Cano-Gamez & Trynka, 2020; Mills & Rahal, 2019; Tam et al., 2019), the latter mapping mostly in regulatory regions of the genome and associated as expression quantitative trait loci (eQTLs) to downstream effects on the expression of genes in *cis* and *trans* (GTEx Consortium, 2020; Võsa et al., 2018). More recently, single cell genomics has become instrumental in studying human tissue biology, with the construction of cell atlases of both healthy organs and diseased tissue, related to common disease, rare disease and cancer (Camp et al., 2019; Potter, 2018; G. Sun et al., 2021).

Coupling advances in human genetics and single cell genomics should substantially enhance our understanding of changes in the function and regulation of disease genes, because cells and tissues are the key intermediates in which disease genes act. In particular, studies have shown that tissue (GTEx Consortium, 2020), cell type (Kasela et al., 2017; Kim-Hellmuth et al., 2020; M. G. P. van der Wijst et al., 2018; Zhernakova et al., 2017), time point and stimulation (Cuomo et al., 2020; Strober et al., 2019; Ye et al., 2014) all induce a diversity of expression patterns and interactions with disease-associated genetic loci. Recent studies have combined single cell expression atlases with genetic signals (Jagadeesh et al., 2021; Skene et al., 2018; Smillie et al., 2019; Weeks et al., 2020) to associate risk genes with specific cell types and states in relevant tissues.

However, because complex diseases often manifest in and implicate cells across multiple tissues, fully realizing this opportunity requires generating atlases from diverse tissues across the body and from larger numbers of individuals, spanning different populations. This poses several challenges. First, the collection and processing of fresh tissue samples into single cell suspensions is logistically challenging and inherently difficult for some tissues, such as brain, muscle and adipose (Habib et al., 2016, 2017; Petrany et al., 2020; W. Sun et al., 2020; H. Wu et al., 2019), and hard to scale for most tissues, unless they can be pre-collected and preserved frozen. As a result, large scale single cell profiling studies in human populations (Kang et al., 2018; M. van der Wijst et al., 2020) have focused on peripheral blood mononuclear cells (PBMCs), which can be frozen and thawed for multiplexed single cell analysis. Single nucleus RNA-seq (snRNA-seq), which can be applied to frozen tissues (Delorey et al., 2021; Drokhlyansky et al., 2020), offers a compelling alternative. Second, annotation and classification of cell type and state across multiple tissues requires understanding the biological relationship between parenchymal, immune, and stromal cells across tissue types. Third, we need cross-tissue analytical frameworks, for data integration (*e.g.*, removing unwanted variation while preserving biological differences), interpretation (*e.g.*, identification of cell types and states), and synthesis with genes from studies of monogenic and complex traits.

Here, we develop a framework for snRNA-seq of multiple human tissues, and apply it to 8 archived, frozen tissue types, previously preserved as part of the GTEx project (GTEx Consortium, 2020) from the lung, skeletal muscle, heart, esophagus mucosa and muscularis, prostate, skin, and breast. We generate a cross-tissue atlas of 209,126 nuclei profiles from 25 samples, spanning 16 donors, using four snRNA-seq protocols (Drokhlyansky et al., 2020; Slyper et al., 2020). We integrate our data across tissues, donors, and protocols with a conditional variational autoencoder (cVAE) and annotate each cell subset based on literature-derived marker genes. We identify shared and tissue-specific features of tissue resident immune cells, including a dichotomy between *LYVE1*- and *HLA* class II-expressing macrophages, and the presence of lipid-associated macrophage-like cells (LAMs) across tissues. We relate rare, monogenic muscle diseases to the cell types that likely underlie the variable presentation of these diseases and nominate biological processes and cell-cell interactions involved in their pathology. We also relate common complex diseases and trait loci analyzed by GWAS to cell types and gene modules as putative disease mechanisms. Finally, we demonstrate pooling of tissues from multiple donors to illuminate a path towards profiling much larger numbers of individuals in cell atlases for human genetics and disease studies.

## RESULTS

### A multi-tissue, multi-individual single-nucleus reference atlas from archived frozen human tissues

We constructed a cross-tissue snRNA-seq atlas from 25 frozen archived tissue samples, previously collected and banked by the GTEx project, spanning 3-4 samples from each of 8 tissue sites: breast, esophagus mucosa, esophagus muscularis, heart, lung, prostate, skeletal muscle and skin from 16 individuals (7 males, 9 females) (Figure 1A). We selected the samples by RNA quality (from matching bulk tissue aliquots), tissue autolysis scores (by pathology review), and the availability of existing bulk RNA-seq and genome sequencing data (**Methods**, **Table S1**). Histology slides corresponding to each tissue were reviewed by a pathologist to provide detailed annotations, including descriptions of broad cellular composition and sample quality (**Table S1**). Because different nucleus extraction protocols can be optimal for the characteristics of different tissues (Drokhlyansky et al., 2020; Slyper et al., 2020), we isolated nuclei from each sample using four protocols (CST, NST, TST and EZ, **Methods, Table S1**), followed by droplet-based scRNA-seq (**Methods**).

**Figure 1.**
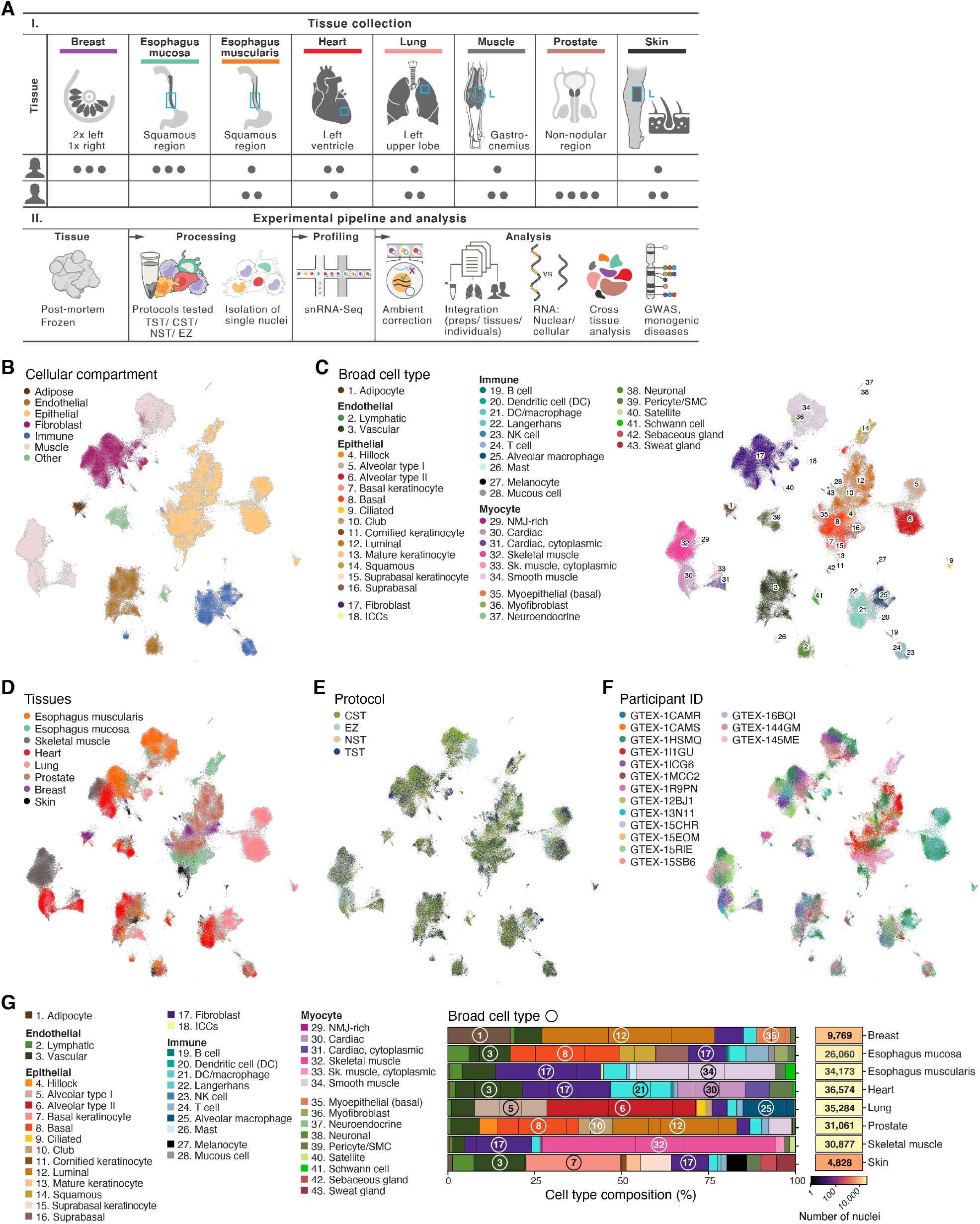
Cross tissue snRNA-seq atlas in eight adult human frozen archived tissues. **(A)** Study design. Tissue sites (I. top) and individuals sampled (I. bottom), along with experimental and computational pipelines (II). (**B-F**). Cross-tissue single nucleus atlas. Uniform Manifold Approximation and Projection (UMAP) representation of single nucleus profiles (dots) colored by main compartments **(B)**, broad cell types **(C)**, tissues **(D)**, isolation protocol **(E)** and individual donors **(F)**. **(G)** Distinct cell type composition across tissue. Overall proportion of cells (%) of each type (color legend) in each of eight tissues (rows), and number of nuclei profiled in each tissue (right). Numbers in circles: corresponding broad cell type in the legend. Black vertical lines within each colored bar: relative proportion of nuclei from each individual.

We processed the initial snRNA-seq profiles to retain high-quality nuclei profiles and remove the effects of contaminant transcripts from ambient RNA (**Methods**). In breast and skin, the vast majority of the nuclei profiles were recovered from only one individual sample each (breast: 61.3%, skin: 93.1%, **Table S1**). Some tissues and protocols had higher levels of ambient RNA contamination, reflected by detection of spurious expression of highly-expressed transcripts from one cell type in profiles of cells of other types. Such effects were more prominent in skeletal muscle and heart (FDR < 0.05), irrespective of protocol, but were also present in other tissues (**Methods**, Figure S1). We corrected for ambient RNA contamination with CellBender v2.1 (Fleming et al., 2019) (Figure S1) and further applied standard quality control metrics (**Methods**), retaining 209,126 nuclei profiles across the eight tissues, with a mean of 918 genes and 1,519 transcripts (Unique Molecular Identifiers (UMIs) detected per profile.

### Cross-tissue atlas annotation recovers diverse cell types, including difficult-to-profile and rare cell subsets

To facilitate exploration of the entire dataset and guide the identification of cell subsets, including rare types, we integrated data from all samples and methods using a conditional variational autoencoder (cVAE). We designed the cVAE to explicitly correct for multiple sources of variation in expression, such as individual-, sex- and protocol-specific effects, while preserving tissue- and cell type–specific variation (**Methods**, Figure 1B-F, Figures S2A and S3). Cells grouped first by cell type, and then by tissue-specific sub-clusters (Figure 1B-D), suggesting that the variation between cell types is larger than the variation within a cell type across tissues. We comprehensively annotated cell types within each tissue after dimensionality reduction and graph-based clustering (**Methods**) by identifying genes differentially expressed between clusters and comparing them with known literature-based markers (**Methods**, **tables S2 and S3**). Through multiple iterations, we hierarchically defined cellular compartments shared across tissues (*e.g.*, adipose, endothelial, epithelial, fibroblast, immune, muscle) (Figure 1B), broad cell types (*e.g.*, luminal epithelial cells, vascular endothelial cells) (Figure 1C, Figure S4 **and** S5), and granular cell subsets (*e.g.*, luminal epithelial cell 1 and 2, vascular endothelial cell 1 and 2) (Figure S6). The annotations were robust across extraction protocols, tissues, and donors (Figure S2B,D).

The atlas features 43 broad cell classes (Figure 1C, **tables S2** and **S3**), including both tissue-shared cell types and tissue-specific subsets (*e.g.*, Figure 1G, Figures S2B,D **and** S4). For example, tissue-specific cell types such as pneumocytes (alveolar type I and II), keratinocytes, and luminal epithelial cells were the dominant cell types in the lung, skin, and breast, respectively. Macrophages (analyzed in depth below) comprised the largest immune population, with diverse subsets of tissue-resident cells. Shared cell types such as immune and stromal cells were detected across all tissues (Figure S2D,E).

In particular, the single nucleus atlas captured profiles from cell classes that are difficult-to-profile by dissociation-based scRNA-seq (Kim et al., 2020; W. Sun et al., 2020; Wolfien et al., 2020), including 2,350 adipocytes, 21,607 skeletal myocytes, and 9,619 cardiac myocytes. We detected adipocytes in five of the eight tissue types, most prominently breast tissue (86% of adipocytes; 18% of all breast nuclei profiles; Figure 1G), as well as muscle, heart, esophagus muscularis, and skin. The skeletal muscle and cardiac myocytes we detected included key myocyte subtypes (Litviňuková et al., 2020; Tucker et al., 2020). Cardiac myocytes primarily included the previously distinguished classical myocytes and “cytoplasmic myocytes” with high myoglobin (*MB*) expression and a high ratio of exon- *vs*. intron-mapping reads (**Methods**) (Tucker et al., 2020). In skeletal muscle, we similarly observed classical and “cytoplasmic”-like myocytes, as well as neuromuscular junction-rich skeletal muscle myocytes, as previously reported from sc/snRNA-seq studies in mice (Kim et al., 2020; Petrany et al., 2020; Verma et al., 2021). The high diversity of myocyte nuclei may be attributed to nucleus specialization in multinucleation in muscle syncytia. Furthermore, we distinguished “fast-twitch” and “slow-twitch” states in both myocyte and cytoplasmic myocyte populations (Figure 1B,C, Figure S6G).

Cross-tissue and cross-sample integration enhanced our ability to resolve multiple rare cell subsets (Figure 1C, Figures S2B,D, S5 **and** S6). We identified Schwann cells in multiple tissues by differential expression of *CDH19, CADM1, CADM2*, and *S100B*; Interstitial Cells of Cajal (ICCs, expressing *KIT, ANO1* and *PRKCQ*) and enteric neurons (*NPY, VIP, NOS1, ELAVL4, GAL, (Drokhlyansky et al., 2020)*) in esophagus muscularis; neuro-muscular junction (NMJ)-rich myocytes (*CHRND, ETV5*) and skeletal muscle satellite cells (*PAX7, CALCR, MUSK*) (Petrany et al., 2020); hillock and club epithelial cells in prostate (*KRT13*, *MUC4* and *SCGB1A1*, *SCGB3A1*); neuroendocrine cells (*CHGA, CHGB*) in esophagus mucosa and prostate and cornified keratinocytes (*FLG2, LOR*) in skin (Figures S5 and S6).

### snRNA-seq protocols perform well across tissues and correspond to scRNA-seq

We benchmarked the performance of our snRNA-seq nucleus extraction protocols (Drokhlyansky et al., 2020; Slyper et al., 2020) relative to each other across all 8 profiled tissues and to other snRNA-seq, scRNA-seq and bulk RNA-seq data sets in relevant tissues. For each dataset we compared standard QC metrics per profiled cell/nucleus, captured cell type diversity and cell type proportions.

Among the four tested nucleus isolation protocols (CST, NST, TST and EZ) the EZ protocol displayed lower performance in each of the eight profiled tissues according to multiple quality metrics (Luecken & Theis, 2019) (**Methods**) (Figure 2A, Figure S7), including the lowest total number of nuclei captured, higher levels of ambient RNA (FDR<0.05, Figure S1A,B), and separate grouping of EZ-profiled samples (Figure S2C, **Methods**). The extraction protocols varied in the proportion of nuclei recovered from each cell type (Figure S2B,D and S8A). TST, CST and NST protocols had comparable cell-type diversity as measured by Shannon entropy (Figure 2A, **Methods**), with higher variability in skin, breast, and prostate (p-value = 0.06241, Fligner-Killeen test), whereas the EZ protocol resulted in lower diversity (Figure 2A, linear mixed effects model effect-size=-1.08, p=5*10^-11^).

**Figure 2.**
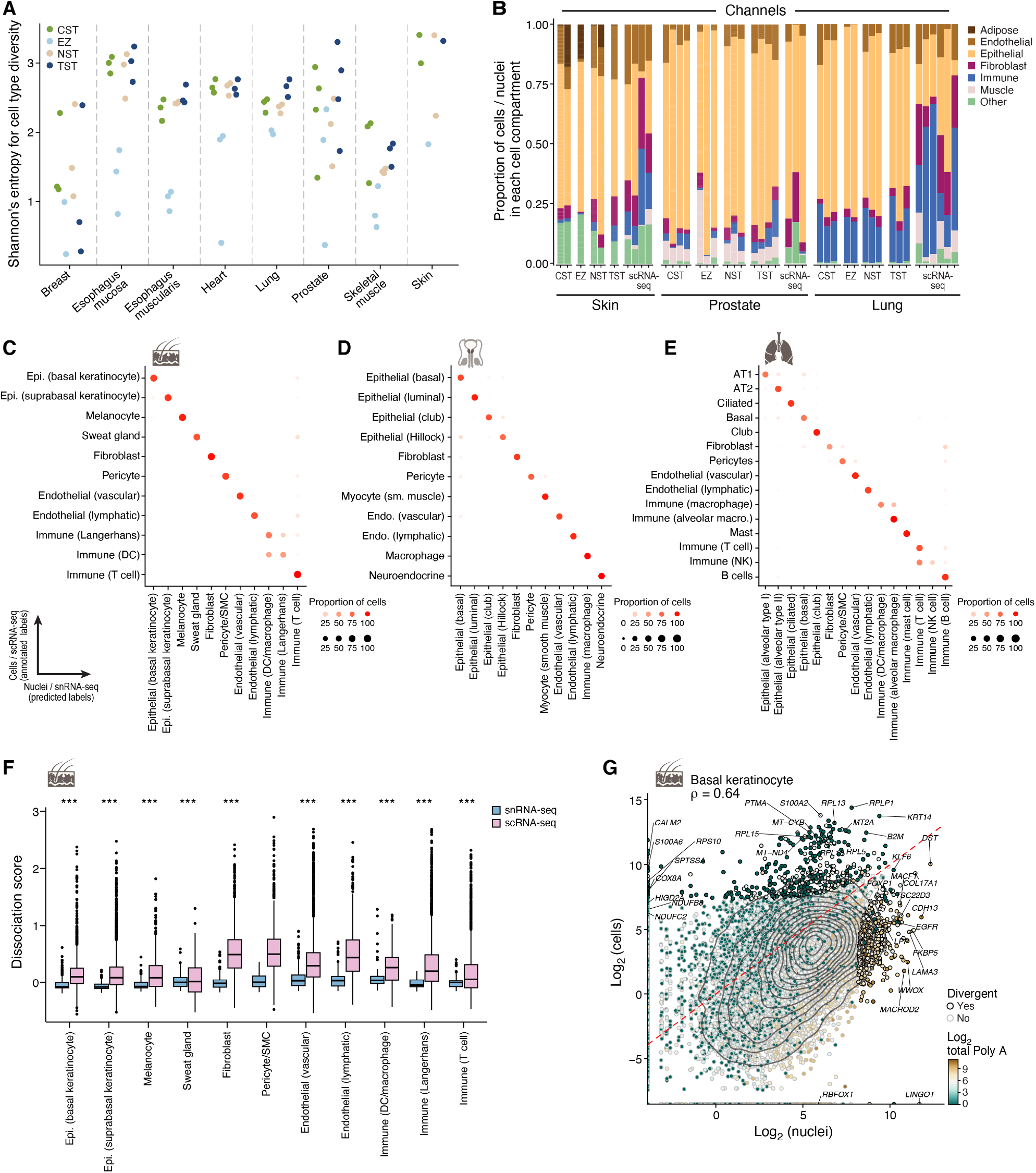
Concordance of cell type diversity and cell intrinsic profiles between snRNA-seq, and scRNA-seq. **(A)** The EZ protocol captures the least cell type diversity. Cell-type diversity (Shannon entropy, *y* axis) of each snRNA-seq dissociation protocol (color legend) in each sample (dot) and tissue (*x* axis). **(B)** Difference in cell proportions captured by snRNA-seq vs. scRNA-Seq. Proportion (% y axis) of cells from major categories (color legend) in each individual, stratified by tissue by each snRNA-seq protocol and by scRNA-seq. **(C-E)** Broad concordance of cell intrinsic programs between scRNA-seq and snRNA-Seq. Proportion of cells (dot color and size) of a test cell class (rows) predicted to belong to a given nucleus class (columns) by a random forest classifier trained on nuclei and applied to cells of the same tissue for skin **(C)**, lung **(D)** or prostate **(E)**. **(G)** Limited number of divergent genes between cell and nuclei profiles are associated with longer polyA stretches. Averaged pseudobulk expression values (**Methods**) of protein-coding genes (dots) in skin basal keratinocytes nuclei (*x* axis) and cells (*y* axis). Divergent genes (black dot outline) deviating from straight line regression fit by size of residuals (**Methods**). Color scale: total length of polyA stretches with at least 20 adenine bases in log_2_ scale. **(F)** Induction of tissue dissociation expression signatures in scRNA-seq but not snRNA-seq profiles. Distribution of the score (y axis, average background corrected log(TP10K+1)) of a dissociation–related stress signature van den Brink et al. [REF] in scRNA-seq (pink) and snRNA-seq (blue) profiles in each major lung cell type (x axis). (*** Benjamini-Hochberg FDR < 10^-16^, Wilcoxon rank-sum test). Box plots show median, quartiles, and whiskers at 1.5 IQR (interquartile range).

Next, we compared cell type compositions recovered in frozen heart left ventricle samples by either our CST, TST, NST and EZ protocols or heart in two public snRNA-seq studies (Litviňuková et al., 2020; Tucker et al., 2020), finding agreement in broad cell types (*e.g.*, mast cells, adipocytes, “cytoplasmic” cardiac myocytes, and Schwann cells (Figure S8B,C **and Table S4**) but differences in some of their proportions (Figure S8D,E). Both the published studies and our EZ snRNA-seq had a higher proportion of muscle cells, whereas CST, NST and TST had a higher proportion of endothelial cells (Figure S8E). There was also high concordance between the expression profiles of bulk RNA-seq (from GTEx, (GTEx Consortium, 2020)) and pseudobulk profiles derived from our snRNA-seq (accuracy 92.3%, Figure S9), with a few samples showing lower agreement (heart-EZ, breast-EZ, and two breast-TST samples).

We also compared snRNA-seq and fresh-tissue scRNA-seq of lung (MS, ORR, AR, Avinash Whagry, Alexander Tsankov, Jay Rajagopal et al., unpublished results), skin (current study, **Methods**), and prostate (Henry et al., 2018). For cell composition (Figure 2B), we recovered the same main cell groups across compartments, as also reflected by the accuracy of a multiclass random forest classifier trained on scRNA-seq data in predicting cell types on snRNA-Seq data (**Methods**, Figure 2C-E) and vice versa (Figure S8F-H), and in overall similarity of cell type intrinsic (pseudobulk) profiles of protein coding genes (average Spearman ρ across cell types = 0.58 (skin), 0.69 (prostate), 0.53 (lung), **Table S4**). Notable divergences include the greater expression in cells vs. nuclei profiles of a stress/dissociation signature (Denisenko et al., 2020; van den Brink et al., 2017) (Wilcoxon rank-sum test, Benjamini-Hochberg FDR < 10^-16^, Figure 2F, Figure S10A,B), as we previously reported (Slyper et al., 2020), and of ribosomal and nuclear-encoded mitochondrial protein genes (Figure 2G, **Methods**, linear model), consistent with their longer half-lives and higher cytoplasmic levels (Rabani et al., 2014; Zaghlool et al., 2021). Conversely, nuclei had higher levels of longer transcripts (Figure S10H,I) and of transcripts with higher counts of adenine stretches (Figure 2G, Figure S10C-G), consistent with previous reports (Solnestam et al., 2012). Notably, snRNA-seq generally captured relatively lower proportions of lymphocytes overall and as a component of the immune compartment. For example, for lung and skin, respectively, T cells represented 1.7% or 1.4% of all cells and 9% or 29.5% of the immune compartment by snRNA-seq, compared to 18.4% or 6.83% overall and 56.8% or 43% of the immune compartment by scRNA-seq (with similar patterns for B cells). Proportions varied across samples and protocols. Note that a recent study comparing snRNA-seq and *in situ* measurements (Hwang et al., 2020) suggested that scRNA-seq may over-sample immune cells.

### A dichotomy between *LYVE1*- and *HLA* class II-expressing macrophages preserved across tissues

Our cross-tissue atlas allowed us to characterize tissue-specific and shared features of resident immune cells, focusing on macrophages. Tissue-resident immune cells play key roles in immune surveillance and tissue support and are shaped by both their ontogeny and tissue residence (Chakarov et al., 2019).

Integration and annotation of 14,156 myeloid nuclei profiles (**Methods**) (60% of all immune nuclei) revealed distinct monocyte, macrophage and DC populations (Figure 3A, and Figure S11A,B). These included CD16^+^ monocytes (*FCGR3A/CD16, LILRB1, LILRB2*), CD14^+^ monocytes (*VCAN, S100A8, FAM65B*), two transitional Mo/MΦ FCGR3A^lo^ and Mo/MΦ FCGR3A^hi^ populations with co-expression of both monocyte and macrophage markers, DC1s (*C1orf86, CLEC9A, XCR1*), DC2s (*CD1C, CLEC10A*) (Villani et al., 2017), mature DCs (*CCR7, LAMP3, BIRC3*) and Langerhans cells (*CD207/Langerin, CD1A, RUNX3*) (Figure 3B). Tissue macrophage states included lung macrophages (*PPARG, SLC11A1, MARCO*) expressing a reported alveolar macrophage signature (Reyfman et al., 2019) (Figure S11C), proliferating macrophages (*TOP2A, MKI67, UBE2C*), cytokine/chemokine-expressing inflammatory macrophages (*IL1B, CCL4L2, CCL4*) and two additional macrophage subsets: MΦ *LYVE1*^hi^ (*F13A1, SEPP1, LYVE1*) and MΦ *HLA*II^hi^ (*APOE*, *APOC1*, *HLA-DRB1*) (Figure 3B). Finally, we identified lipid-associated macrophage (LAM)-like nuclei profiles, with high expression of LAM-specific genes (*SPP1, FABP5, CD9, TREM2*) (Figure 3B and Figure S11C) (Jaitin et al., 2019) and of lipid metabolism-related expression modules, including “regulation of lipid localization” and “response to lipoprotein particle” (Figure S11D).

**Figure 3.**
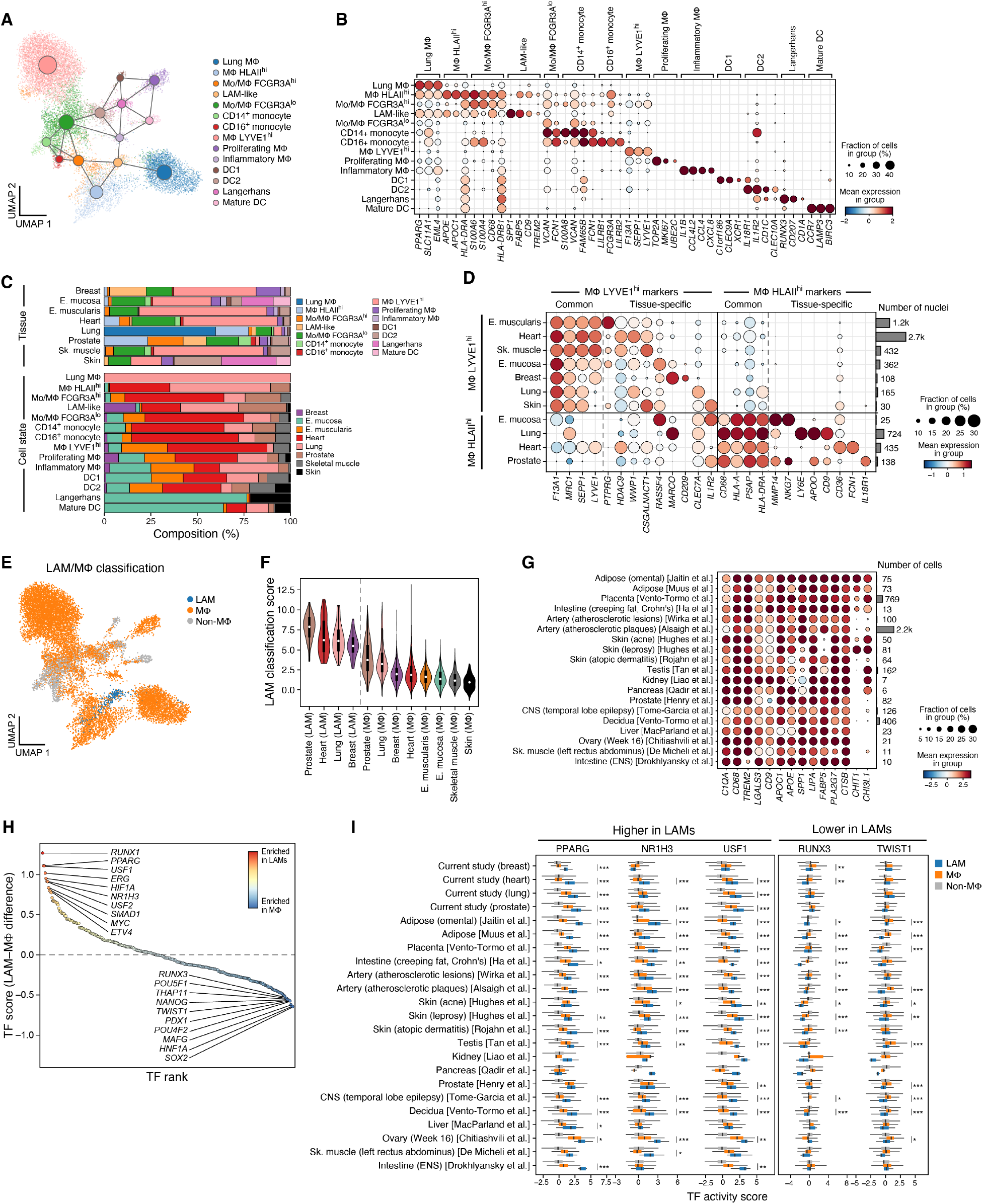
Cross-tissue analysis of myeloid cells highlights a dichotomy between LYVE1- and HLA class II-expressing macrophages and broad presence of LAM-like populations. **(A-C)** Myeloid cell states in the cross-tissue atlas. **(A)** UMAP visualization of snRNA-seq profiles of myeloid cells (dots) in the cross-tissue atlas, colored by cell type/state and overlaid with a PAGA graph of myeloid states (large nodes). **(B)** Mean expression (circle color) and fraction of expressing cells (circle size) of marker genes (columns, labels at bottom) associated with each myeloid subset (rows, and labels on top). **(C)** Myeloid cell distribution across tissues. Top: Overall proportion of myeloid cells of each subset (%, colors) in each tissue (bars). Bottom: Overall proportion of cells from each tissue (%, colors) for each myeloid subset (bars). **(D)** Cross-tissue and tissue-specific markers of MΦ *LYVE1*^hi^ and MΦ *HLA*II^hi^ states. Mean expression (circle color) and fraction of expressing cells (circle size) of marker genes (columns, labels at bottom) associated with the two myeloid subsets (labels on left) in each tissue (rows). Common (cross-tissue) and tissue-specific markers are labeled on top. Right bar plot: number of nuclei for each subset in each tissue. **(E-G)** LAM-like cells across tissues. **(E)** UMAP visualization of myeloid cells (dots) colored by their classification as LAM-like, other macrophages (Mɸ) and non-macrophages (non-Mɸ) using a linear classifier (color legend, **Methods**). **(F)** Distributions of classification scores (y axis) of LAM-like and other macrophages across tissues (x axis). **(G)** Mean expression (circle color) and fraction of expressing cells (circle size) of LAM marker genes (columns) in cell or nuclei profiles from other studies (rows) classified as LAM-like by the classifier. **(H,I)** *PPARG*, *NR1H3*, *USF1* are inferred as TFs regulating the LAM-like program. (**H**) TF differential activity score between LAMs and other macrophages (y axis) for each TF (dot) ranked by their score (x axis). **(I)** Distribution of TF differential activity scores (x axis) for three TFs with significantly high (two tailed *t* test, *:Benjamini-Hochberg FDR<0.05, **:FDR<0.01, ***:FDR<0.001) scores in LAMs (*PPARG*, *NR1H3*, *USF1*) or other macrophages (*RUNX3*, *TWIST1*) in LAM-like, other macrophages (Mɸ) and non-macrophages (non-Mɸ) myeloid cells (color legend) in each study (y axis). Box plots show median, quartiles, and whiskers at 1.5 IQR (interquartile range).

Most myeloid subsets were generally present in multiple tissues, with the notable exceptions of lung macrophages and Langerhans cells. Myeloid cell proportions were more highly correlated between samples within a tissue type (Figure S11E,F) than between different tissues (Figure S11F,G), suggesting that tissues have characteristic myeloid state proportions. Moreover, related tissues – such as muscle (heart, esophagus muscularis, skeletal muscle) or mucosa (esophagus mucosa, skin) – grouped by their myeloid composition profiles (Figure S11G). Breast, esophagus mucosa, esophagus muscularis, heart, and skeletal muscle had significantly higher proportions of MΦ *LYVE1*^hi^ (p-value<10^-6^, Dirichlet regression LRT, **Methods**), while lung and prostate had significantly higher proportions of MΦ *HLA*II^hi^ (p-value<10^-8^, Dirichlet regression LRT, **Methods**) (Figure 3C, and Figure S11E,H,I). In contrast to other subsets, lung macrophages were present only in lung, and Langerhans cells only in skin and esophagus mucosa (97%), consistent with their role in antigen sampling within stratified epithelia (Capucha et al., 2015; Deckers et al., 2018) (Figure 3C, and Figure S11H). Interestingly, while LAMs have first been reported in human visceral adipose tissue (Jaitin et al., 2019) and multiple pathological contexts (Deczkowska et al., 2020), LAM-like cells were more widely distributed across healthy human tissues in our atlas, predominantly breast, heart, lung and prostate (Figure 3C, 94%, 268 of 283), as we discuss below.

Analysis of MΦ *LYVE1*^hi^ and MΦ *HLA*II^hi^ populations across esophagus mucosa, lung and heart revealed that a dichotomy between *LYVE1*^high^*HLA* class II^low^ and *LYVE1*^low^*HLA* class II^high^ states was preserved across the tissues. Each subset was characterized by a combination of a common, tissue-agnostic signature (MΦ *LYVE1*^hi^: *F13A1, MRC1, LYVE1*, *SEPP1* and MΦ *HLA*II^hi^: *CD68, HLA-A, PSAP, HLA-DRA*) along with tissue-specific markers of MΦ *LYVE1* hi or MΦ *HLA*II^hi^ states (Figure 3D, and Figure S11J), such as *MARCO* and *CD209* in breast MΦ *LYVE1*^hi^, *PTPRG* in esophagus muscularis MΦ *LYVE1*^hi^, *LY6E* in lung MΦ *HLA*II^hi^, *IL18R1* in prostate MΦ *HLA*II^hi^, and *NKG7* in esophagus mucosa and prostate MΦ *HLA*II^hi^ (all FDR<0.05, Welch’s t-test). The dichotomous *LYVE1*^high^*HLA* class II^low^ and *LYVE1*^low^*HLA* class II^high^ states were enriched for distinct functions: tissue supporting modules (*e.g.*, “neurogenesis” and “endothelial cell fate specification”) for MΦ *LYVE1*^hi^ and immune-related processes (*e.g.*, “myeloid leukocyte mediated immunity”, “interferon-mediated signaling pathway”, “antigen-processing and presentation”) for MΦ *HLA*II^hi^ (Figure S11K), and were remarkably similar to *Lyve1*^high^MHC II^low^ and *Lyve1^low^*MHC II^high^ resident macrophage populations in mouse tissues (Chakarov et al., 2019) (Figure S11L).

### LAM-like macrophages are prevalent across human tissues in health and disease and are associated with a shared regulatory program

LAMs and LAM-like cells have been observed in disease contexts in adipose tissue from obese human individuals and from mice (Jaitin et al., 2019), injured and fibrotic liver (Perugorria et al., 2019; Ramachandran et al., 2019; Xiong et al., 2019), fibrotic lung (Ayaub et al., 2021; Reyfman et al., 2019), atherosclerotic aortic tissue (Cochain et al., 2018; Willemsen & de Winther, 2020), and Alzheimer’s disease brain (Grubman et al., 2019; Keren-Shaul et al., 2017; Mathys et al., 2019). However, an understanding of their distribution and heterogeneity across human tissues is still lacking.

We systematically characterized LAM-like cells in the expanded context of our study and 17 other atlases from 14 tissues, by training a linear classifier with published omental scRNA-seq containing LAMs (Jaitin et al., 2019) (**Methods**) and classifying each myeloid profile in our dataset and the published compendium as LAM-like macrophages, non-LAM macrophages and non-macrophages, recovering 283 LAM-like cells in our study and 4,285 LAM-like cells in 17 reported studies (Figure 3E-G, and Figure S12A-D, **Table S5**).

The LAM-like cells were present among tissue-resident macrophages across a broad range of tissues and pathologies. These included adipose (Jaitin et al., 2019; Muus et al., 2021) and atherosclerotic (Alsaigh et al., 2020; Wirka et al., 2019) tissue, as previously reported, as well as many healthy tissues (placenta, testis, kidney, pancreas, prostate, decidua, liver, ovary, skeletal muscle, intestine) and other disease contexts (skin from acne, leprosy and atopic dermatitis patients, ileum from Crohn’s disease patients). Notably, 126 microglia (high *TMEM119* expression) from the CNS of epileptic patients were also classified as LAM-like cells, indicating that LAM-signature expressing microglia extend beyond Alzheimer’s disease (Grubman et al., 2019; Keren-Shaul et al., 2017; Mathys et al., 2019). A core signature of LAM genes, including *FABP5, SPP1, CSTB, APOC1* and *CD9*, is consistent across most tissues and studies, whereas *CHIT1* and *CHI3L1* showed considerable variation (Figure 3G and Figure S12E).

We predicted transcription factors (TFs) that could mediate LAM-like gene expression, by inferring TF activities in LAMs, non-LAM macrophages and all other cells separately, and ranking TFs by the mean difference between their activities in LAMs *vs*. non-LAM macrophages (**Methods**, Figure 3H). LAM-associated TFs included *PPARG, USF1* and *NR1H3* (*LXRA*) across all classified LAM-like cells (Figure 3I, Figure S12F), consistent with a shared core regulatory mechanism. These are major regulators of lipid metabolism-related gene expression (Kidani & Bensinger, 2012) and have been proposed to regulate *Trem2* expression in mice (Savage et al., 2015).

### Genes from monogenic muscle disease groups are enriched in distinct subsets of myocyte and non-myocyte cells in cardiac, skeletal and smooth muscle tissue

While human genetics has successfully identified many rare monogenic disease genes, the cell type(s) of action are often unknown, or can even be incorrect, as shown for *CFTR*, the gene underlying cystic fibrosis (Montoro et al., 2018; Plasschaert et al., 2018). Leveraging the three muscle types in our atlas—cardiac, skeletal muscle, and smooth muscle—we sought to identify the cell type(s) of action for a broad group of well-curated monogenic muscle disease genes (Benarroch et al., 2020) (**Table S6**). We tested the subset of genes for each disease group (*e.g.*, hereditary cardiomyopathies, motor neuron diseases) for enrichment in cell type-specific markers from our muscle tissues (FDR < 0.1) (**Methods**, Figure 4A).

**Figure 4.**
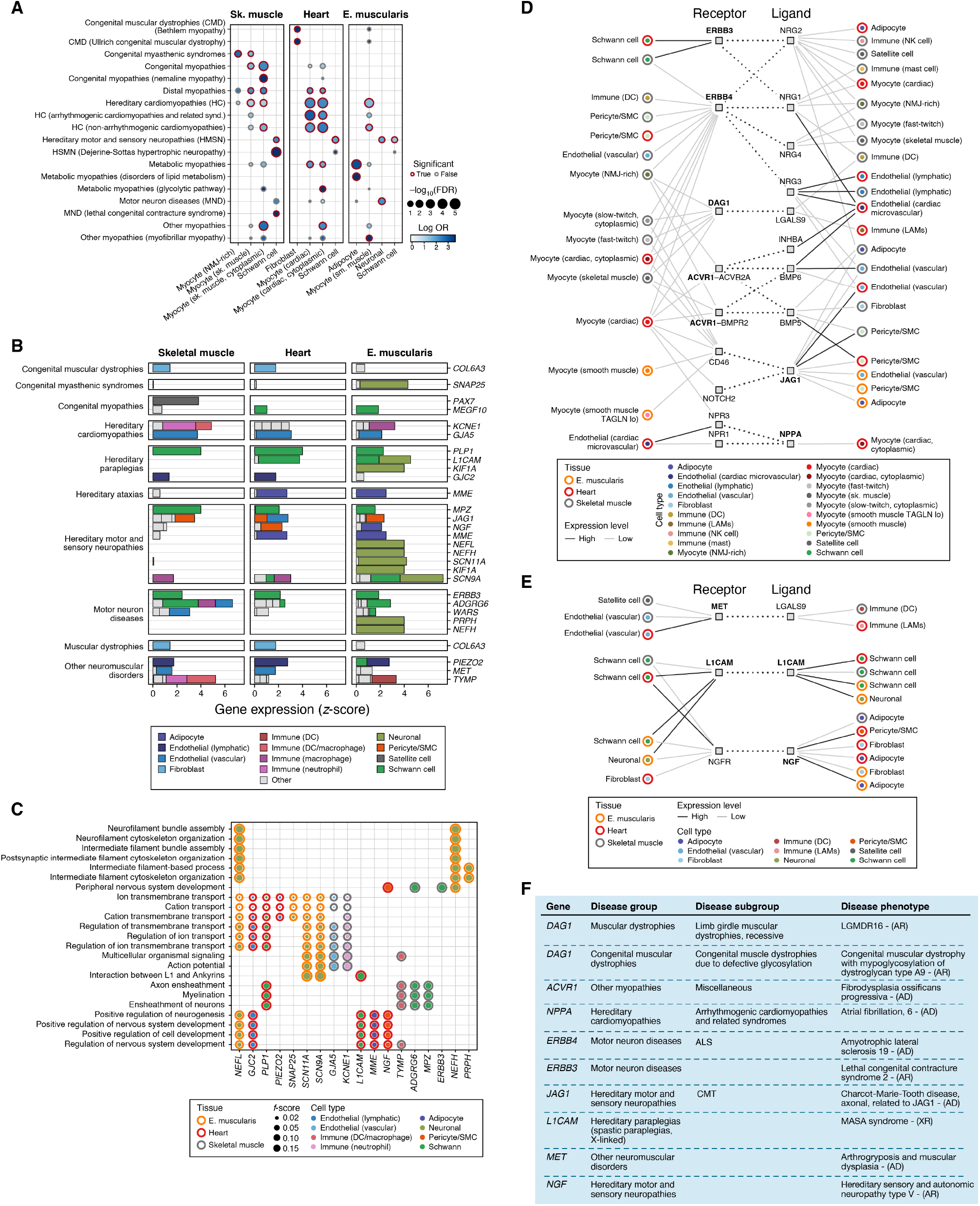
Monogenic muscle disease genes are enriched in subsets of myocytes and non-myocyte cells and their interactions in cardiac, skeletal and smooth muscle tissue. **(A)** Relation of broad cell types to monogenic muscle disease groups. Effect size (log odds ratio, dot color) and significance (-log_10_(Benjamini-Hochberg FDR), dot size) of enrichment of genes from each monogenic muscle disease group (rows) for cell type markers of broad cell subsets in each tissue (columns). Red border: Benjamini-Hochberg FDR<0.1. **(B)** Monogenic muscle disease genes expressed in non-myocytes in three muscle tissues. Expression (z score, x axis) of monogenic muscle disease genes (y axis) ordered by disease category (labels on left) in each cell type (color legend) in each of three tissues (labels on top). Grey (“other”): Cell types with low expression of the indicated genes. **(C)** Biological processes enriched in monogenic muscle disease genes expressed in non-myocytes. F-score (circle size, harmonic mean of precision and recall) of the degree of overlap between a functional gene set (rows; GO biological process and Reactome, **Methods**) and the muscle disease genes in (B) that are expressed in non-myocytes. An entry is shown if the functional gene set is enriched (FDR<0.1, Fisher’s exact test) and the gene is a member of the functional set. **(D,E)** Putative cell-cell interactions in muscle implicating monogenic muscle disease genes. Cell types (circle nodes, inner dot color) from muscle tissues (circle nodes, outer circle color) connected by putative interactions (dotted edges) between a receptor (left square nodes) expressed in one cell type and a ligand (right square nodes) expressed in the other (solid edges; black: cell type with high expression), where either the receptor or the ligand is a monogenic muscle disease gene (bold), shown for interactions involving myocytes **(D)** or only non-myocyte **(E)** cells. **(F)** Diseases and disease groups of monogenic disease genes highlighted in the cell-cell interactions in (D,E). **Acronyms** ALS: Amyotrophic lateral sclerosis; CMT: Charcot-Marie-Tooth disorder; AR: Autosomal recessive; AD: Autosomal dominant; NMJ: Neuromuscular junction.

Different disease group gene subsets were associated not only with different myocyte subsets (spanning 113 of 606 genes; Figure S14), but also with neurons, Schwann cells, fibroblasts, and adipocytes (127 genes, Figure S15, **Table S7**), in patterns that recapitulated known disease mechanisms as well as highlighted new relationships (Figure 4A). Known associations in myocytes included skeletal muscle myocytes with congenital myopathy genes (FDR: 7.07*10^-5^, *e.g.*, *ACTA1, BIN1, MYH7, MYL1, TPM2, TTN*), and cardiac myocytes with hereditary cardiomyopathy genes (FDR: 4.24*10^-12^, *e.g.*, *ACTC1, ACTN2, CACNA1C, CACNB2, MYH6, MYH7, MYL4, MYOZ2, SCN2B, SCN5A*). In some cases, associations highlighted finer myocyte subsets, such as congenital myasthenic syndrome (CMS) genes with NMJ-rich skeletal myocytes (FDR: 0.015, e.g., *CHRNA1, CHRND, CHRNE, MUSK*). Notably, *CHRNE* and *MUSK* are specific to NMJ-rich myocytes but not to other skeletal myocytes (**Table S7**), highlighting the importance of finer subsets. Among non-myocyte cell associations were Schwann cells with hereditary motor and sensory neuropathies in all three tissues (FDR: 0.015-0.06; *DST, EGR2, MPZ, NDRG1, PMP22, PRX*) and with Dejerine-Sottas hypertrophic neuropathy in skeletal muscle (FDR: 2.28*10^-5^, *e.g.*, *EGR2, MPZ, PMP22, PRX*); enteric neurons in the esophagus muscularis with hereditary motor and sensory neuropathies (FDR: 0.034, *e.g.*, *KIF1A, KIF1B, NEFH, NEFL, PMP22, SCN11A*) and with motor neuron diseases (FDR: 0.01, *ERBB4, KIF26B, MAPT, NEFH, NEK1*), and esophagus muscularis adipocytes with metabolic myopathies related to lipid metabolism (FDR: 0.0003, *LPIN1, PNPLA2, SLC22A5, SLC25A20*). Notably, some of the enrichments in non-myocytes are also present in the same cell types in other tissues (*e.g.*, breast adipocytes for metabolic myopathies or breast and skin pericytes for cardiomyopathies, Figure S13I). This reflects how a specific pathology may arise from the relation between an accessory cell’s broader function and specialized tissue demand.

Monogenic muscle disease genes that are significantly highly expressed in non-myocytes but not detected in myocytes (Figure 4B, **Table S8**) are enriched for peripheral nervous system development, myelination, ensheathment of neurons, regulation of ion transmembrane transport, interaction between L1 and Ankyrins and neurofilament bundle assembly genes (Figure 4C). Key examples include *PAX7* (congenital myopathy related to PAX7) in satellite cells, *TYMP* (mitochondrial DNA depletion syndrome), and *KCNE1* (long QT syndrome). Although the cell type of action might be functioning in the CNS for some of these diseases, our analysis identified these potential neuronal components of the pathology through enteric neurons, suggesting the commonality between neuron identity signatures of the peripheral and central nervous systems.

We observed similar enrichment patterns when analyzing the orthologous genes in snRNA-seq data from the same muscle tissues in mice (Drokhlyansky et al., 2020) (**Methods**, Figure S13). In particular, we observed significant associations between skeletal muscle myocytes and various dystrophies and myopathies (FDR<0.1, Fisher’s exact test), cardiomyocytes and cardiomyopathies (FDR<0.1), adipocytes in skeletal esophagus and heart and metabolic myopathies (FDR<0.1), Schwann cells in skeletal muscle and hereditary motor and sensory neuropathies (FDR<0.1) (Figure S13H).

### Some monogenic muscle disease genes may impact cell interactions in the tissue

To better understand the potential impact of monogenic muscle disease genes on the interplay between cell types in the tissue, we related cells in each of the three muscle tissues by receptor-ligand interactions, where at least one of the genes was a monogenic disease gene (**Methods**). Interactions involving myocytes were mediated by the disease genes *DAG1* (congenital muscular dystrophy), *ACVR1* (fibrodysplasia ossificans progressiva), *NPPA* (atrial fibrillation), *JAG1* (Charcot-Marie-Tooth disease), *ERBB3* (lethal congenital contracture syndrome) and *ERBB4* (Charcot-Marie-Tooth disease), with implications for how myocyte interactions with other cell-types might be disrupted in these diseases. *(Becker et al., 2014; C. Wu et al., 2018)* (Figure 4D, **Table S9**). For example, there is a putative Schwann-myocyte interaction, mediated by Schwann cell–specific expression of the disease gene *ERBB3* and its ligands *NRG1* and *NRG2* in myocytes, as well as by *ERBB4* (with a broader expression pattern). Putative cell-cell interactions involving only non-myocytes included the disease genes *L1CAM* (MASA syndrome), *MET* (arthrogryposis and muscular dysplasia) and *NGF* (hereditary sensory and autonomic neuropathy), each potentially affecting multiple cell pairs, including neurons and Schwann, satellite, immune and stromal cells (Figure 4E,F, **Table S9**).

### Cell type–specific enrichment of QTL genes mapped to GWAS loci

Studies associating genetic variants to changes in expression (eQTLs) or splicing (sQTLs) traits have demonstrated tissue-specific colocalization with multiple loci from GWAS of human traits, including disease risk (Barbeira et al., 2021; Finucane et al., 2015; Gamazon et al., 2018; GTEx Consortium, 2020), but lacked cellular resolution (Kim-Hellmuth et al., 2020). To relate e/sQTLs to cell types and prioritize causal genes for complex diseases and traits in specific cells and tissues, we tested if GWAS loci from 21 complex traits (**Table S10**), with likely effects in at least one of the 8 tissues analyzed, are enriched for genes with high cell type–specific expression in each tissue. We defined putative causal genes for each GWAS locus as the set of genes whose e/sQTLs(GTEx Consortium, 2020) were in LD (r^2^>0.8) with the lead GWAS variant(s) (Figure 5A and **Methods**). We further included genes prioritized by additional genomic data (*e.g.*, Hi-C) and linkage to predicted deleterious protein coding variants (Ghoussaini et al., 2021). Since more than one gene typically maps to a GWAS locus using this approach (mean=2, max = 37 for selected traits and 170 for null traits), we scored loci by the fraction of cell type-specific genes in the locus (Figure 5A and **Methods**). We assessed the enrichment of loci with cell type–specific genes above the 95^th^ percentile against a null distribution of thousands of LD-clumped GWAS loci from hundreds of complex traits from Open Targets Genetics (Ghoussaini et al., 2021), using a Fisher’s exact test-based approach (Figure 5A and **Methods**).

**Figure 5.**
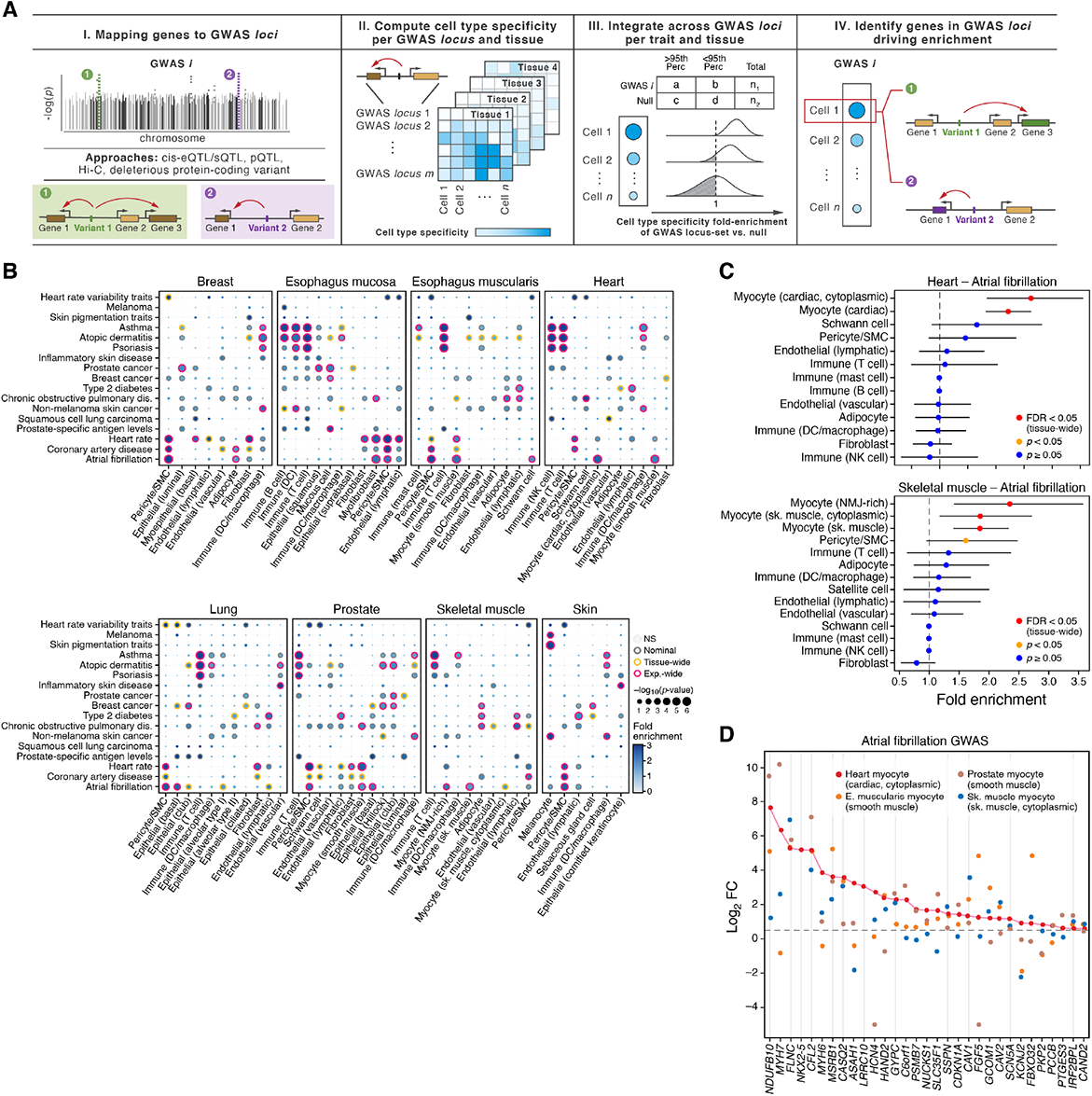
Cell type-specific enrichment of expression and splicing QTL-mapped genes to GWAS loci in 17 diseases and traits relevant to the 8 atlas tissues. **(A)** Schematic of the GWAS cell type specificity enrichment method. **(B)** Cell type enrichment of genes mapped to GWAS loci for 17 of the 21 complex traits tested with at least one tissue-wide significant result (Benjamini Hochberg (BH) FDR<0.05 correcting for all cell types tested per tissue per trait) across 8 GTEx tissues. Significance (circle size, −log_10_(P-value)) and effect size (circle color, fold-enrichment) of enrichment of GWAS locus sets (rows) in each broad cell type category (columns) in the eight tissues in the cross-tissue atlas (panels). Grey, orange, red borders: nominal, tissue-wide, and experiment-wide (BH FDR<0.05 correcting for all cell types tested across 8 tissues and 21 traits) significance results. Cell types with no tissue-wide significant enrichment were omitted. **(C,D)** Myocyte and pericyte genes enriched in atrial fibrillation GWAS loci (P<0.05). (**C**) Fold-enrichment (*x* axis) of the cell types (*y* axis) for atrial fibrillation GWAS in heart (top) and skeletal muscle (bottom). Error bars: 95% credible intervals. Red: tissue-wide significant; Orange: nominal significance; Blue: non-significant (P≥0.05). **(D)** Differential expression (log_2_(Fold-change), y axis) in myocytes compared to all other cell types from heart (red, cardiac myocytes), skeletal muscle (blue, skeletal muscle cytoplasmic myocytes), esophagus muscularis (orange, smooth muscle) and prostate (brown, smooth muscle) of the genes (x axis) driving the enrichment signal of the atrial fibrillation GWAS loci in heart cardiac myocytes. Gray vertical lines: genes with log_2_(fold-change) >0.5 and FDR<0.1 in myocytes in all four tissues.

Seventeen (17) of the traits were enriched in both expected and previously undescribed cell types at tissue-wide FDR<0.05 (Benjamini-Hochberg), 16 of which were also significant (FDR<0.05) across tissues (Figure 5B, **Tables S11** and **S12**, Figures S16 and S17). These included expected associations such as skin pigmentation traits in melanocytes, autoimmune and inflammatory diseases in T and NK cells, lung fibroblasts in COPD, prostate cancer in luminal epithelial cells, atrial fibrillation and heart rate in myocytes, heart rate in lymphatic endothelial cells (Brakenhielm & Alitalo, 2019), and coronary artery disease in vascular endothelial cells (Figure 5B). Interestingly, type 2 diabetes loci were strongly enriched in skeletal muscle adipocytes as well as in lymphatic endothelial cells in multiple tissues, which might underpin the predisposition of type 2 diabetes to vascular disease (Figure 5B). Less studied examples include DCs in non-melanoma skin cancer, adipocytes and breast cancer (Kothari et al., 2020), and adipocytes and atrial fibrillation. In many cases, when GWAS loci were enriched in a specific cell type from a known tissue of action, a similar enrichment was observed in the same cell type from other uninvolved tissues. For example, atrial fibrillation GWAS loci were enriched in myocytes in heart, skeletal muscle, esophagus muscularis and prostate (Figure 5B,C), coronary artery disease and heart rate loci were enriched in pericytes in 5-6 other tissues in addition to heart, and prostate cancer loci were enriched in luminal epithelial cells in both prostate and breast (Figure 5B, Figures S18 and S19).

Cell type enrichment enables prioritization of specific causal genes in GWAS loci with multiple LD-mapped genes and cell types, such as *NDUFB10*, *MYH7*, *FLNC*, *CFL2*, *MYH6*, *MSRB1*, and *CASQ22* for atrial fibrillation and heart myocytes (cardiac, cytoplasmic) (Figure 5D, **Tables S11** and **S12**). Only a subset of the genes driving the cell type enrichment for different traits (mean = 66%, 60-71.4% [95% CI]) were common across tissues (*e.g.*, *NDUFB10, FLNC, CFL2, MSRB1*, and *CASQ22* for atrial fibrillation and myocytes; Figure 5D), which may point to a high confidence set of causal genes in the particular cell type and suggest additional tissue-specific genes in specific cell types (*e.g.*, the cardiac myosin subunits *MYH6* and *MYH7* in heart myocytes).

### Associating GWAS genes with gene programs across cell types reveals six main trait groups

To chart the cellular programs and processes that may be impacted by genetic variants, we next associated a larger set of >2,000 complex phenotypes with cell types and the co-varying gene modules they express (**Methods**). We defined gene modules in the cells in our atlas by correlation-based gene clustering across cells, scored modules for their overlap with GWAS genes (defined by variant to gene mapping in Open Target Genetics (Ghoussaini et al., 2021)) that are also highly expressed by cell types (Figure S20, **Table S13**), and then grouped GWAS phenotypes into major groups by the similarity of their module enrichment across cell types (Figure 6A-C, Figure S21). For each major group of traits, we identified the relevant cell types associated with the underlying modules (Figure 6A,C) and tested the GWAS genes that overlapped with gene modules for functional enrichments (Figure 6A,D,E).

**Figure 6.**
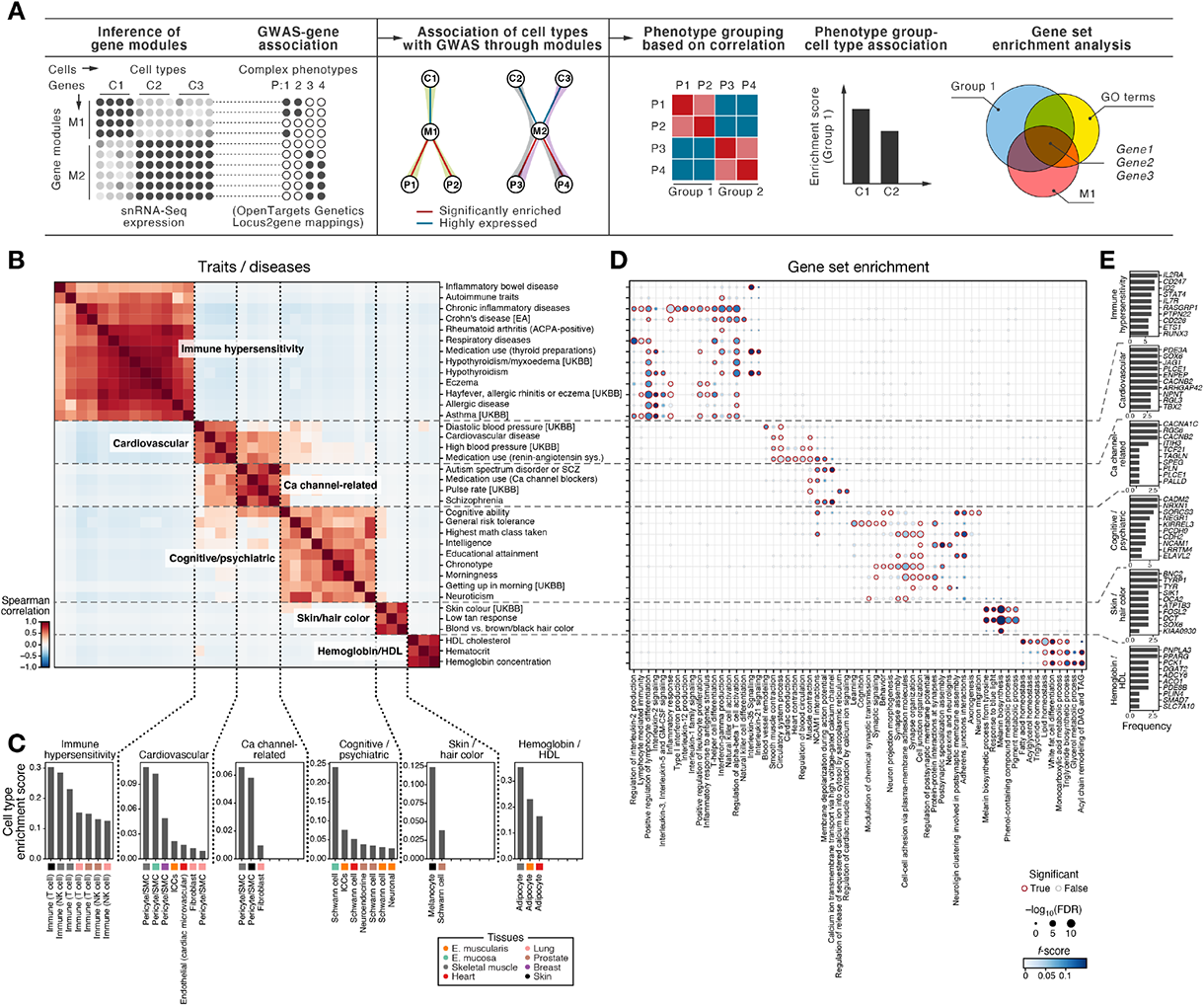
GWAS module enrichment suggests cell types and gene modules relevant for trait groups. **(A)** Schematic of the GWAS gene module enrichment analysis. Shaded edges in the middle panel represent associations between cell types and phenotypes. **(B-E)** Six disease groups identified by GWAS-cell type relationships. (**B**) Similarity (Spearman correlation coefficient, color bar) between GWAS traits/diseases (rows, columns) based on their enriched cell types. Six GWAS trait/disease modules are labeled and marked by dashed lines. **(C)** Cell type enrichment scores (y axis) for cell types (x axis) for each of the GWAS-enriched modules. **(D)** Significance (-log_10_(FDR (Benjamini-Hochberg), circle size) and F score (circle color) of enrichment of functional gene sets (columns) with the genes in the intersection of a gene module and GWAS genes for each trait/disease in (B) (rows). Red border: significant enrichment, Benjamini-Hochberg FDR<0.1. **(E)** Number of traits (x axis) in each module in (B) where a gene (y axis) is detected as the driver of the association between the genes in the 3-way intersection of gene modules, a trait enriched in the module, and a functional gene set, for the top 10 most frequently identified genes in the enrichment analysis of each module. **Acronyms** UKBB: UK Biobank, EA: East Asian, SCZ: Schizophrenia.

We partitioned 6 major groups of module-cell type-trait associations, spanning immune hypersensitivity, cardiovascular, calcium channel–related, cognitive/psychiatric, pigmentation, and high-density lipoprotein (HDL) cholesterol related traits and/or disease (Figure 6B-E). The immune hypersensitivity disorders included asthma, inflammatory bowel disease, Crohn’s disease, and rheumatoid arthritis, as well as three hypothyroidism traits known for autoimmune links (Stassi & De Maria, 2002). This group was associated with T cells, including the expected relation between lung T cells and hay fever, allergic rhinitis, and respiratory disease (Galli et al., 2008; Kay, 2001). GWAS genes in the modules associated with these traits were enriched for lymphocyte activation and differentiation and TCR signaling and cell-cell adhesion (*e.g., PTPRC*, *IL2RA, IL7R, IFNG, FASLG*, Figure 6E), with IL-35 signaling genes enriched in hypothyroidism and IBD, consistent with IL-35 upregulation in Hashimoto’s thyroiditis (Yilmaz et al., 2016) and IBD (Y. Li et al., 2014)) and downregulation in Graves’ disease (Saeed et al., 2021). The cardiovascular traits group included hypertension, atrial fibrillation, mean arterial pressure and renin-angiotensin system, was associated with pericytes/smooth muscle cells (SMCs, responsible for modulating total peripheral resistance in arterioles), and was enriched with blood circulation, smooth muscle contraction, muscle structure development and cardiocyte differentiation genes (*e.g.*, *CACNA1C, CACNA1D, CASQ2, PLN*). Notably, the cardiovascular group overlapped with the calcium-channel related group, which also included blood pressure medication, pulse rate, medication use of calcium channel blockers and vascular system traits, as well as schizophrenia (SCZ) and autism spectrum disorder (ASD), psychiatric disorders with known L-type calcium channels associations (Pinggera et al., 2017), and was enriched with membrane depolarization and calcium ion channel genes (*e.g.*, *CACNA1C, CACNA1D*). Other cognitive/psychiatric phenotypes (*e.g.*, educational attainment, intelligence, depression, neuroticism) grouped separately, and were enriched with neuronal synapse organization, structure, and activity genes (*e.g.*, *NRXN1, CDH2, CNTN4*). A skin/hair color group was driven by melanocytes and enriched with pigmentation and melanin-related processes (*TYR, OCA2, TYRP1, MLANA*). Finally, a group of HDL cholesterol traits was associated with adipocytes in all three muscle tissues, and enriched for fatty acid, triglyceride, and lipid homeostasis and related metabolic processes (monocarboxylic and glycerol metabolism), including *ANGPTL8* and *PNPLA3*, genes involved in lipolysis regulation in adipocytes that might impact extrahepatic cholesterol transport via HDLs (Mysore et al., 2017; Yang & Mottillo, 2020).

### Toward large-scale snRNA-seq of human tissues with pooling

To enable future studies at population scale, we tested whether frozen samples from different individuals can be pooled for snRNA-seq, followed by computational demultiplexing, as a cost-effective and scalable approach (Xu et al., 2019), as previously applied to large-scale studies of human PBMCs (Kang et al., 2018). To this end, we processed lung and prostate samples jointly from three individuals, using the CST and TST protocols. We pooled tissue samples from the three individuals and then processed the pool for single nucleus extraction, thus minimizing technical batch effects and wet-lab time. After sequencing, we removed ambient RNA (Fleming et al., 2019), and performed *de novo* genotype-based demultiplexing to assign nuclei to donors (using souporcell (Heaton et al., 2020), Figure S22, **Methods**). We validated the demultiplexing by comparing the genotype-based assignments to those from an expression-based multinomial logistic classifier that assigned donor identity to each nucleus profile after training with unpooled samples of the same donors, which showed high concordance between the two approaches (accuracy 88%-96%, Figure S22). Moreover, genotype-based doublet calls were concordant with expression-based doublet calls in both lung and prostate (mean balanced accuracy 63%) (Figure S22).

## Discussion

To enable large-scale studies of the human body at single-cell resolution, we developed robust wet-lab and analytical frameworks, and applied them to generate a cross-tissue atlas from banked frozen tissues, spanning 209,126 nuclei profiles across 25 samples from eight tissue types. We benchmarked and optimized four lab protocols for nucleus extraction, developed a robust framework for data integration and cross-tissue annotation, and demonstrated multiplexing in two tissue types for scaling these approaches to larger populations of individuals.

Cross-tissue atlases allow us to characterize tissue-specific and -agnostic features of cells of a common type that serve accessory roles in tissues, such as immune and stroma cells. For example, we found a conserved dichotomy between *LYVE1*- and *HLA* class II-expressing macrophages among tissue-resident macrophages. Building on previously reported cross-tissue analyses in the mouse (Chakarov et al., 2019), our results reinforce the notion of functional specification of these two macrophage states into tissue support and tissue immunity, respectively. In mice, *Lyve1*^hi^ macrophages are localized perivascularly, whereas MHCII^hi^ macrophages are found in proximity to neurons (Chakarov et al., 2019). Future studies will need to address if the human macrophage subsets described here display similar tissue localization as their murine counterparts. The nature of the monocyte populations and differentiation trajectories that give rise to these states, and the tissue-specific signals that govern the tissue-specific ratios of *LYVE1*-*vs. HLA* class II-expressing phagocytes, remain unknown.

Our analysis further highlighted the presence of LAM-like cells in multiple tissues. Initially identified in adipose tissue (Jaitin et al., 2019), prior reports have indicated a broader distribution of LAMs (Deczkowska et al., 2020; Subramanian et al., 2021). Our data further demonstrate the prevalence of LAM-like cells across tissue contexts and pathologies and identified several common themes. In line with a model of lipid-induced differentiation of macrophages towards the LAM state (Deczkowska et al., 2020), our classifier recovered LAMs in artherosclerotic arteries, creeping fat in Crohn’s small intestine and skin acne, all pathologies characterized by lipid accumulation. Analysis of LAM-specific transcription factor activity points towards *PPARG* and *NR1H3*-driven control of the LAM expression program. These findings might suggest a model where signaling through lipid-bound receptors on macrophages such as *TREM2* up-regulate the expression of more lipid receptors, as well as of lipid modifying enzymes through *PPARG* and *NR1H3*. Importantly, as we observe LAMs in healthy organs and in other conditions not previously linked to the accumulation of lipids, such as leprosy infection, it will be important to analyze alternative tissue-specific signals that can trigger macrophage polarization towards the LAM-like state.

A key challenge in human biology is to understand how disease-associated genes affect cellular function, both for monogenic and complex diseases. We demonstrated the utility of a tissue atlas for monogenic disease biology, where we inferred the primary and secondary pathobiology of monogenic muscle diseases by analyzing skeletal, cardiac and smooth muscle. Disease groups were enriched not only for genes expressed in myocytes, but also for non-myocytes, including nervous system, immune and stromal cells (Benarroch et al., 2020). Note that we observed multiple subtypes of myocytes within one tissue, including cytoplasmic myocytes (high myoglobin (*MB*) expression and exon:intron ratio), which suggests possible specialization of different nuclei in one syncytium. Future work on syncytia can help understand nucleus heterogeneity in multinucleate cells and dissect the role of cross-nuclear circuits in muscle, the placenta (Marsh & Blelloch, 2020) and viral infection (Cifuentes-Muñoz et al., 2018), as well as whether and how dysfunction of a subset of nuclei within syncytia can result in a broader pathology. We also suggest that some disease risk genes may disrupt cell-cell interactions in the muscle, such as between myocytes and Schwann cells. Future work may further reveal the mechanisms by which disease perturbations of different genes, cell types and nuclei can elicit similar disease phenotypes.

For common complex diseases, we related cell types and programs with GWAS variants across multiple diseases and traits. We found significant enrichment in specific cell groups for multiple traits, including autoimmune diseases in T cells and NK cells, atrial fibrillation in myocytes, cardiovascular traits in pericytes and smooth muscle cells, and cognitive/psychiatric traits in Schwann cells and neurons. For over half of the traits we inspected, we observed enrichment for the same cell type across different tissues, but this enrichment was driven by both a set of common, and tissue-specific, genes. Gene programs allowed us to parse six major trait groups, whose GWAS genes were enriched in similar modules and cell types.

Advances in single cell epigenomics (Kelsey et al., 2017) and multi-omics (Fiskin et al., 2020; Ma et al., 2020; Mimitou et al., 2021) should further enable linking GWAS variants to their target genes and the cell types and programs in which they act. Recent findings indicating that a large fraction of genetic regulatory effects linked to GWAS variants can only be detected at the cellular level (Kim-Hellmuth et al., 2020; M. van der Wijst et al., 2020) suggest that cell-level eQTL maps will be essential. The experimental and computational methods we developed for a cross tissue atlas, and the biological queries we defined will provide a basis for scaling such efforts to hundreds of individuals and diverse populations.

## Methods

### Biospecimens

#### Donor and sample characteristics

All samples were selected from donors that were enrolled as part of the GTEx project. As previously described (GTEx Consortium, 2020), all GTEx tissue samples were derived from deceased donors, with study authorization obtained via next-of-kin consent for the collection and banking of de-identified tissue samples for scientific research.

While the vast majority of tissues collected from GTEx donors were preserved in PAXgene fixative (Carithers et al., 2015), which is not compatible for scRNA-seq, a subset of duplicate samples were collected from 8 tissue sites (Breast – Mammary tissue, Esophagus – Mucosa, Esophagus – Muscularis, Heart – Left Ventricle, Lung, Muscle – Skeletal, Prostate, and Skin – Sun Exposed (Lower leg)) and flash frozen. We selected 3-4 samples from each of the 8 tissues that satisfied the following criteria: (1) both genders and all available ancestral/ethnic groups are represented; (2) approximately span the age range collected for GTEx (21-70 years old); (3) RNA quality from the matched PAXgene-preserved tissue fulfilled GTEx requirements for bulk RNA-Seq (RIN ≥ 5.5); and (4) RNA-seq (from bulk matched tissue) and a donor genotype (from Whole Genome Sequencing) were available.

All GTEx samples underwent pathology review as part of that study protocol (Carithers et al., 2015), to validate tissue origin, content, and integrity. Tissues were also reviewed for evidence of any disease (cancer, infectious disease, inflammatory disease) to confirm that collected biospecimens were “normal” or non-diseased and were acceptable for inclusion in the GTEx inventory and study. For the samples selected for this study, we performed a second pathology review to provide greater detail about the broad cellular makeup of each tissue sample (**Table S1**). The accompanying H&E images for each sample are available on the GTEx portal Histology Viewer (http://gtexportal.org/home/histologyPage).

#### Sample processing

Approximately 400mg of frozen tissue was obtained from each sample to allow the application of multiple nucleus extraction methods. 20-50 mg of tissue was cut per preparation. For donor pooling, ∼10-15 mg of tissue from each donor were combined. Tissue remained frozen during cutting and weighing.

Each tissue sample was prepared as previously described for the EZ, CST, NST and TST protocols (Drokhlyansky et al., 2020). Briefly, each tissue piece was subjected to nucleus isolation and snRNA-Seq, using each of the four conditions, with either commercial EZ buffer and mechanical breakdown using dounsing (Habib et al) (EZ), or with nucleus isolation in salt-Tris buffer with detergent, either NP-40, CHAPS, or Tween-20 (NST, CST and TST receptively), with chopping to assist mechanical breakdown of the tissue (Drokhlyansky et al., 2020).

##### For the CST, NST, TST isolations

On ice, each piece of frozen tissue was placed into one well of a 6-well plate with salt-Tris buffer containing 146 mM NaCl (Cat#S6546-1L, Sigma-Aldrich), 1 mM CaCl_2_ (Cat#97062-820, VWR), 21 mM MgCl_2_ (Cat#M1028-10X1ML, Sigma-Aldrich), 10 mM Tris pH 8.0 (CAT#AM9855G, Thermo Fisher Scientific), supplemented with detergent: CHAPS (Cat#220201-1GM, EMD Millipore) at 0.49% (w/v), Tween20 (Cat#100216-360, VWR) at 0.03% (w/v), or Nonidet™ P40 Substitute (Cat#AAJ19628AP, Fisher Scientific) at 0.02% (w/v). Tissue was then chopped with Tungsten Carbide Straight 11.5 cm Fine Scissors (14558-11, Fine Science Tools, Foster City, CA) for 10 min on ice. Samples were then filtered through a 40 µm Falcon cell strainer (Thermo Fisher Scientific, cat. no. 08-771-1) into a 50ml conical tube. The well and filter were washed with an additional 1 ml of detergent-buffer solution, followed by a wash with 3ml of buffer without detergent. The ∼ 5 ml sample volume was then transferred to a 15 ml conical tube and centrifuged for 5 minutes, 500g at 4°C in a swinging bucket centrifuge with soft break. Following centrifugation, the sample was placed on ice, supernatant was removed carefully, and the pellet was resuspended in salt-Tris buffer without detergent. The nucleus solution was then filtered through a 35 µm Falcon cell strainer (Corning, cat. no. 352235). Nuclei were counted, and 7,000 nuclei of the single-nucleus suspension were loaded onto the Chromium Chips for the Chromium Single Cell 3′ Library (V2, PN-120233) according to the manufacturer’s recommendations (10x Genomics). Mouse tissues were prepared using the CST nucleus isolation protocol.

##### For the EZ protocol

Tissue samples were cut into pieces <0.5 cm and homogenized using a glass Dounce tissue grinder (Sigma, cat. no. D8938). The tissue was homogenized 20 times with pestle A and 20 times with pestle B in 2 ml of ice-cold nuclei EZ lysis buffer (NUC101-1KT, Sigma-Aldrich). Then, a volume of 3 ml of cold EZ lysis buffer was added, and sample was incubated on ice for 5 min. Sample was then centrifuged at 500g for 5 min at 4 °C, washed with 5 ml ice-cold EZ lysis buffer and incubated on ice for 5 min. After additional centrifugation as in the previous step, supernatant was removed and the nucleus pellet was washed with 5 ml nuclei suspension buffer (NSB; consisting of 1X PBS, 0.01% BSA and 0.1% RNase inhibitor (Clontech, cat. no. 2313A)). Isolated nuclei were resuspended in 2 ml NSB, filtered through a 35 μm cell strainer (Corning-Falcon, cat. no. 352235) and counted. A final concentration of 1,000 nuclei per µl was used for loading on a 10x V2 channel.

#### snRNA-seq library preparation and sequencing

Nuclei were partitioned into Gel Beads in Emulsion (GEMs) using the GemCode instrument. Lysis and barcoded reverse transcription of RNA occurred in GEMs, followed by amplification, shearing and adaptor and sample index attachment according to the manufacturer’s protocol (10x Genomics). Libraries were sequenced on an Illumina Next-Seq 500 or Nova-seq 5000/6000.

#### snRNA-seq data pre-processing

Raw sequence files were demultiplexed into the fastq format using the *cellranger mkfastq* command (*Cell Ranger* v2.1.0, 10X Genomics). A pre-mRNA reference genome was generated including both introns and exons using the commands recommended by 10X’s Cell Ranger pipeline (https://support.10xgenomics.com/single-cell-gene-expression/software/pipelines/latest/advanced/references). The “cellranger count” command was used to generate gene expression matrices.

To remove ambient RNA (Fleming et al., 2019; Heaton et al., 2020; Smillie et al., 2019; Young & Behjati, 2020), we used *CellBender* (Fleming et al., n.d.) v2’s probabilistic model to generate corrected gene expression matrices after ambient removal. *CellBender* was run on a *Terra* cloud computing environment (https://app.terra.bio) on all raw gene expression matrices using the *remove-background-v2-alpha* workflow with FPR=0.01 option. The total number of nuclei identified by CellBender was 439,772.

Following ambient RNA correction, we removed “low-quality” nucleus profiles defined as those with either <200 detected genes, >5,000 detected genes, or <400 UMIs, retaining 265,831 nuclei passing QC. Protocol- and individual-specific clusters and the clusters with high predicted doublet proportions and mixed marker sets were removed manually retaining 209,126 nuclei for all subsequent analyses.

#### Estimation of exon:intron ratios

We computed exon:intron ratios from read counts in exonic, intronic, and intergenic regions obtained with Scrinvex (https://github.com/getzlab/scrinvex) applied to the processed BAM-file. First, a collapsed annotation file was created using the “genes.gtf” created in the Cellranger v2 reference using the GTEx collapse annotation script (https://github.com/broadinstitute/gtex-pipeline/tree/master/gene_model). Then, Scrinvex was run as “scrinvex ${collapsed_gtf} ${cellranger_bam} -b ${barcodes} -o {out_file} -s {summary_file}” where cellranger_bam and barcodes are output from cellranger count.

#### Data harmonization using disentangled conditional VAEs

We used a conditional β-TCVAE (total correlation variational autoencoder) (Chen et al., 2018) to obtain disentangled representation of cells, while simultaneously factoring out unwanted sources of biological and technical variation (age, sex, self-reported ethnicity, nucleus isolation protocol, and ischemic time) from the latent representation of cells via conditioning. Python source code of β-TCVAE implemented by Yann Dubois (https://github.com/YannDubs/disentangling-vae) was adapted for tabular single-cell data by adding fully connected encoder and decoder layers. Furthermore, we added the conditioning support for continuous and categorical variables.

Conditional β-TCVAE with 64 latent dimensions was applied to log(TP10K+1)-transformed counts after subsetting the genes to 10,000 highly variable protein coding genes. Age, sex, nucleus isolation protocol, ischemic time and self-reported race/ethnicity were used for conditioning. We fitted the model on the entire dataset with beta values 1 (where β-TCVAE reduces to a standard cVAE), 2, 3, 5 and 10. Higher beta values lead to over-smoothened reconstructions and higher loss of variation in the data (Figure S3). To minimize the loss of structure in the UMAP embeddings of nuclei and to keep the distinctness of tissue-specific neighborhoods, we used beta=2.0 as a final hyperparameter. We used a softplus output activation to produce non-negative outputs. Mean squared error (MSE) was used as a loss function. UMAP embeddings and *k*-nearest neighbors (*k*-NN) graph-based clusters were inferred using a 15-nearest neighbor graph built with the mean VAE latent space embeddings of the nuclei profiles using sc.pp.neighbors(), sc.tl.umap() and sc.tl.leiden() functions of Scanpy (Wolf et al., 2018).

#### Mouse skeletal muscle snRNA-seq data pre-processing, integration and annotation

After generating the count matrices from the raw files with CellRanger (v2.1.0, 10X Genomics), we removed the “low-quality” nucleus profiles defined as those with either <200 detected genes, >5,000 detected genes, or <400 UMIs, retaining 31,904 nuclei passing QC. Processed heart and esophagus mouse snRNA-seq datasets from Drokhlyansky et al. (Drokhlyansky et al., 2020) were downloaded from https://singlecell.broadinstitute.org/single_cell/study/SCP1038/the-human-and-mouse-enteric-nervous-system-at-single-cell-resolution. Three muscle datasets were then concatenated.

Similar to the human nuclei profiles, we used a conditional β-TCVAE (Chen et al., 2018) to correct for mouse- or batch-specific effects. Conditional VAE with 64 latent dimensions was applied to log(TP10K+1)-transformed counts after subsetting the genes to the 10,000 highly variable protein coding genes. We used beta=2.0 as a hyperparameter with a softplus output activation for non-negative outputs. UMAP embeddings and graph-based clusters were inferred using a 15-nearest neighbor graph built with the mean VAE latent space embeddings of the nuclei profiles using sc.pp.neighbors(), sc.tl.umap() and sc.tl.leiden() functions of Scanpy. After differential expression via the sc.tl.rank_gene_groups() function, we manually annotated the clusters by comparing highly expressed genes with the literature-based marker lists.

For muscle disease gene set enrichment, we mapped mouse genes to human genes using the ortholog gene list from NCBI HomoloGene (https://ftp.ncbi.nih.gov/pub/HomoloGene/current/homologene.data). Fisher’s exact test (implemented in fisher Python package) was used for the enrichment test.

#### Curation of reference genes from prior studies

To curate a literature-based set of reference genes (**Table S3**), we first screened primary literature for cell-types found in histology samples for each targeted region. We then curated, at minimum, 3 marker genes that were identified by a primary experimental method (i.e., FISH) in the human target tissue of interest that overlap with significant, differentially expressed genes (one *vs*. rest, Welch’s t-test). For cell-types that are present in multiple tissues, such as certain immune cell-types or vascular endothelial cells, we included “pan” annotation markers that may have been found in any one tissue to confirm a nucleus’s putative cell-type. Finally, we resort to murine experimental confirmation, if needed (denoted in **Table S3** with murine analog gene-name).

#### Data-driven identification of marker genes using differential expression

Raw counts were converted to log(TP10K+1) values prior to differential expression analysis using the “sc.pp.normalize_total” and “sc.pp.log1p” functions of Scanpy. After accounting for the protocol- and sex-specific effects using ComBat (Leek et al., 2012) via “sc.pp.combat” function of Scanpy, we used Welch’s t-test for differential expression by running “sc.tl.rank_genes_groups” function of Scanpy separately in each tissue.

#### Cell type diversity analysis

Shannon entropy was calculated for each sample or channel across all N broad cell classes, according to the formula -∑(pi*log_2_(p_i_)), where p_i_ is the proportion of cells in cell class *i*. The TST, CST and NST protocols generally had comparable cell-type diversity scores in each tissue, with higher variability in skin, breast, and prostate (variance = 0.243 (skin), 0.585 (breast), 0.38 (prostate), < 0.1 (all other tissues)). Entropy was plotted per sample using the ggplot2 and cowplot R packages, and with the “quasirandom” function from the ggbeeswarm package to avoid overlap of plotted points. We used a linear mixed-effects model (“lmer” function from the lme4 R package) to check for association of entropy with the nuclei preparation protocol. The Fligner-Killeen test in the R stats package was used for testing the homogeneity of variances.

#### Comparison to published heart snRNA-seq

Our snRNA-seq data for heart left ventricle tissues was compared to two published snRNA-seq dataset for the heart left ventricle (Litviňuková et al., 2020; Tucker et al., 2020). For Tucker et al. (Tucker et al., 2020), we retrieved a final, annotated Scanpy AnnData object (health_human_4chamber_map_unnormalized_V3.h5ad) and raw counts (gene_sorted-matrix.mtx) from the Broad Institute’s Single Cell Portal at https://singlecell.broadinstitute.org/single_cell/study/SCP498/transcriptional-and-cellular-diversity-of-the-human-heart. *CellBender*-corrected counts for the GTEx and Tucker *et al*. datasets were converted to log(TP10K+1) values using the “sc.pp.normalize_total” and “sc.pp.log1p” functions of Scanpy. For Litviňuková et al. (Litviňuková et al., 2020), we subset the normalized, profiled nuclei from their “global” Scanpy AnnData object for the “LV” region available at https://www.heartcellatlas.org/. Highly variable genes were selected using the parameters in “sc.pp.highly_variable_genes” “min_mean=0.0125”, “max_mean=3”, and “min_disp=0.5.” Data were scaled using “sc.pp.scale” and highly variable genes were used for PCA via “sc.tl.pca”. Data were integrated between the three cohorts using Harmony (Korsunsky et al., 2019). We identified a population of ventricular myocytes in Litviňuková *et al*. that clustered with our dataset and Tucker *et al.’s* ventricular, cytoplasmic myocytes. We compared sample proportions as described below in *Proportional Analysis*.

#### Identification of contaminant transcripts from ambient RNA

We identified contaminant transcripts (from ambient RNA) by comparing the log(TP10K+1)-transformed expression values before and after the removal of ambient RNA using *CellBender* (Fleming et al., n.d.). In the comparison, we calculated the *L*2-norm of the differences between the expression values before and after ambient RNA removal for each gene separately. We used the linalg.norm function from the NumPy Python package. Norms averaged over the genes were compared between the protocols via two-sided t-test using the scipy.stats.ttest_ind function from SciPy Python package.

By this analysis, we detected markers of epithelial cell types in breast (*e.g.*, *KRT15, KRT7*), esophagus mucosa (e.g., *S100A9, S100A8, KRT13*), lung (e.g., *SFTPC, SFTPB, SFTPA1*), prostate (e.g., *MSMB, KLK3, KLK2*), and skin (e.g., *KRT10, DST, COL7A1*) as major contributors to spurious estimates of gene expression and misidentification of cell types in those tissues. In muscle tissues, genes that were highly expressed in myocytes were among the top contaminants (e.g., *MYL9, TAGLN, DES* in esophagus muscularis; *MYL2, TPM1, TNNC1* in heart, and *ACTA1, TPM2, TNNT1, TTN* in skeletal muscle).

### Comparison of snRNA-seq and scRNA-seq data

#### Lung

Samples from the Right Upper Lobe (RUL) of n=6 deceased lung transplant donors (MS, ORR, AR, Avinash Whagry, Alexander Tsankov, and Jay Rajagopal *et al*., unpublished results) were profiled for scRNA-seq using the Chromium 3’ gene expression kits (version 2 chemistry) as detailed in Slyper et al. (Slyper et al., 2020). (The snRNA-seq in this study was from Left Upper Lobe (LUL) samples). The scRNA-seq data was demultiplexed and quantified into cell-by-gene matrices using the Cellranger software version 3. Cells of high quality (minimum number of UMI=1,000, minimum genes detected=400 and maximum percentage of mitochondrial genes detected=10%) were retained, gene counts were log normalized (“NormalizeData”) after total sum-scaling with a multiplicative factor of 10^5^ resulting in log(TP10K+1) units, highly variable genes calculated (“FindVariableGenes”) and scaled (“ScaleData”) using standard functions of Seurat version 3. Data from all donors were merged and subjected to dimensionality reduction (number of principal components=20), a *shared*-NN graph was constructed (*k*=20) followed by Louvain clustering (resolution =1.2) using default parameters as implemented in Seurat version 3. No substantial batch effects were observed after merging and clustering, as cells from different individuals clustered by cell type identity. Cell clusters were manually annotated at a level of broad cell categories by expression of canonical cell type markers, as previously described (Muus et al., 2021). 15 cell classes were shared with the snRNA-seq data. All 46,751 scRNA-seq profiles from the 15 shared cell categories were compared with 11,983 nuclei profiles from the TST protocol matching the same cell categories: immune (alveolar macrophage), immune (macrophage), epithelial cell (ciliated), immune (NK cell), epithelial cell (alveolar type II), epithelial cell (alveolar type I), epithelial cell (basal), fibroblast, pericyte/SMC, immune (T cell), epithelial cell (club), immune (mast cell), immune (B cell), endothelial (vascular) and endothelial (lymphatic). Goblet and secretory cells, smooth muscle cells, monocytes and mesothelium cell categories were found in scRNA-seq but not snRNA-seq.

#### Skin

##### Collection and experimental processing

Samples for scRNA-seq were obtained from the abdomen, from discarded excess tissue removed during abdominoplasty (n=2, IRB 2017P001913/PHS). (Samples for snRNA-seq in this study were from the left or right leg, 2cm below the patella on the medial side as described above).

For processing fresh skin, hair and fat were removed and tissue was cut into small pieces, followed by two washes with 30 mL cold PBS in a 50 mL tube. Skin was then dissociated for single cell suspension using either the Miltenyi Biotec Whole Skin Dissociation Kit, human (cat no. 130-101-540) according to manufacturer’s guidelines, or with dissociation medium containing 5 mL (2%) FBS in RPMI, 100 µg / mL Liberase TM (Sigma Aldrich, cat. no. 5401127001), 100 µg / mL Dispase (Sigma Aldrich, cat. no. 4942078001), 100 µg / mL DNase I (Sigma Aldrich, cat. no. 11284932001). For this protocol, samples were incubated in a rotating 50 mL tube, at 37°C for 3 hours, with pipetting every hour. After incubation, large undigested pieces were removed, and suspension was placed in a new tube, spun down for 3 minutes at 400g, followed by a wash with cold PBS. RBC lysis was performed using 500 µL ACK (Thermo Fisher Scientific, cat. no. A1049201) on ice for 1 minute, followed by a wash with 8 mL cold PBS. Samples were then pelleted and treated with 200 µL TrypLE (Life Technologies, cat. no. 12604013) for two minutes while pipetting, and washed with 1 mL 10% FBS in RPMI to quench, followed by a wash with 1 mL of cold PBS. Pellet was then resuspended in 1 mL of 0.4% BSA (Ambion, cat. no. AM2616) in PBS, and filtered through a 70 µm strainer (Falcon, cat. no. 352350). Cells were counted by mixing 5 µl of Trypan blue (Thermo Fisher Scientific, cat. no. T10282) with 5 µl of the sample and loaded on INCYTO C-Chip Disposable Hemocytometer, Neubauer Improved (VWR, cat. no. 82030-468). Cells were loaded onto a 10x Genomics Single-Cell Chromium Controller, using V2 3’ GEX kit.

##### Computational analysis

Sequencing files were demultiplexed and quantified using Cellranger 2.0.1. and human reference genome version GRCh38-1.2.0 using commands “cellranger mkfastq” and “cellranger count”, respectively. We then ran CellBender as described above to remove ambient RNA, and retained high quality cells (at least 200 genes detected and at most 20% of reads mapping to mitochondrial genes per cell) for a total of 27,199 cells. We performed standard processing using the Seurat R package (v3) including total sum normalization and log transformation with a multiplicative factor of 10^5^ (“NormalizeData”) resulting in log(TP10K+1) units, determination of variable genes (“FindVariableFeatures” using the vst method), scaling (“ScaleData”), dimensionality reduction by principal component analysis (“RunPCA”, k=50) and graph-based clustering (“FindNeighbors” for *k*=50 and “FindClusters” at a resolution of 1). Using differential gene expression (“FindAllMarkers”) and annotation by curated marker genes, we annotated 15 cell classes. Of these, 11 were shared with this study’s skin snRNA-seq broad cell types: endothelial cell (lymphatic), endothelial cell (vascular), epithelial cell (basal keratinocyte), epithelial cell (suprabasal keratinocyte), melanocyte, immune (Langerhans), sweat gland cell, fibroblast, pericyte/SMC, immune (T cell) and immune (DC/macrophage). Adipocytes, cornified keratinocytes and sebaceous gland cells were unique to the snRNA-seq data, and a small population of neuron-like cells and preadipocytes were found only in the scRNA-seq data.

#### Prostate

Published scRNA-seq data from three anatomical zones from coronal sections of the whole prostate separated post-cystoprostatectomy was downloaded from GEO GSE117403 (D17, D27, Pd). (SnRNA-seq in our study was derived from any representative, non-nodular region, avoiding seminal vesicles). All samples were combined and processed as described above for skin scRNA-seq, resulting in a total of 82,822 cells annotated to 11 shared cell classes: epithelial cell (basal), epithelial cell (luminal), epithelial cell (club), epithelial cell (Hillock), endothelial cell (lymphatic), endothelial cell (vascular), pericyte/SMC, neuroendocrine, immune (DC/macrophage), fibroblast, myocyte (smooth muscle). Additional epithelial subsets identified only in the scRNA-seq data were *FOXI1*^+^, *SEMG1*^+^, stressed and cycling cells, and only the snRNA-seq data had lymphocytes, mast and Schwann cells. The discrepancy in the immune cells may be because the scRNA-seq data was enriched for epithelial and mesenchymal cells.

#### Mapping scRNA-seq and snRNA-seq profiles with a random forest classifier

To compare profiles between scRNA-seq and snRNA-seq datasets, a multi-class random forest classifier was trained on nuclei (cells) profiles as the training set with the function “*randomForest”* from the *randomForest* R package, and used to predict the classes of cells (nuclei) profiles with the “predict” function, with the shared broad cell classes (15 in lung, 11 in skin and 11 in prostate) as class labels (cells/nuclei unique to only scRNA-Seq/snRNA-Seq were removed prior to analysis). The smaller of up to 70% of the cell class size or 1,000 cells were randomly sampled per class to form a training set, and genes were selected as the intersection of the top 3,000 highly variable genes from both nuclei and cell profiles. Highly variable genes were derived using the *FindVariableFeatures* function with “vst” as the selection method. Broad cell types were accurately predicted in scRNA-Seq by a classifier trained on snRNA-seq with a median accuracy of 89%, 89%, 89.3% across skin, lung and prostate, respectively (Figure 2C-E, Figure S8 F-H). There was a similarly high accuracy for predicting snRNA-seq broad cell types by a classifier trained on scRNA-Seq (median 88% in skin, 90% in lung), suggesting that cell intrinsic programs are well-preserved between fresh and frozen samples and techniques.

#### Correlation analysis between scRNA-seq and snRNA-Seq profiles

Pseudobulk expression profiles were correlated between cells and nuclei. For each broad cell type category separately, pseudobulk profiles were computed as follows: The gene (rows) by cell (column) matrix of raw counts for each sample was first scaled per-column by the total counts in each cell (nucleus) and then multiplied by 10^6^ to derive counts per million (CPM) for each cell (nucleus) profile. CPMs for each gene were then averaged across cells (row-wise) for each broad cell type. The resulting expression vector per sample is the pseudobulk profile. Spearman correlation was computed for the log_2_-transformed pseudobulk gene expression profiles of the cell and nucleus data of interest separately for protein-coding and long non-coding RNA genes selected as follows. The total set of human genes was downloaded from the Ensembl database (GRCh38 v103, 04/09/2021), using the BiomaRt R package. Mappability scores for GRCh38, computed using the ENCODE pipeline (Derrien et al., 2012), were obtained from GTEx and averaged across gene bodies. The complete set was then partitioned into genes with both pre-mRNA and mRNA mappability > 90%, and then partitioned into protein coding and long non-coding RNA genes based on their Ensembl biotype classification.

To identify genes that deviate from the overall expected similarity, a linear model was fit to compute residuals using the command “resid(lm(cells-nuclei ∼ 0))”, where the “cells’’ and “nuclei” are log_2_-transformed pseudobulk gene expression profiles of the scRNA-seq and snRNA-seq data, respectively for either protein-coding or long non-coding RNA genes. Divergent genes were defined as those with residuals greater than the 97.5th or lesser than the 2.5th percentile, respectively.

Poly-A content was computed using the *BSgenome.Hsapiens.UCSC.hg38* R package by searching for stretches of at least 20 consecutive adenine “A” bases to qualify as one “polyA” unit. The total polyA width for a gene was defined as the total number of As in such units. The gene length was also computed using the same package.

#### Comparison of tissue dissociation induced-stress signature scores

Single-cell dissociation protocols have been reported to cause dissociation-induced gene expression (Denisenko et al., 2020; van den Brink et al., 2017). We scored a published dissociation signature (van den Brink et al., 2017) in snRNA-seq and scRNA-seq profiles using the “AddModuleScore” function in Seurat for each shared cell classes across snRNA-seq and scRNA-seq datasets. The score was computed on the normalized gene expression units of log(TP10K+1).

#### Proportion analysis

Dirichlet regression was used for proportion analysis via DirichletReg R package version 0.7.0. For analysis of broad cell type proportions, “proportions ∼ protocol + tissue + Age + Sex” formula was used with common parameterization, where lung and CST were used as references for protocol and tissue categorical variables.

We applied this to compare proportions of (**1**) broad cell types, (**2**) myeloid states, (**3**) comparison between scRNA-seq and snRNA-seq, and (**4**) comparison of heart snRNA-seq studies. For myeloid proportions, the formula “proportions ∼ tissue + protocol | protocol” was used with alternative parameterization to correct both mean and precision estimates for the protocol effects. We fitted eight models, each with a different tissue used as a reference. For comparing proportions between protocols, the formula “counts ∼ protocol | protocol” was used with the alternative parameterization. For comparison of scRNA-seq *vs*. snRNA-seq, scRNA-seq was used as the reference. For comparison between snRNA-seq datasets in heart, this study’s data was used as the reference. Samples with fewer than 30 nuclei profiles were excluded from the proportion models. In the comparison of proportions among heart snRNA-seq studies, data from this study’s EZ protocol, Litviňuková et al. (Litviňuková et al., 2020) and Tucker et al. (Tucker et al., 2020) showed a higher proportion of muscle cells in comparison to the CST, NST and TST protocols (Dirichlet regression, LRT with CST as baseline: Benjamini-Hochberg FDR=2.5*10^-4^ (EZ), 0.002 (Tucker et al., 2020), 6.3*10^-4^ (Litviňuková et al., 2020)), and lower proportions of endothelial cells (adj. *P*=0.002 (Tucker et al., 2020), 4*10^-10^, (Litviňuková et al., 2020)). The immune and adipose compartments were comparable in proportions (Benjamini-Hochberg FDR>0.1) across protocols and studies, however, Litviňuková et al.(Litviňuková et al., 2020) had lower proportions of fibroblasts (Benjamini-Hochberg FDR= 0.1). Taken together, all four protocols retain expected cell groups shared with published protocols.

#### Normalization of proportions

To account for the variation in total number of nuclei profiled in each tissue (e.g. 5,327 in skin and 36,574 in heart) in Figures S2D and S13F, where the proportions of nuclei from each tissue for each cell type are visualized, we normalized each proportion by the total number of nuclei profiled in that tissue. To visualize the proportion of nuclei from tissue q for the cell type p, instead of N_p,q_/ ∑_i_N_p,i_ where N_x,y_ represents number of nuclei for cell type x in tissue y, we used (N_p,q_/ ∑_i_N_i,q_) / (∑_j_ (N_p,j_/ ∑_i_N_i,j_)) formula. Therefore these visualizations approximately show what the proportions would be, if the same number of nuclei were recovered in each tissue.

#### Co-embedding of bulk and snRNA-Seq pseudobulk profiles

GTEx v8 bulk RNA-seq expression data were downloaded from the GTEx portal https://gtexportal.org and converted into an AnnData object concatenated with the pseudobulk profiles of the nuclei profiled in this study. The number of highly variable genes (calculated only on the pseudobulk data), the number of PCs, the number of neighbors, *k*, used in the *k*-NN graph and the similarity metric used in the *k*-NN graph were determined by the Tree of Parzen Estimators (TPE) random hyperparameter search method implemented in hyperopt Python package (v0.2.5) (Bergstra et al., 2013). The accuracy metric that was optimized in the search was defined as the fraction of bulk neighbors of pseudobulk data points that are from the same tissue site. Final parameters used were 3,522 HVGs, 70 PCs, 42 nearest neighbors and *L*1 as the *k*-NN distance metric. Bulk and pseudobulk samples were integrated using the PyTorch implementation of harmony (v0.1.5, https://github.com/lilab-bcb/harmony-pytorch) (Korsunsky et al., 2019) in each iteration.

#### Sample embeddings based on cell type compositions

Broad cell type proportions in each profiled sample were used to construct an *k*-mutual nearest neighbors graph of channels in Scanpy. We used *k*=7 and Fruchterman-Reingold layout where connected components of the graph were considered clusters.

#### Preprocessing and visualization of myeloid nuclei profiles

We used log(TP10K+1)-transformed counts to estimate 2,000 highly variable genes. PCA was fitted using the HVGs and a *k*-nearest neighbors (*k*-NN, *k*=15) graph was constructed using 50 PCs. We integrated all myeloid nuclei profiles from eight tissues using the PyTorch implementation of Harmony (Korsunsky et al., 2019) (https://github.com/lilab-bcb/harmony-pytorch) to correct for individual- and protocol-specific variation in the PC space. A PAGA (Wolf et al., n.d.) graph was inferred based on the myeloid annotations and UMAP visualization was performed with PAGA initialization. Proportion bar plots and dotplots were generated using the Python implementation of the ggplot framework, plotnine version 0.7.

#### Classification of lipid-associated macrophages

We used a published annotated adipose scRNA-seq study (Jaitin et al., 2019) to train a logistic classifier that can classify profiles into three groups: LAMs, non-LAM macrophages and other cell types. We performed 5-fold stratified cross-validation that maximizes weighted F1-score, using the LogisticRegressionCV function from the scikit-learn Python package to optimize the *L*2-regularization parameter. To take highly imbalanced class frequencies, we employed a weighting scheme, where rarer classes (*e.g.*, LAMs) contributed more to the overall loss function. log(TP10K+1)-transformed expression values of 17,612 protein coding genes were used in the training.

#### Preprocessing of the external datasets used in LAM classification

Datasets with IDs GSE115469, GSE117403, GSE127246, GSE128518, GSE153643, GSE131685, GSE131778, GSE131886, GSE140393, GSE143380, GSE143704, GSE144085, GSE150672, GSE153760, GSE156776, GSE159677 were downloaded from Gene Expression Omnibus (GEO). Placenta and decidua datasets were downloaded from EBI Single Cell Expression Atlas (https://www.ebi.ac.uk/gxa/sc/home) using the E-MTAB-6701 accession via the “sc.datasets.ebi_expression_atlas” function of the Scanpy Python package. Human dataset of the enteric nervous system was downloaded from the Single Cell Portal (URL: https://singlecell.broadinstitute.org/single_cell/study/SCP1038/the-human-and-mouse-enteric-nervous-system-at-single-cell-resolution). log(TP10K+1)-transformed counts were used for classification.

#### Gene set enrichment of LAM markers

LAM profiles were compared to other myeloid profiles by ranking genes by their mean z-scored log(TP10K+1) expression values. Top 100 genes were then evaluated for GO term enrichment using the “sc.queries.enrich” function of Scanpy (Wolf et al., 2018) after excluding genes from the ferritin (*FTL, FTH1*) and metallothionein (*e.g.*, *MT1A, MT2A*) families. Finally, REVIGO (Supek et al., 2011) was used to project enriched GO terms in 2D, while preserving GO term similarities.

#### Differential expression of MΦ *LYVE1*^hi^ *vs*. MΦ *HLA*II^hi^ states

To compare MΦ *LYVE1*^hi^ *vs* MΦ *HLA*II^hi^ states in heart and lung, a Wald test with a negative binomial regression model was used with the formula “∼ 1 + cell_state + participant_id + protocol + log_number_of_genes’” on raw counts. In esophagus mucosa, participant_id and protocol covariates were not used since most MΦ *HLA*II^hi^ nuclei were detected in the GTEX-15SB6 CST sample.

#### Receptor-ligand analysis

CellPhoneDB version v2.1.4 was used for receptor-ligand analysis. “cellphonedb method statistical_analysis” command was used for the analysis of each tissues with the arguments “meta.tsv counts.tsv --counts-data hgnc_symbol --project-name tissue --threads 30 --subsampling --subsampling-num-cells 50000 --subsampling-log false”. Receptor-ligand plots were created with the ggraph R package using the “sugiyama” graph layout.

#### Gene set enrichment of monogenic muscle disease genes

Fisher’s exact test implemented by the “fisher” Python package was used to test for the enrichment of muscle disease genes in broad and granular cell type markers.

#### Linking snRNA-seq to genetic variation using GWAS/eQTL enrichment analysis

To test whether the expression of genes in GWAS loci associated with a given complex disease or trait is enriched in specific cell types more than expected by chance, was used to compare the cell type specific expression of genes mapped to known GWAS loci for a complex trait of interest to a background distribution of GWAS loci. GWAS variant associations were obtained from Open Targets Genetics (Ghoussaini et al., 2021) from the NHGRI-EBI GWAS catalog and UK Biobank GWAS. Only GWAS with at least part of their samples from European ancestry and genome-wide significant associations (P<5×10^-8^) were considered. 23 traits were selected (**Table S10**) based on pathophysiology related to one of the 8 tissues in this study. Different GWAS that mapped to the same trait were considered together for the given trait (**Table S10**). For each tissue, a background of GWAS loci (null set) was defined as the GWAS variants for all traits excluding the set of traits selected for the particular tissue (on average 71,411 variants). The enrichment analysis consisted of three consecutive steps, as follows:

##### Gene mapping

Genes were mapped to trait-associated loci for the selected and background (null) traits, using the 95% credible set of fine-mapped *cis*-eQTLs and *cis*-sQTLs from each of the 49 GTEx tissues (v8) computed using DAP-G (Barbeira et al., 2021; GTEx Consortium, 2020; Wen et al., 2017). Specifically, all variants in LD (r^2^>=0.8) with each of the GWAS variants were identified using the GTEx whole genome sequencing variant calls as the reference panel (GTEx Consortium, 2020) and PLINK 2.0 (Plink --bfile 1KG_chr_files --r2 --ld-snp-list variant_list_file --ld-window-kb 5000 --ld-window-r2 0.8 --ld-window 99999). If a GWAS variant was not present in the GTEx samples, LD proxies for the variant were searched in the 1000 Genomes Project panel (Zheng-Bradley et al., 2017) at r^2^>0.8, and these proxies were subsequently checked for LD variants in the GTEx panel. GWAS associations whose variant or LD proxy variants were significant eVariants or sVariants (FDR<0.05) in any of the 49 GTEx tissues were assigned the corresponding eGene/s and sGene/s to their locus. We further included genes mapped to GWAS variants based on the ‘bestLocus2Genes’ mapping from Open Target Genetics, which included additional omic data (e.g., Hi-C) and predicted deleterious protein coding variants in LD with the GWAS variant (Ghoussaini et al., 2021). To avoid inflation of enrichment due to LD between GWAS variants, GWAS variants that shared LD proxy variants, or eGenes or sGenes were collapsed into a single locus. This was done separately for the GWAS variants for each selected trait and for all null traits per tissue. On average, 40% of the null variant sets and 80% of the 23 selected traits had at least one mapped gene, and of the mapped loci, on average 2 genes mapped per locus, ranging from 1-37 for the selected traits and 1-170 for the null traits.

##### Locus scoring

For each combination of trait of interest, tissue, and cell type, each GWAS locus was first scored based on the fraction of log_2_ fold-change > 0.5 and FDR < 0.1 of all genes mapped to the locus.

##### Cell type specificity significance estimate

The significance of the cell type specificity scores of a GWAS locus set for each cell type was assessed against the distribution of values of the background GWAS loci from a Bayesian Fisher’s exact test. Specifically, the counts of GWAS loci with their scores greater than the 95^th^ percentile of scores from the background loci for a given cell type were modeled as Binomial distributions, with the parameters (*θ*_1_, *n*_1_) and (*θ*_2_, *n*_2_) for the GWAS locus set and background loci, respectively, where *n*_1_and *n*_2_ are the total number of loci in the GWAS locus set and background, respectively. Uninformative uniform priors were specified for *θ*_1_ and *θ*_2_, leading to the conjugate Beta distributed posteriors. Next, fold enrichment was defined as the ratio of *θ*_1_ and *θ*_2_, and the 95% credible interval was constructed from 10^6^ Monte Carlo draws from the posteriors. Of 23 selected traits, 21 had 5 or more GWAS loci with at least one mapped gene, which were analyzed for cell type specificity enrichment (**Table S10**). Multiple hypothesis correction was applied tissue-wide (correcting for all cell types tested per tissue and trait) and experiment-wide (correcting for all traits by cell types and tissues tested) using the Benjamini-Hochberg FDR.

#### Variant-to-gene mapping using the OpenTargets Genetics API

Publicly available JSON files (v20022712) and the GraphQL API (v20.02.07) of the OpenTargets Genetics (OTG) portal were used to obtain genes mapped to the independent GWAS loci in Figures 5 and 6. For study level information (*e.g.*, study IDs, number of individuals, number of significant loci), JSON files were downloaded from https://ftp.ebi.ac.uk//pub/databases/opentargets/genetics/20022712/lut/study-index. For variant-to-gene mapping and variant-level details, such as Locus2Gene scores, the manhattan() and studyInfo() functions were used through the GraphQL API endpoint http://genetics-api.opentargets.io/graphql using the sgqlc Python package. The manhattan() function provides the list of all significant and independent lead SNPs as well as the genes associated with them using the Locus2Gene scoring model for the studies stored in the OTG portal. Closest protein coding genes were used in cases where Locus2Gene score was not available. GWAS with fewer than two significant loci or fewer than 3,000 individuals were excluded. UK BioBank (UKBB) traits containing “None of the above” were also removed. For the remaining studies, only the largest GWAS (based on the nInitial field of the study) was considered for a given phenotype, resulting in 4,062 studies.

#### Module-based GWAS enrichment

To infer gene modules enriched with GWAS risk genes and the cell types expressing these modules, genes were first hierarchical clustered with the complete linkage method and a correlation distance, i.e. *dist*(*g*_1_, *g*_2_) = 1 − *r*(*g*_1_, *g*_2_) where *r* is the Pearson correlation coefficient between genes *g*_1_ and *g*_2_, calculated with the scipy.cluster.hierarchy.linkage function from the scipy Python package. Models were fit separately in each tissue using all protein coding genes. To speed up the inference, clustering was performed in PC space, where gene loadings for 100 PCs were taken into account in the distance calculations. Gene modules were obtained at different resolutions by cutting the linkage tree at 100 different levels starting from only two clusters (*i.e.*, modules) to a highly granular level, where the number of clusters is equal to half of the number of genes. In a post-processing step, modules that were exactly the same or had fewer than three genes were removed. Next we scored all cells using each module as a signature to quantify average expression of the modules using the scanpy.tl.score_genes() function. Finally, gene module enrichment with the GWAS risk genes was estimated by testing all modules against all GWAS phenotypes using Fisher’s exact test implemented in “fisher” Python package. Final cell type enrichment score was defined as the product of the gene set overlap metric (f-score) and the signature score (module expression). To prioritize the associations with high expression and high gene set overlap we used additional cutoffs such as at least four genes in the intersection of the GWAS risk genes and the module genes and enrichment score higher than 0.15.

#### Preprocessing and demultiplexing of pooled samples

For genotype-based demultiplexing and doublet detection, souporcell (Heaton et al., 2020) was used as available in a Docker image from Cumulus (B. Li et al., 2020) (Cumulus version 2020.03, souporcell version eeddcde). Vartrix v1.1.20 (https://github.com/10XGenomics/vartrix) was used instead of the older version included in the Docker image. Souporcell was applied to four samples with the following command line arguments “-t 32 -o outputdir --min_alt 10 --min_ref 10 --restarts 100 --common_variants common_variants_grch38.vcf -k 3”. Using the unpooled lung and prostate samples from the same individuals, an expression-based multinomial logistic classifier was trained to predict the individuals. The “LogisticRegression(max_iter=500, penalty=’l2’, solver=’liblinear’, C=0.001, class_weight=’balanced’)” function from the scikit-learn Python package (v0.24.1) was used for the training. scrublet Python package for doublet detection was used to compare to the souporcell doublet detection method.

## Acknowledgments

We thank Leslie Gaffney and Anna Hupalowska for help with figure preparation. We thank Alexander Tsankov, Avinash Waghray and Jayaraj Rajagopal for assistance with the lung scRNA-seq data. We thank Tyler Harvey and Daniel G. MacArthur for sharing expertise on muscle diseases.

## Author contributions

Conceptualization: K.G.A., O.R.R., A.R.; data curation: A.S., D.D., F.A., G.E., O.A., P.A.B., S.A.; formal analysis: A.S., G.E., J.Wan., J.M.R., O.A., S.A.; funding acquisition: K.G.A., G.G., O.R.R., A.R., A.V.S.; investigation: E.D., M.S., N.V.W., T.S.W., D.D., J.Wal., M.S.C., O.K., J.L.M.; methodology: A.V.S., A.S., F.A., G.E., J.M.R., J.Wan., E.F.; resources: F.A.; software: G.E., S.A., A.S., J.M.R., J.Wan.; supervision: A.R., A.V.S., F.A., K.G.A., O.R.R.; validation: S.A., A.S., G.E.; visualization: S.A., G.E., A.S., J.Wan.; writing of original draft: A.S., E.D., S.A., G.E., E.F., F.A., K.G.A., A.V.S., J.M.R., J.Wan., A.R.; and writing review & editing: G.E., F.A., K.G.A., A.R., A.G., E.F., A.V.S.

## Data and materials availability

Raw sequence data are available at AnVIL (https://anvil.terra.bio/#workspaces/anvil-datastorage/AnVIL_GTEx_V9_hg38; dbGaP accession phs000424). Gene expression matrices are available from the GTEx Portal (www.gtexportal.org) and the Single Cell Portal (https://singlecell.broadinstitute.org/single_cell/study/SCP1479/single-nucleus-cross-tissue-molecular-reference-maps). The code used in the analysis is available at https://github.com/broadinstitute/gtex-single-nucleus-reference.

---

**Fig. S1.**
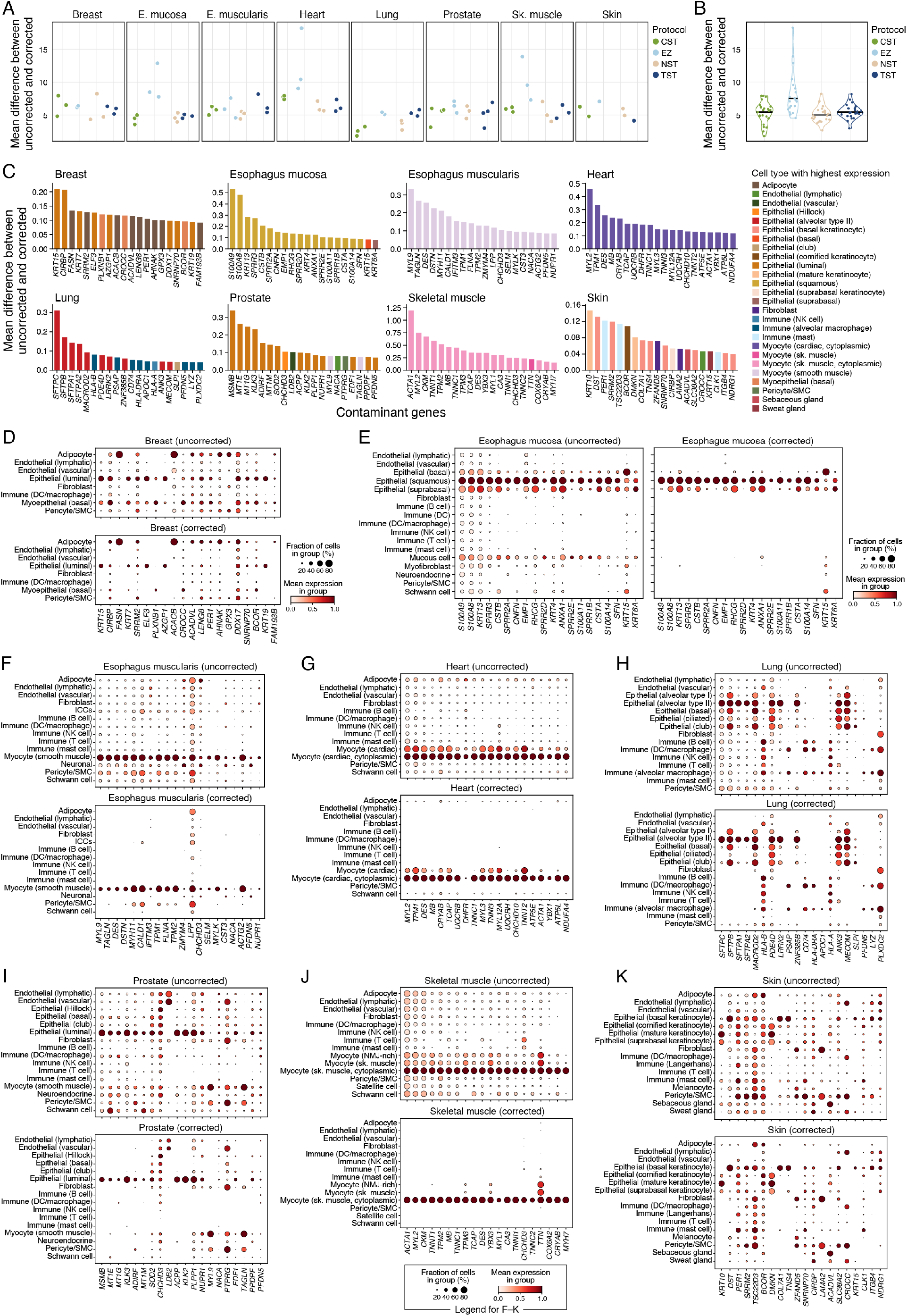
Ambient RNA correction improves cell type specificity of gene markers. (**A, B**) Ambient contamination level. Average contamination level (*y*-axis, *L*2-norm of the difference between uncorrected and corrected log(TP10K+1)-transformed expression levels averaged across genes) for each protocol and tissue **(A)** or for all samples from one protocol **(B)**. Horizontal bar in violin plots: median. **(C–K)** Potential top sources of the ambient RNA. (**C**) Mean difference between corrected and uncorrected expression values (*y*-axis) for the top genes (*x*-axis) in each tissue, colored by their cell type of highest expression in the tissue (color legends). **(D–K)** Mean expression (circle color) and fraction of expressing cells (circle size) for each of the contaminant genes from (C) (columns) in each cell type (rows) in each tissue before and after the correction for ambient RNA (as labeled on top).

**Fig. S2.**
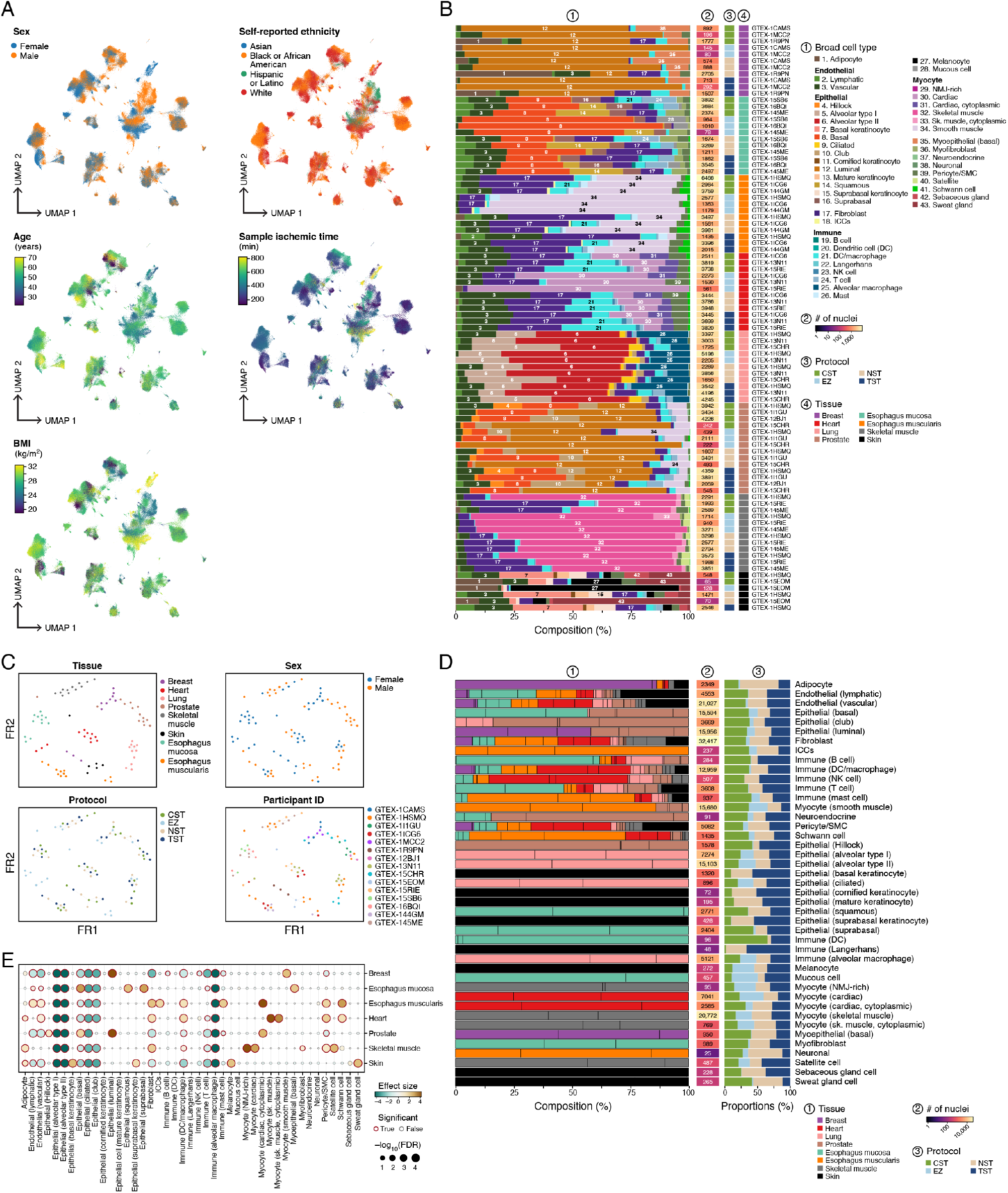
Characterization of the cross-tissue snRNA-seq atlas. **(A)** Sample characteristics in integrated atlas. UMAP visualization of snRNA-seq profiles (as in Figure 1B), colored by donor sex, self-reported ethnicity, age, BMI and sample ischemic time. **(B)** Distinct cell type composition of each tissue. Overall proportion of cells (%) of each type (color legend) in each experiment (rows, 1), along with number of nuclei profiled (2), lab protocol (3), tissue (4) and specimen (label on right). Numbers on bars: broad cell type numbers (color legend). **(C)** EZ isolation protocol is more distinct. Force-directed graph layout embedding of samples (dots), where sample similarity is calculated based on cell type composition (**Methods**). Samples are colored by tissue, sex of the donor, nuclei prep and participant ID. **(D,E)** Tissue specific and shared cell types. **(D)** Overall proportion of cells from each cell subset (bars, (1)) that are derived from each tissue (colors; normalized for the total number of nuclei profiled in each tissue, **Methods**), along with total number of nuclei from that cell type (2), and the proportion of nuclei from each protocol (3). Black vertical lines: relative proportion of nuclei from each individual. **(E)** Significance (–log_10_(Benjamini– Hochberg FDR), circle size) and effect size (circle color) of the differential abundances of each cell type (columns) in each tissue (rows) compared to lung as a reference. Samples with fewer than 30 nuclei are not shown.

**Fig. S3.**
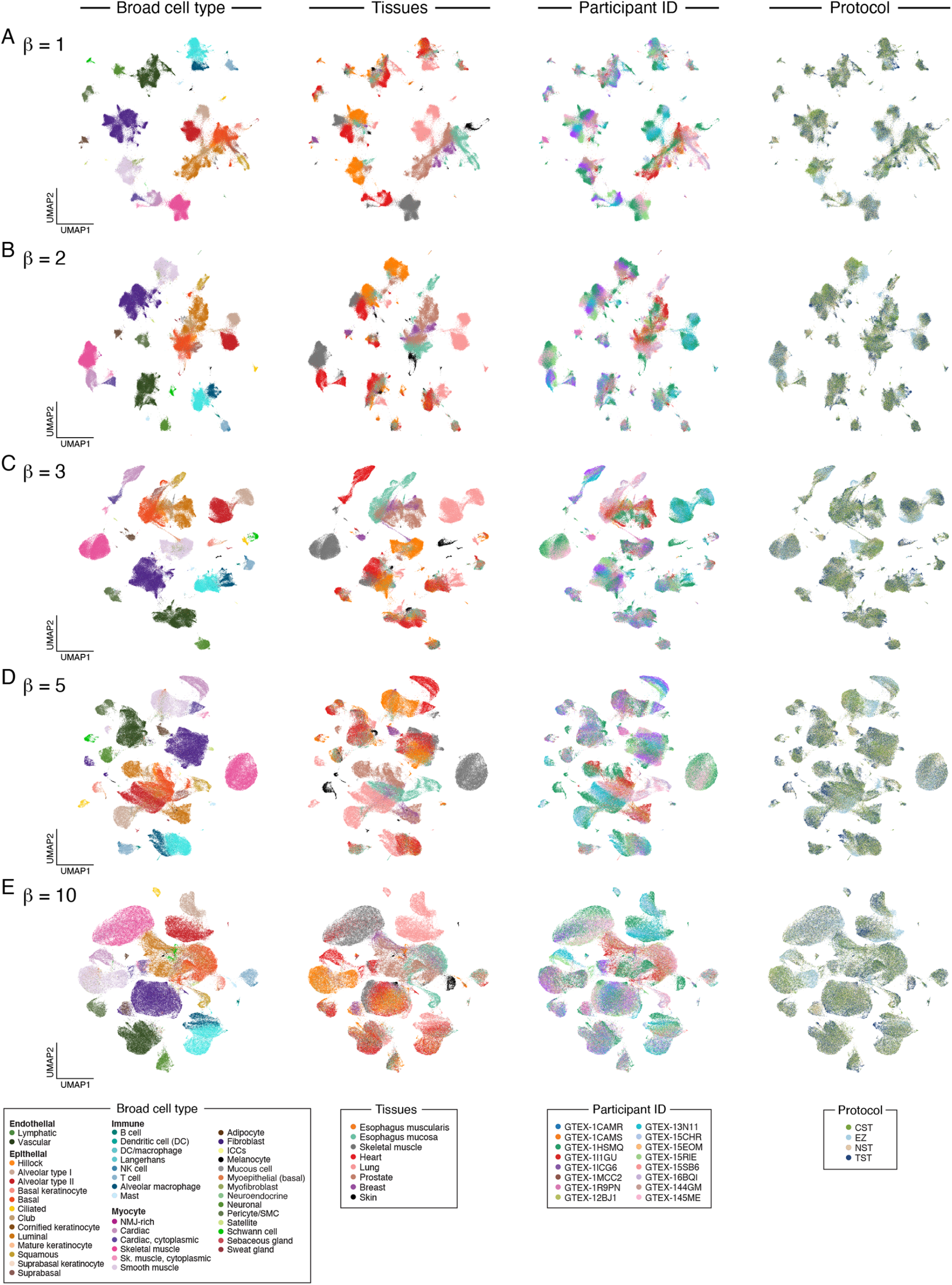
Hyperparameter sweep for the beta parameter of the conditional variational autoencoder. (**A–E**) Higher beta values over-smoothened data representations. UMAP representation of single-nucleus profiles (dots) colored (from left to right) by broad cell type, tissue, individual, or protocol using the mean latent space coordinates from conditional beta-TCVAEs trained with beta hyperparameter values of (**A**) 1.0 (**B**) 2.0, (**C**) 3.0, (**D**) 5.0, and (**E**), 10.0. The beta-TCVAE with beta = 2.0 was selected to minimize loss of granularity.

**Fig. S4.**
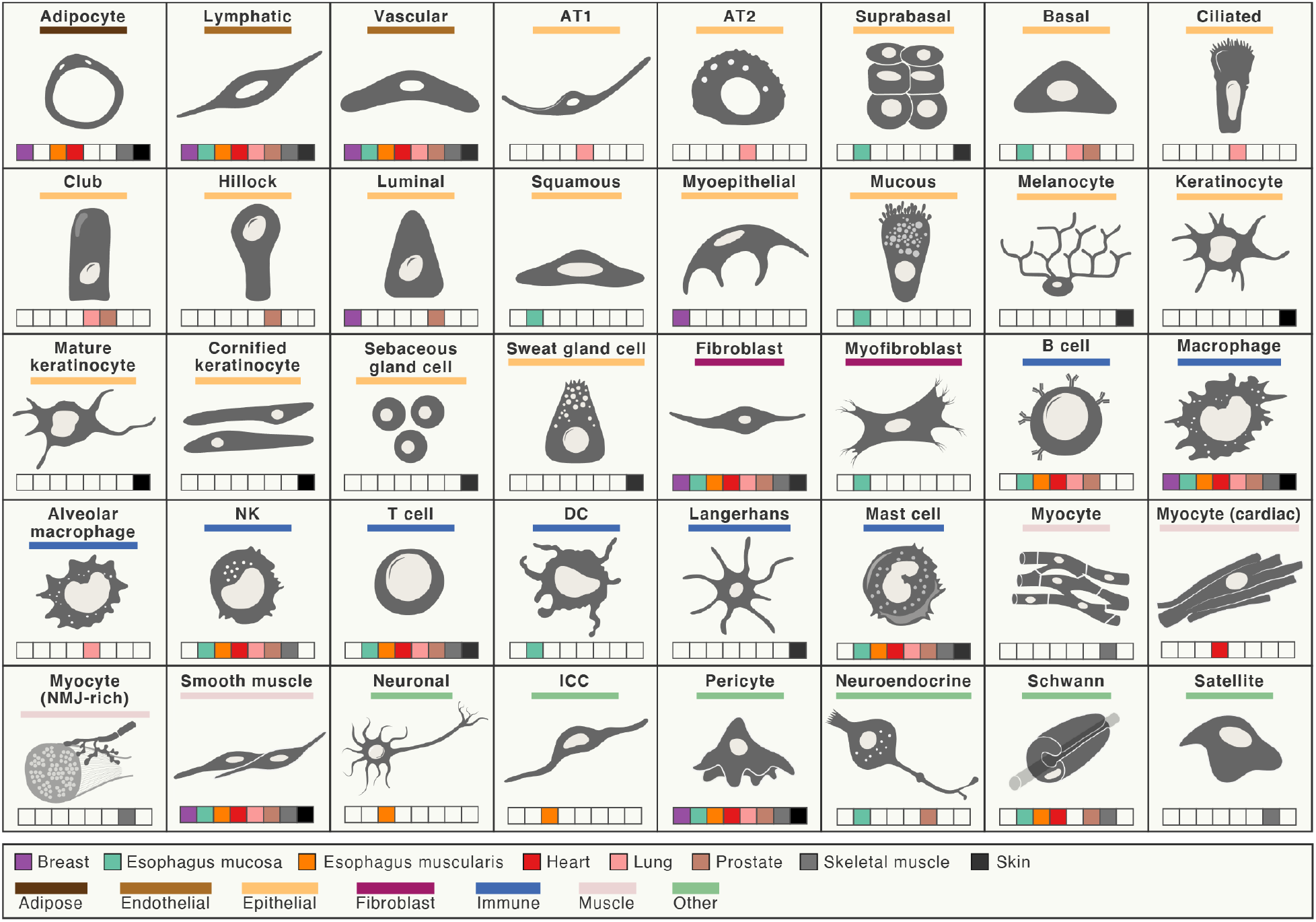
Broad cell types in the cross-tissue atlas. Illustration of 40 (of 43) broad cell types in the cross-tissue atlas labeled by cellular compartments (horizontal color line) and tissues in which the cell type is detected (colored boxes). Cytoplasmic myocytes are not shown and suprabasal epithelial cells from esophagus mucosa and skin are shown as a single cell type.

**Fig. S5.**
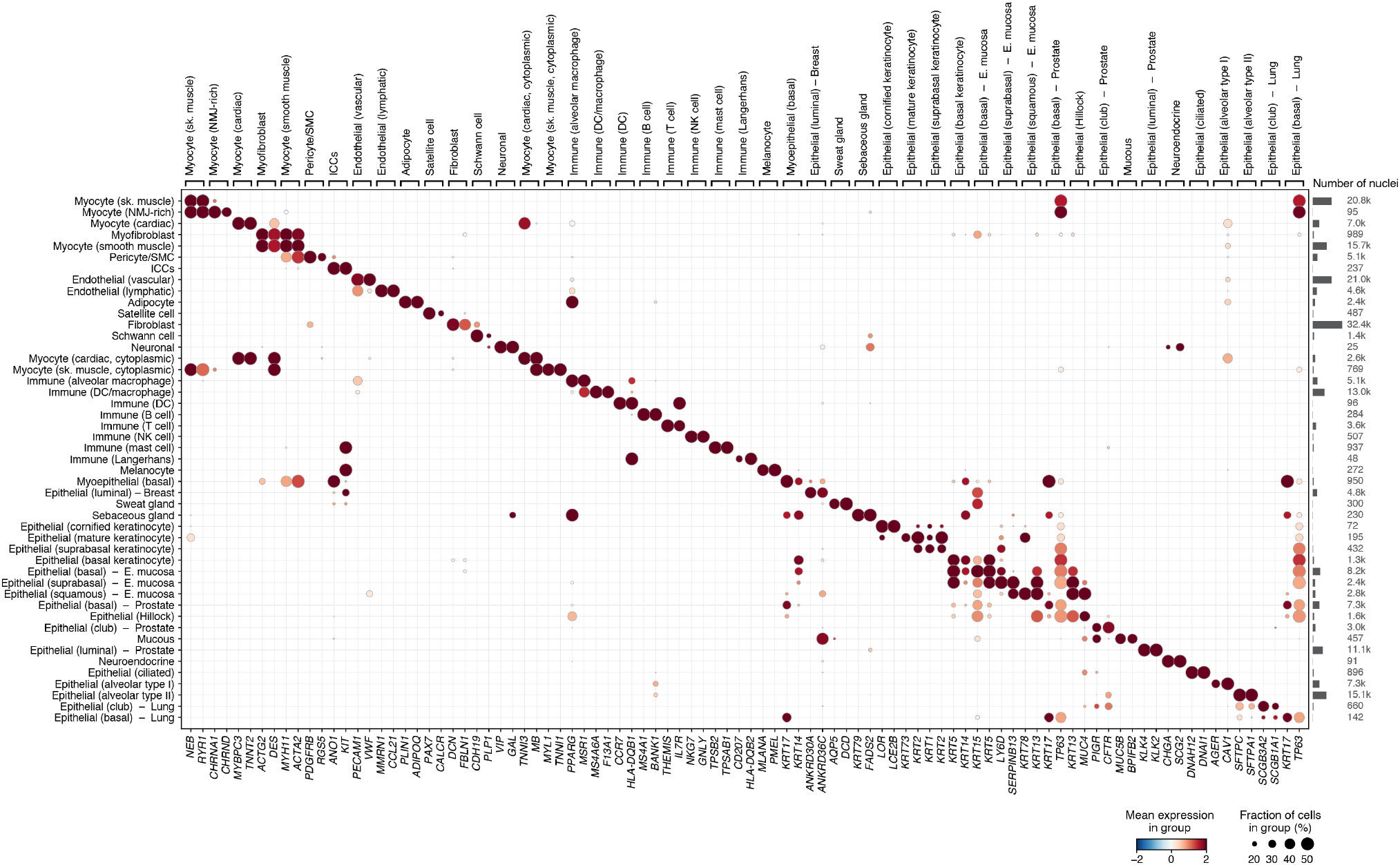
Broad cell type gene markers. Scaled mean expression (*z*-score, circle color) and fraction of expressing cells (circle size) of marker genes (columns, labels at bottom) associated with each granular cell subset (rows, and labels on top) with epithelial cells with tissue-specific markers shown separately. Right: number of nuclei in each cell subset.

**Fig. S6.**
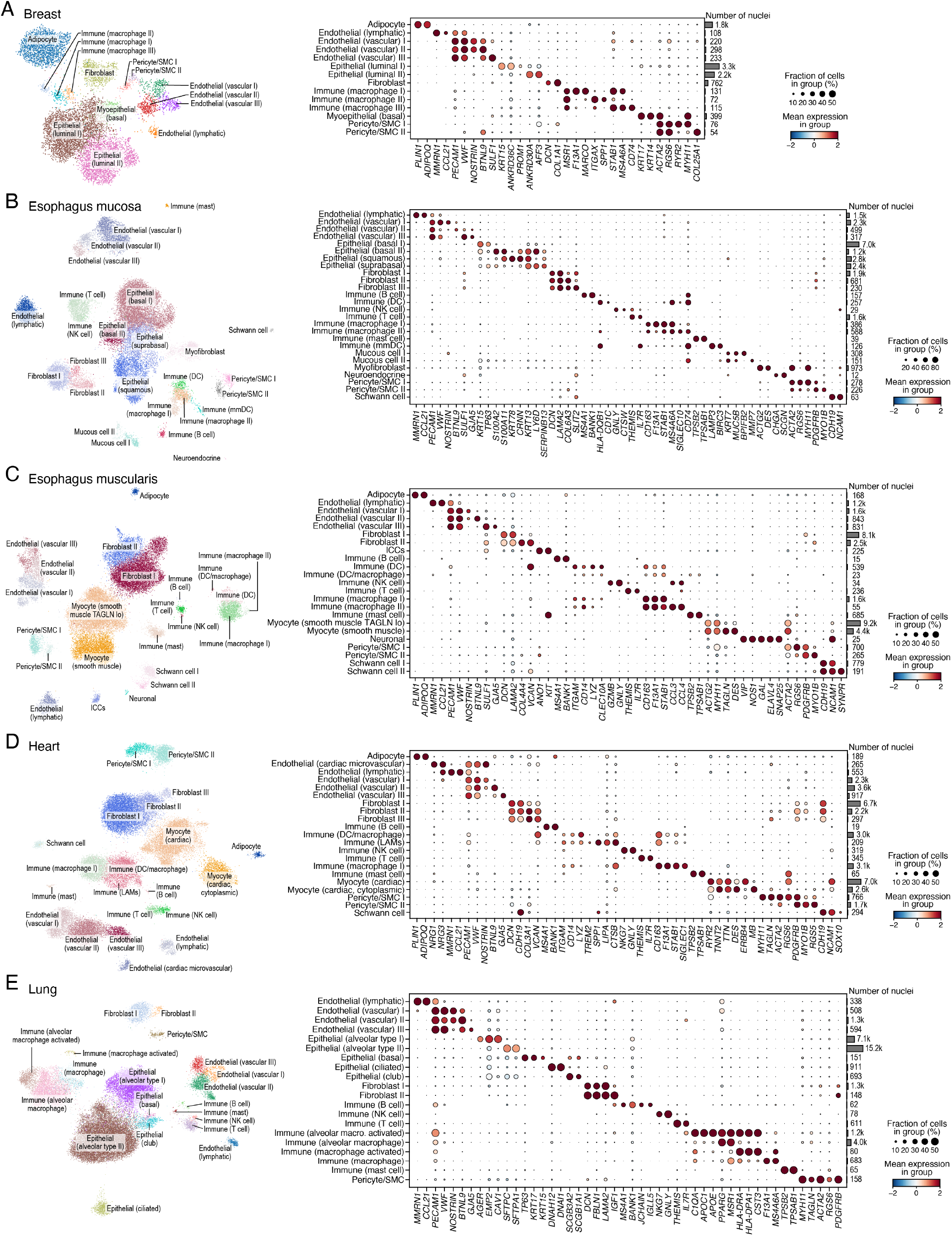
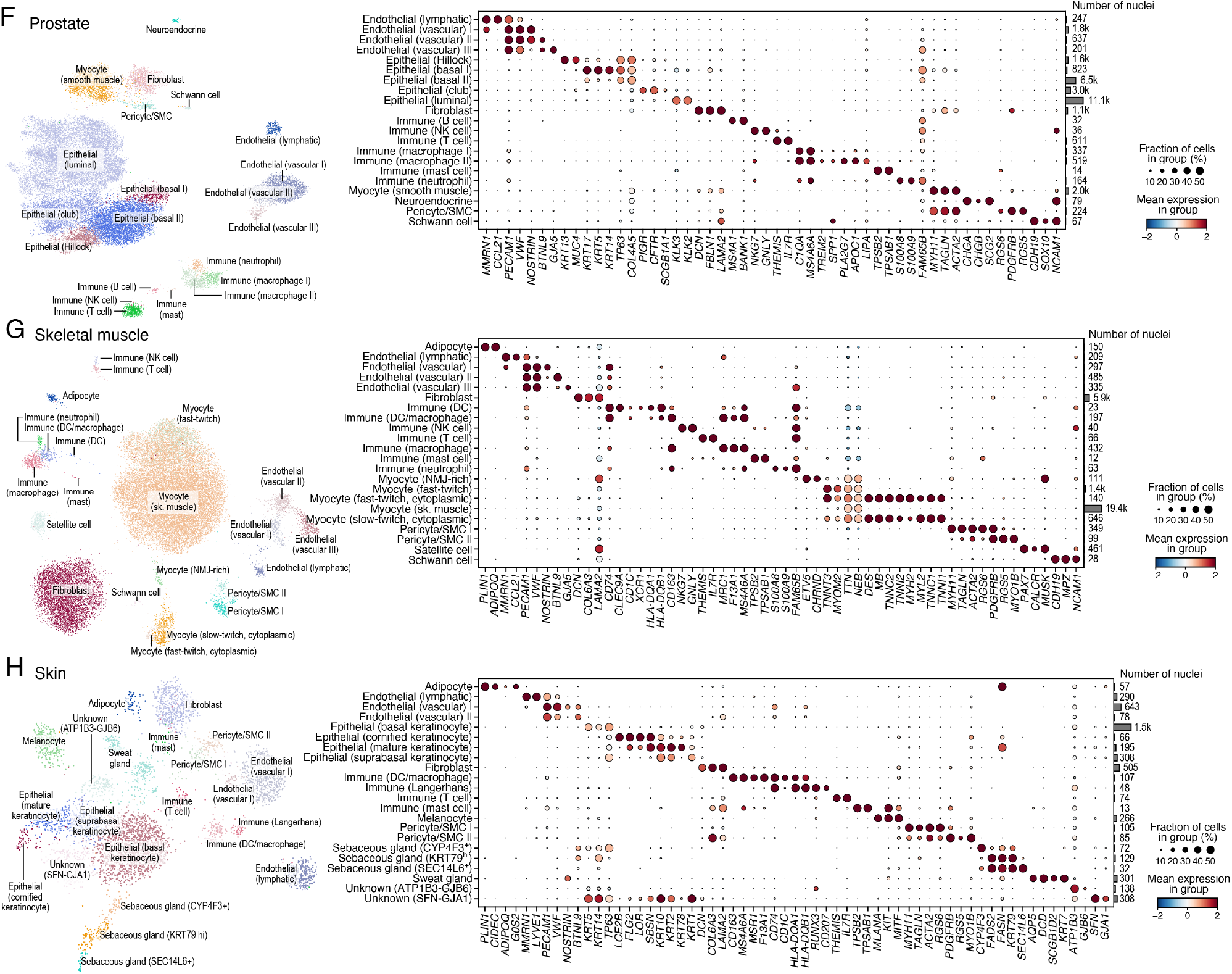
Granular cell subsets in each tissue. UMAP visualization (left) of single nucleus profiles (dots) colored by granular cell type annotation and scaled mean expression (*z*-score, circle color) and fraction of expressing cells (circle size) of marker genes (columns, labels at bottom) associated with those subsets (rows, with nuclei number on right and labels on top) for breast **(A)**, esophagus mucosa **(B)**, esophagus muscularis **(C)**, heart **(D)**, lung **(E)**, prostate **(F)**, skeletal muscle **(G)**, and skin **(H)**.

**Fig. S7.**
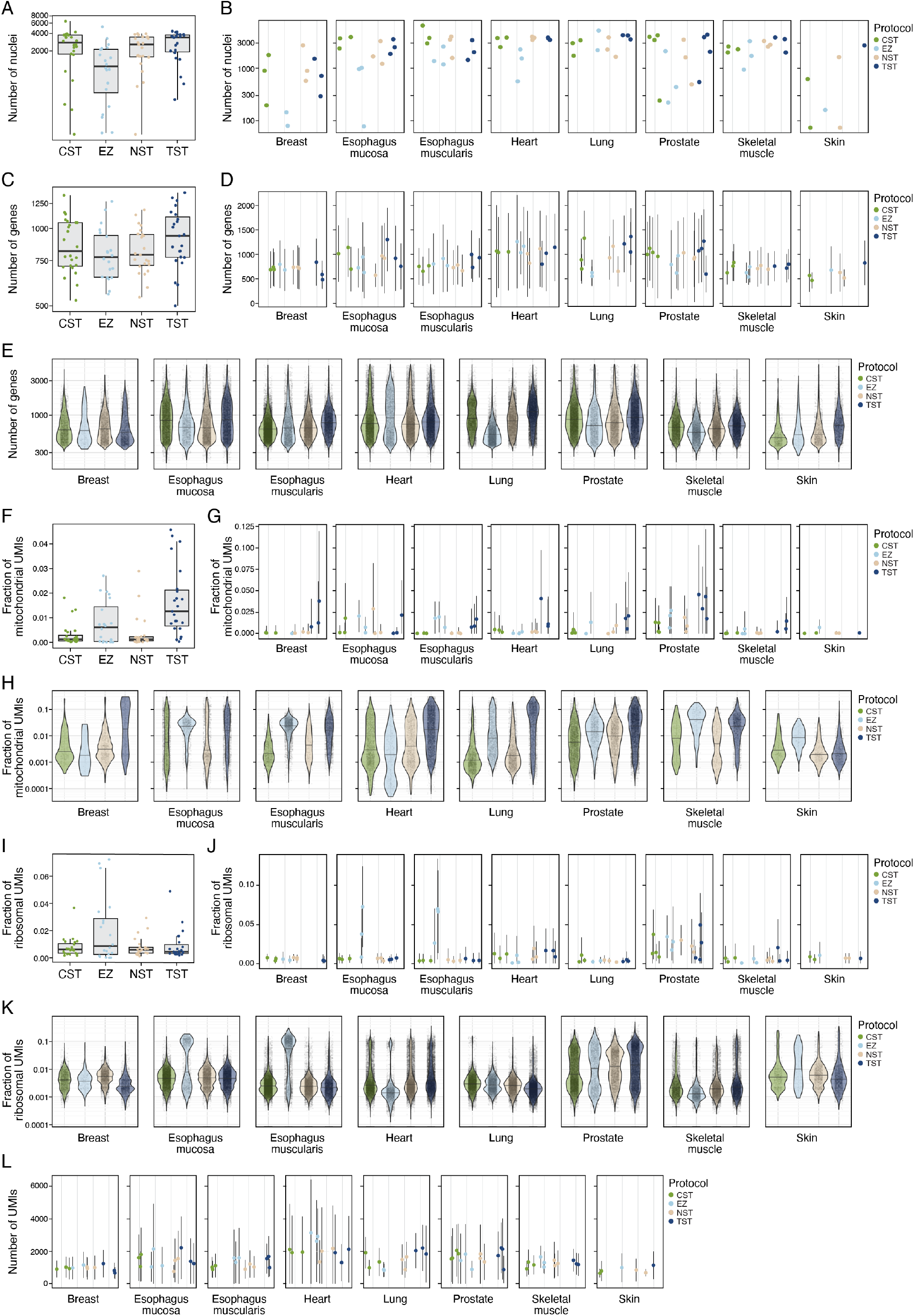
Relative benchmarking of four snRNA-seq protocols on different QC measures. **(A, B)** Number of recovered nuclei. Distribution (A) and number (B) of high-quality nuclei profiles (*y*-axis) recovered from each protocol in aggregate (A, *x*-axis) and for each sample in each tissue (B, *x*-axis). **(C–E)** Number of recovered genes. Distribution (C, E) and mean number (D) of genes (*y*-axis) recovered from each protocol in aggregate (C, *x*-axis), for each sample in each tissue (D, *x*-axis), and for each protocol in each tissue (E, *x*-axis). **(F–H)** Fraction of mitochondrial transcripts. Distribution (F, H) and mean (G) of the fraction of mitochondrial transcripts (*y*-axis, Unique Molecular Identifiers (UMIs)) recovered from each protocol in aggregate (F, *x*-axis), for each sample in each tissue (G, *x*-axis), and for each protocol in each tissue (H, *x*-axis). **(I–K)** Fraction of ribosomal transcripts. Distribution (I, K) and mean (J) of the fraction of ribosomal transcripts (*y*-axis, UMIs) recovered from each protocol in aggregate (F, *x*-axis), for each sample in each tissue (G, *x*-axis), and for each protocol in each tissue (H, *x*-axis). **(L)** Number of transcripts. Mean number of UMIs (*y*-axis) in each sample in each tissue. Box plots show median, quartiles, and whiskers at 1.5 IQR (interquartile range). The horizontal bar in violin plots represents the median. Error bars in (D,G,J,L) show one standard deviation above and below the mean.

**Fig. S8.**
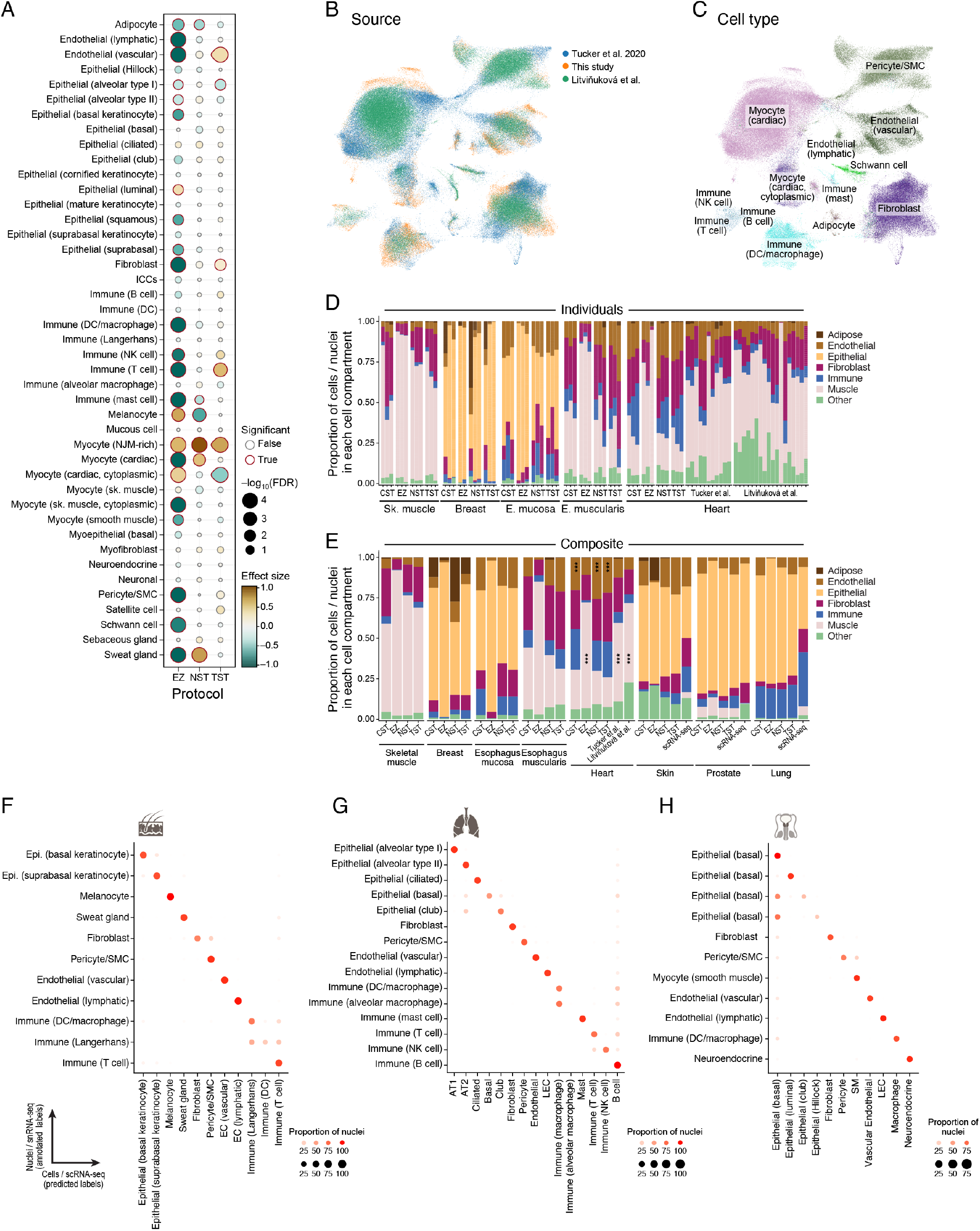
Benchmarking of snRNA-seq protocols by cell composition and comparison to scRNA-seq. **(A–E)** Impact of snRNA-seq protocol on recovered cell composition. (**A**) Significance (Benjamini–Hochberg FDR < 0.1; circle outline color) of the enrichment of each cell type (rows) in each protocol (columns) relative to the CST protocol across all tissues and samples after correction for tissue specific effects (**Methods**). **(B,C)** UMAP representation of snRNA-seq profiles (after batch correction by individual with Harmony (Korsunsky et al. 2019), **Methods**) colored by study **(B)** and cell type **(C)**. **(D)** Proportion of cells or nuclei (*y*-axis) across broad cell groups (color legend) in each sample (*x*-axis), stratified by tissue and protocol. (**E**) Proportion of cells or nuclei (*y*-axis) across broad cell groups (color legend) in each tissue (*x*-axis), stratified by tissue and protocol. Asterisks indicate significantly higher (muscle EZ, Tucker et al. and Litviňuková et al.) or lower (endothelial, Tucker et al. and Litviňuková et al.) proportions compared to the CST protocol (Dirichlet regression, Benjamini–Hochberg FDR < 0.01, **Methods**). **(F–H)** Agreement in cell-intrinsic expression profiles between scRNA-Seq and snRNA-Seq. Proportion of nuclei (circle size and color) of each subset (rows) that are predicted to be in each cell class (columns) by a random forest classifier trained on cells, for skin **(F),** lung **(G)** or prostate **(H)**.

**Fig. S9.**
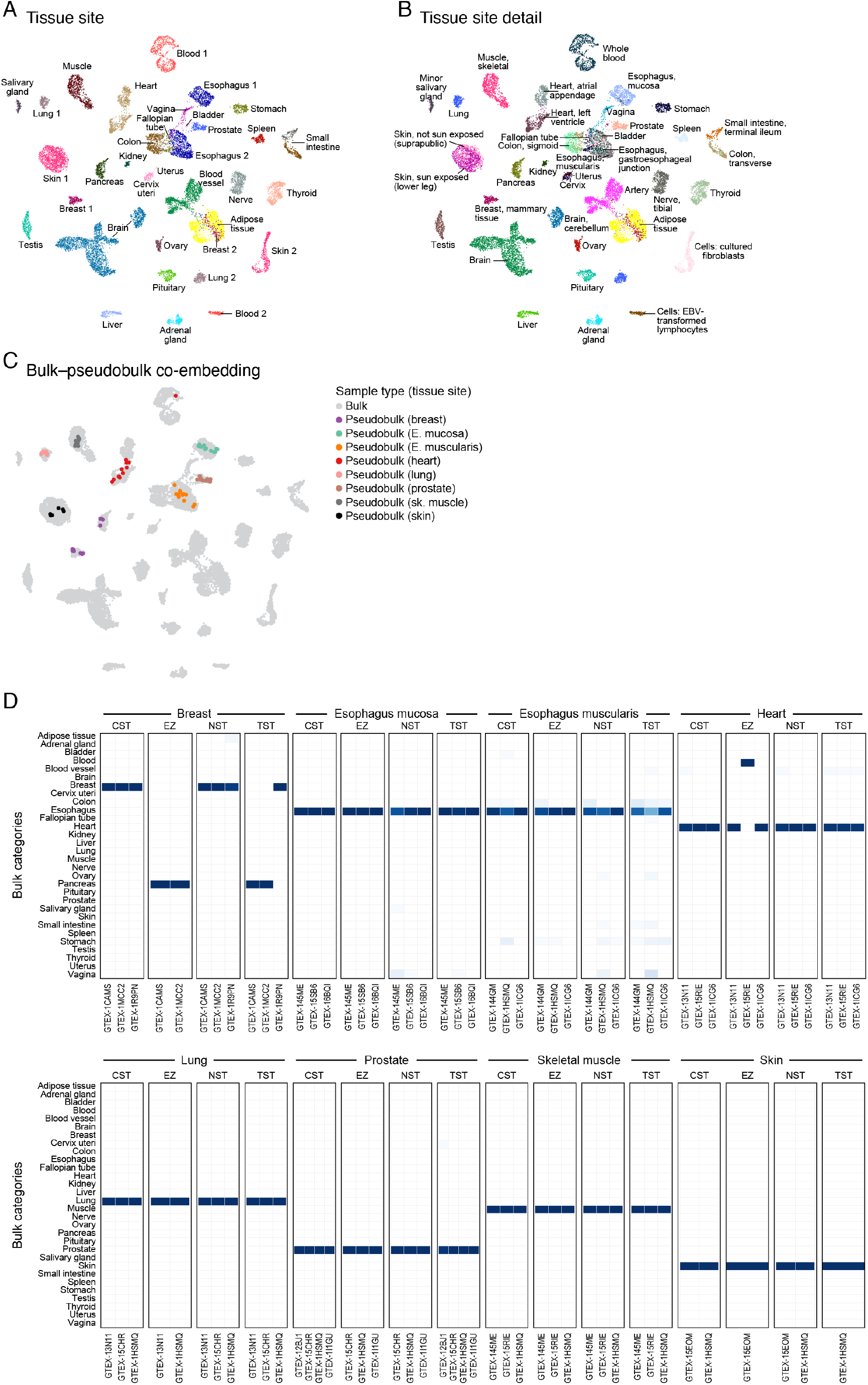
snRNA-Seq pseudobulk matches bulk RNA-seq of matched samples. (**A–C**) Co-embedding of bulk and pseudobulk RNA-seq profiles. UMAP representation of bulk RNA-seq and pseudobulk snRNA-seq samples (dots), with each sample colored by tissue site **(A)** detailed tissue site (**B**), or highlighting only the pseudobulk samples from each site **(C)**. **(D)** Co-embedding successfully places bulk and pseudobulk samples from the same tissue in proximity, as shown by the fraction of bulk nearest neighbors (color bar) of each pseudobulk sample (columns) that are derived from each tissue site (rows).

**Fig. S10.**
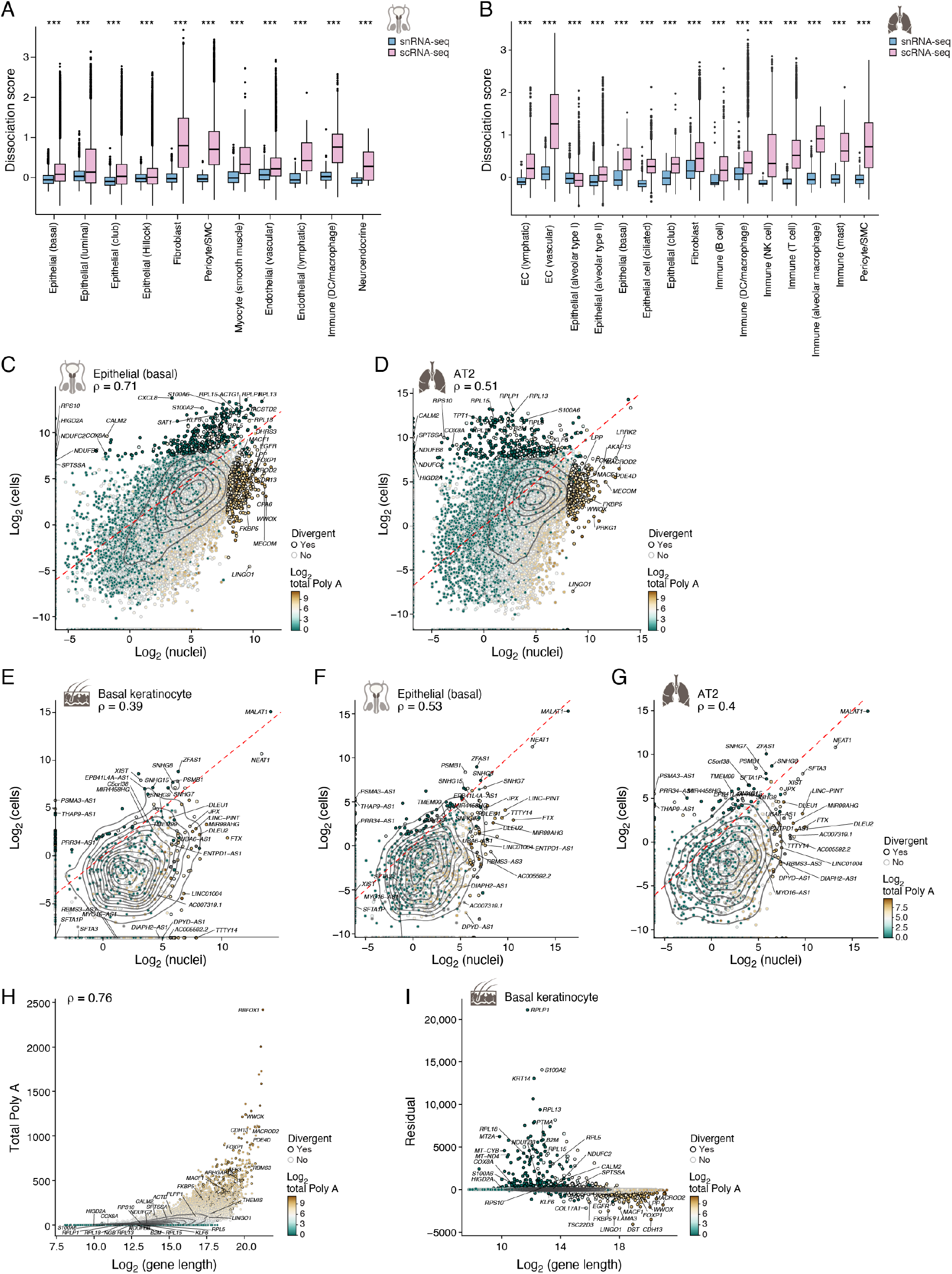
Differences in gene expression between snRNA-seq and scRNA-seq. **(A, B) Expression of tissue dissociation signatures in scRNA-seq but not snRNA-seq.** Distribution of the score (*y*-axis, average background corrected, log(TPX+1)) of a dissociation signature van den Brink et al. in scRNA-seq (pink) and snRNA-seq (blue) profiles in each major lung cell type category (*x*-axis). (*** Benjamini-Hochberg FDR < 10^−16^, Wilcoxon rank-sum test). Box plots show median, quartiles, and whiskers at 1.5 IQR (interquartile range). (**C–G**) Limited divergence of gene expression between scRNA-seq and snRNA-Seq. Averaged pseudobulk expression values (**Methods**) of genes (dots) in nuclei (*x*-axis) and cells (*y*-axis), shown for protein-coding genes in (**C**) prostate epithelial (basal) cells **(D)** lung alveolar type II cells, and long non-coding RNA genes in (**E**) skin basal keratinocytes, (**F**) prostate epithelial cells (basal), and (**G**) lung alveolar type II cells. Divergent genes (dot outline color) deviating from straight line regression fit by size of residuals (**Methods**). Color scale: total length of polyA stretches with at least 20 adenine bases. (**H, I**) Relation between gene expression differences in nuclei *vs*. cells, gene length and polyA stretches. **(H)** The number of polyA stretches (*y*-axis) and length (*x*-axis) of each gene. (**I**) Divergence (*y*-axis, residual of straight-line regression fit) between pseudobulk gene expression of single cell and single nucleus RNA-seq and gene length (*x*-axis) for each protein coding gene expressed in skin basal keratinocytes in both datasets.

**Fig. S11.**
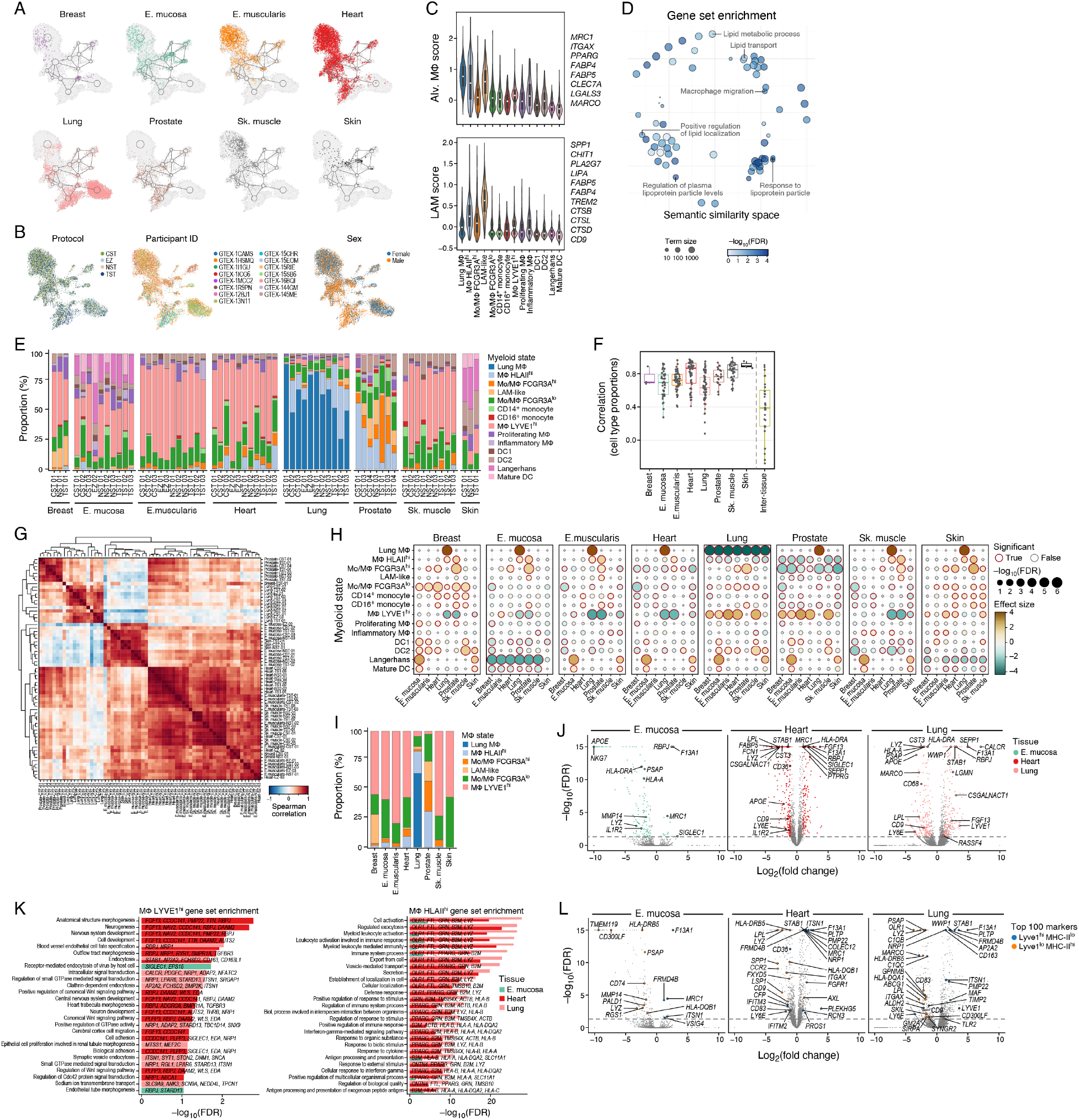
Cross-tissue analysis of myeloid cell subsets. **(A, B)** Myeloid cell subsets across tissues. UMAP visualization of myeloids snRNA-seq profiles, highlighting cells from each tissue **(A)** or colored by protocol, donor, or sex **(B)**. **(C)** Alveolar and lipid-associated macrophage signatures. Distribution of signature scores (*y*-axis) of alveolar macrophages (top) and lipid-associated macrophage (LAM) (bottom) for each myeloid cell subset (*x*-axis). Signature genes are listed on the right. **(D)** Key processes enriched in the LAM-like signature. Semantic similarity space of functional gene sets (circles; circle size proportional to gene set size) preserving distances between similar GO terms colored by enrichment (–log_10_(Benjamini– Hochberg FDR), Fisher’s exact test) of LAM-like markers. **(E)** Myeloid cell proportions across tissues. Proportion (%, y axis) of different myeloid subsets in each sample (*x*-axis). **(F, G)** Myeloid cell composition is similar in the same tissue and different between tissues. (**F**) Distribution of pairwise Spearman correlation coefficients (*y*-axis) of myeloid cell subset proportion profiles for samples within each tissue and between different tissues (*x*-axis). Box plots show median, quartiles, and whiskers at 1.5 IQR (interquartile range). **(G)** Spearman correlation coefficient (color bar) of myeloid cell subset proportion profiles of each pair of samples (columns, rows). Rows and columns were hierarchically clustered using the Euclidean distance and complete linkage. **(H)** Differences in myeloid cell distributions across tissues. Significance (circle size, –log_10_(FDR)) and effect size (circle color) of the difference in proportion in each myeloid cell subset (rows) between each tissue (columns) relative to one reference tissue (label on top). **(I)** Macrophage cell proportions across tissues. Proportion (%, y axis) of different macrophage subsets in each individual tissue (*x*-axis). **(J-L)** Mɸ *LYVE1*^hi^ and Mɸ *HLA*II^hi^ distinctive signatures. (**J**) Difference in expression (*x*-axis, log_2_(Fold change), *x* < 0: enriched in Mɸ *HLA*II^hi^ *x* < 0: enriched in Mɸ *LYVE1*^hi^) and its significance (– log_10_(FDR (Benjamini-Hochberg)), Wald test) between Mɸ *LYVE1*^hi^ vs. Mɸ *HLA*II^hi^ profiles in three tissues where both populations are observed. **(K)** Significance of enrichment (*x*-axis, –log_10_(Benjamini-Hochberg FDR), Fisher’s exact test) of GO gene sets (*y*-axis) in genes differentially expressed between Mɸ *LYVE1*^hi^ and Mɸ *HLA*II^hi^ populations in the three tissues in **J**. **(L)** Differential expression plots as in (J) with marker genes of mouse *Lyve1*^hi^MHCII^lo^ and *Lyve1*^lo^MHCII^hi^ cells (Chakarov et al.) highlighted.

**Fig. S12.**
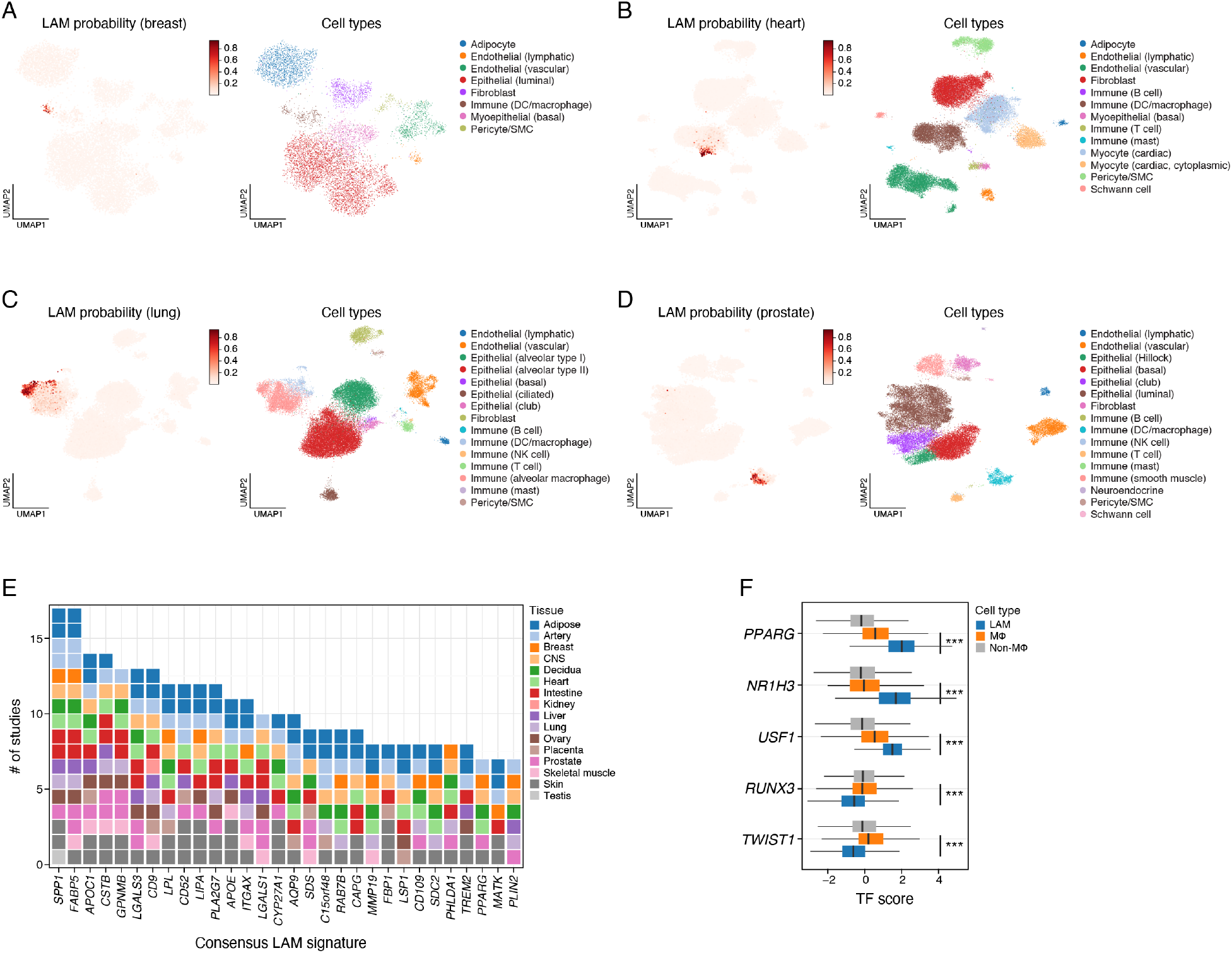
Tissue distribution of LAM-like cells. **(A–D)** UMAP visualization of nuclei profiles from breast **(A)**, heart **(B)**, lung **(C)** and prostate **(D)** (where LAM-like profiles are detected), colored by classification probabilities of LAMs (left) or by broad cell type annotations (right). **(E)** Consensus LAM signature. The number of published studies (*y*-axis) and tissue type (color) in which each LAM marker gene (*x*-axis, **Methods**) is detected as a marker. **(F)** LAM-associated TFs. Distribution of activity scores (*x*-axis) of TFs (*y*-axis) that are significantly high in LAMs or non-LAM macrophages. *** = *p*-value < 10^− 20^ (*p*-values in Figure 3I are combined across studies using Fisher’s method).

**Fig. S13.**
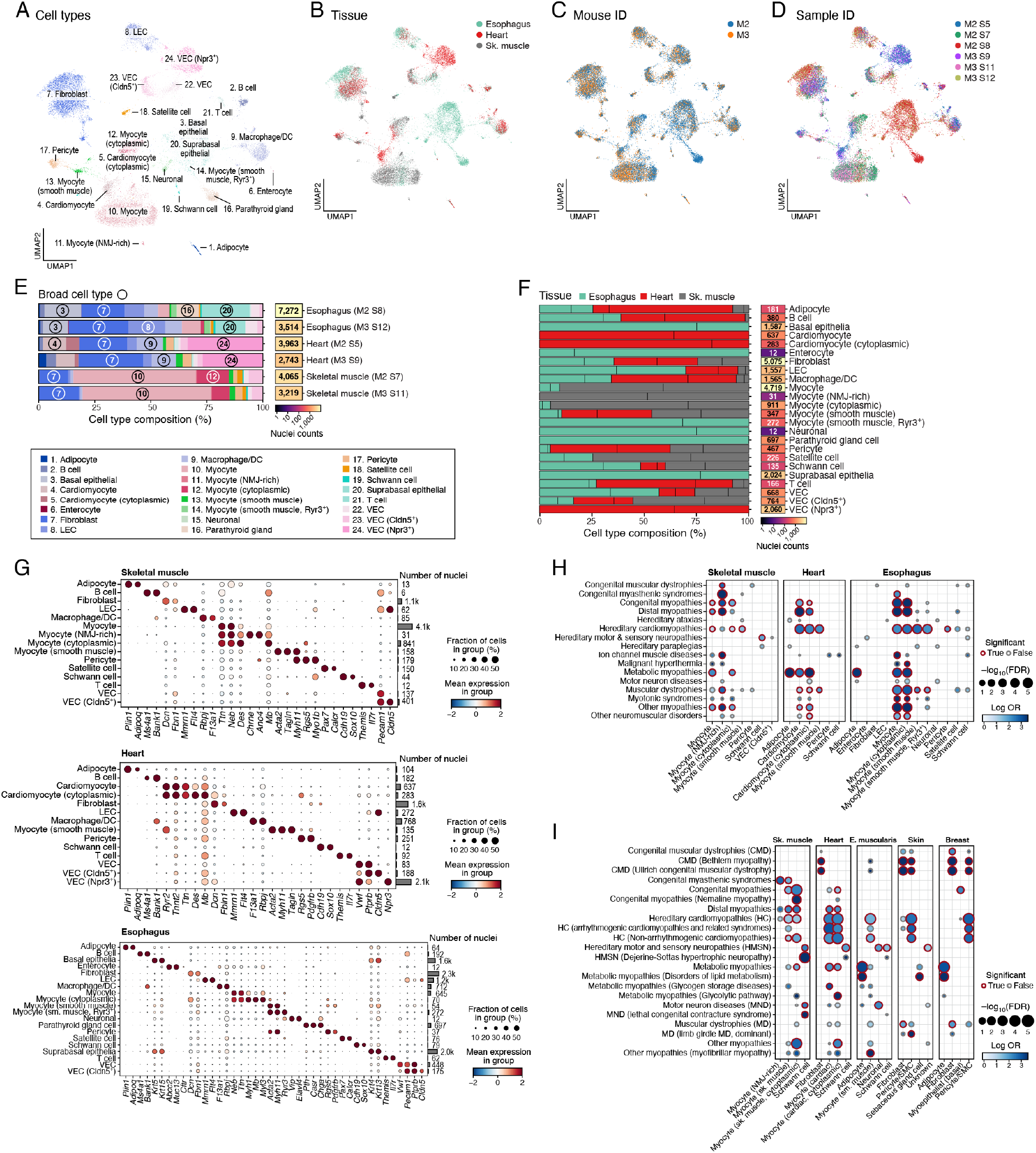
Analysis of monogenic muscle disease gene in mouse muscle tissues. (**A–D**) Mouse muscle tissue atlas. UMAP visualization of snRNA-seq profiles from skeletal muscle, heart and esophagus in mouse, colored by cell type **(A)**, tissue **(B)**, mouse (**C**) or sample IDs **(D)**. (**E, F**) Mouse muscle atlas cell type composition. Proportion of nuclei (*x*-axis) of each type (color) in each tissue (sample) (**E**), and of each tissue in each cell type (**F,** after normalizing for the total number of nuclei profiled in each tissue, **Methods**) along with the number of profiled nuclei (right). Circled numbers refer to cell types in the color legend. Black vertical lines in bars in F: relative proportion from each mouse. **(G)** Cell type markers. Scaled mean expression (dot color, *z-*score) and fraction of expressing cells (dot size) for marker genes (columns) in each cell subset (rows). (**H, I**) Enrichment of broad cell type markers with monogenic muscle disease genes. Effect size (log odds ratio, dot color) and significance (–log_10_(Benjamini–Hochberg FDR), dot size) of enrichment of mouse orthologs (**Methods**) of genes from each monogenic muscle disease group (rows) for (**H**) cell type markers of broad cell subsets (columns) in each mouse tissue (**I**) or from human muscle tissues (as in Figure 4A) as well as skin and breast. Red border: Benjamini–Hochberg FDR < 0.1.

**Fig. S14.**
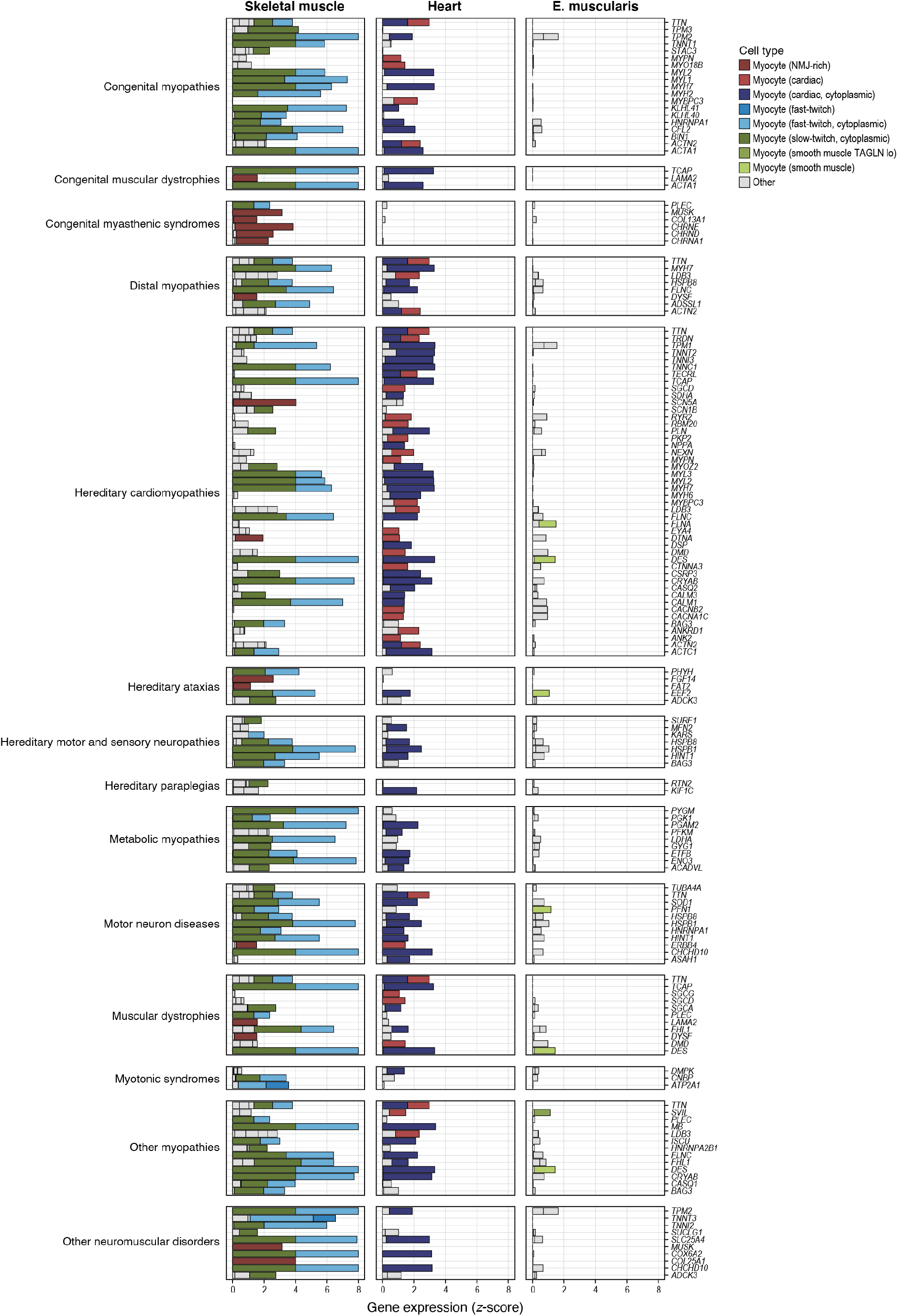
Expression of monogenic muscle disease genes highly expressed in myocytes in three muscle tissues. Expression (*z*-score, *x*-axis) of monogenic muscle disease genes (*y*-axis) that are highly expressed in myocytes ordered by disease category (labels on left) in each cell type (color legend) in each of three mouse tissues (labels on top). Grey (“other”): Cell types with low expression of the indicated genes.

**Fig. S15.**
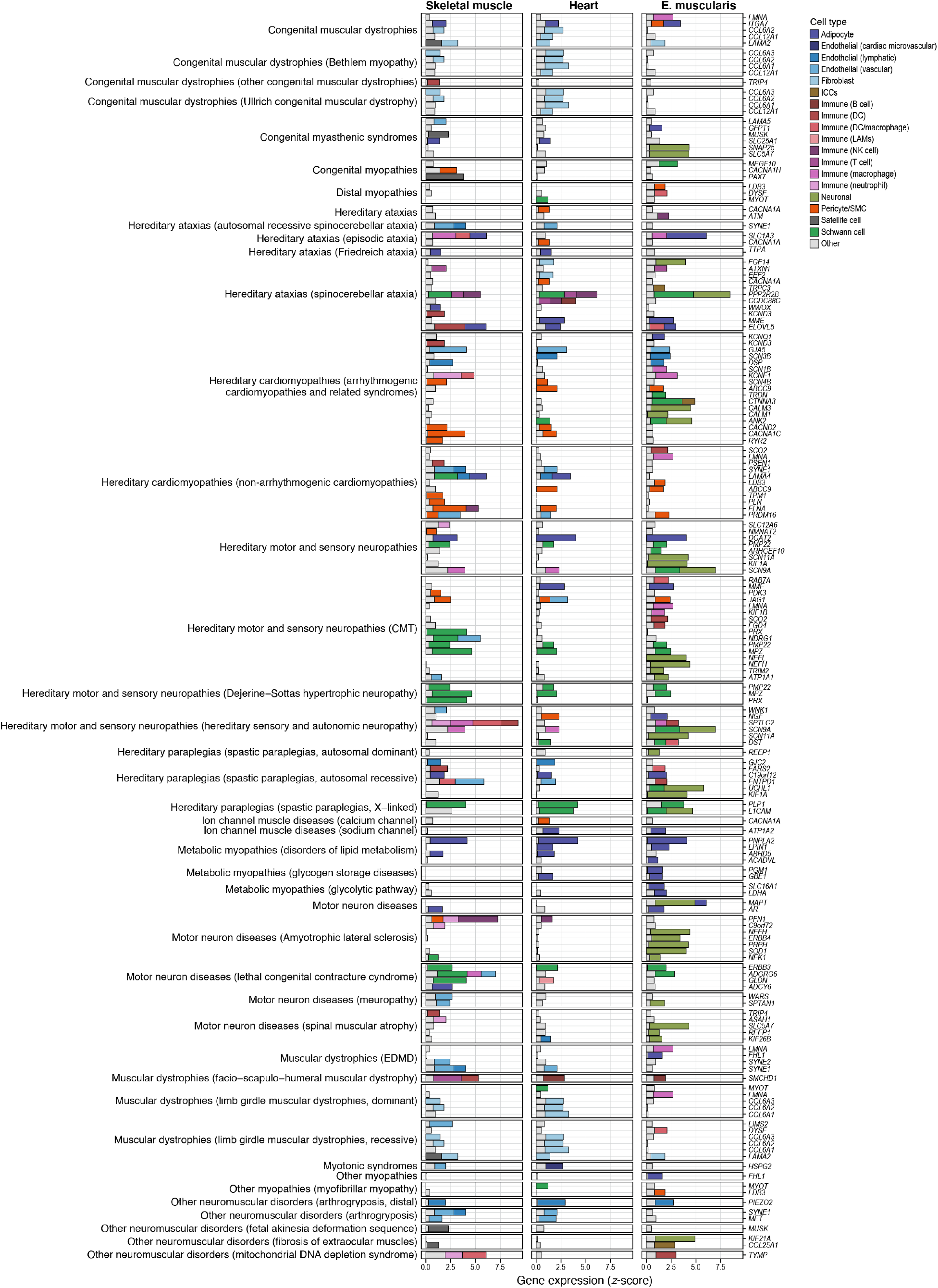
Expression of monogenic muscle disease genes in non-myocytes in three muscle tissues, including genes expressed also in myocytes. Expression (*z*-score, *x*-axis) of monogenic muscle disease genes (*y*-axis) expressed in non-myocytes (and possibly in myocytes too) ordered by disease category (labels on left) in each cell type (color legend) in each of three mouse tissues (labels on top). Grey (“other”): Cell types with low expression of the indicated genes.

**Fig. S16.**
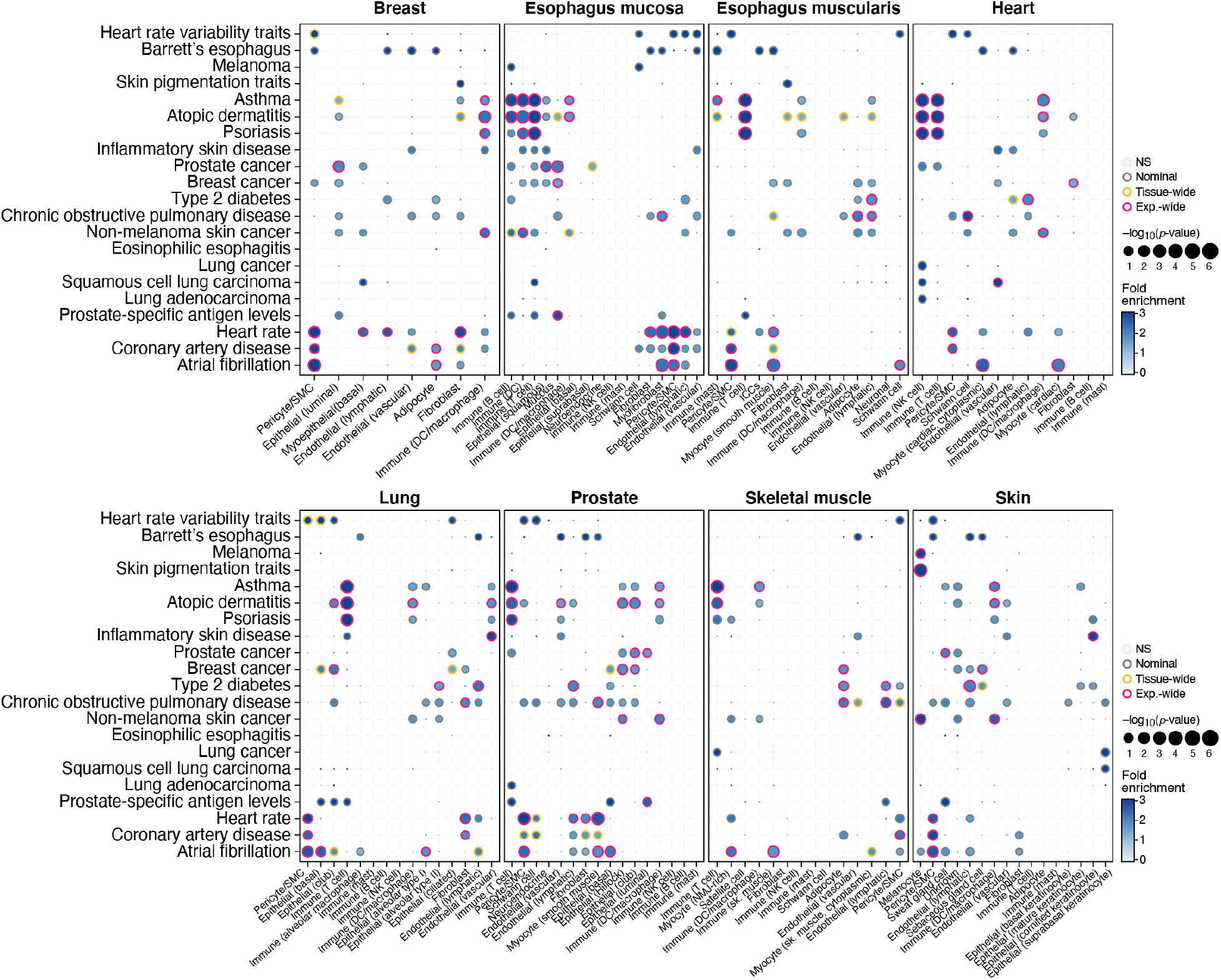
Cell type enrichment of GWAS loci from 21 complex traits in 8 tissues using broad cell type annotations. Significance (circle size, –log_10_(*p*-value)) and effect size (circle color, fold-enrichment) of enrichment of GWAS locus sets of complex traits (rows) in each broad cell type category (columns) in the eight tissues in the cross-tissue atlas (panels). Grey, orange, red borders: nominal, tissue-wide (Benjamini–Hochberg (BH) FDR < 0.05 correcting for all cell types tested per tissue and per trait) and experiment-wide (BH FDR < 0.05 correcting for all cell types tested across 8 tissues and 21 traits) significance results, respectively.

**Fig. S17.**
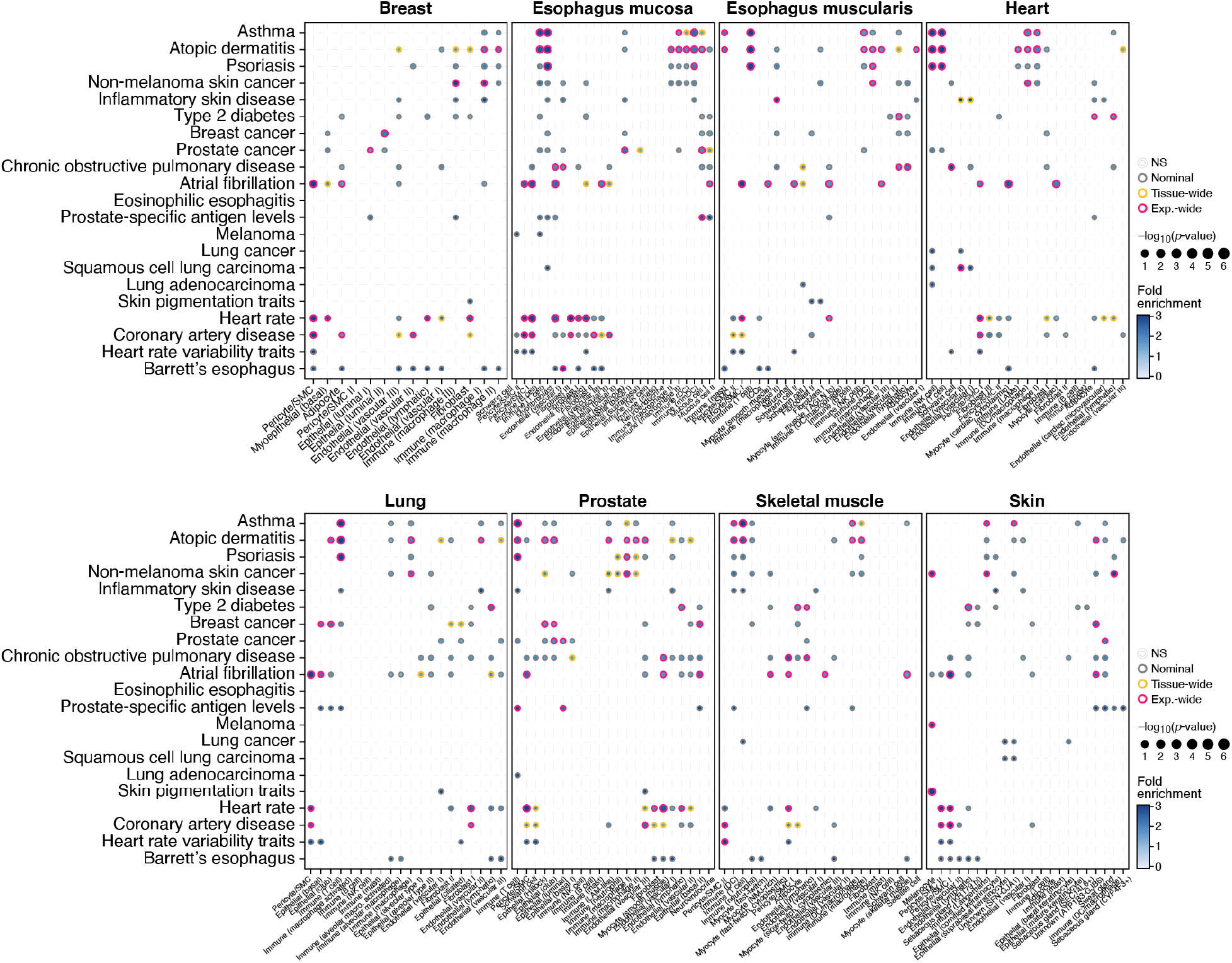
Cell type enrichment of GWAS loci from 21 complex traits in 8 tissues using granular cell type annotations. Significance (circle size, –log_10_(*p*-value)) and effect size (circle color, fold-enrichment) of enrichment of GWAS locus set (rows) in each granular cell type category (columns) in the eight tissues in the cross-tissue atlas (panels). Grey, orange, red borders: nominal, tissue-wide (Benjamini–Hochberg (BH) FDR < 0.05 correcting for all cell types tested per tissue and trait) and experiment-wide (BH FDR < 0.05 correcting for all cell types tested across 8 tissues and 21 traits) significance results.

**Fig. S18.**
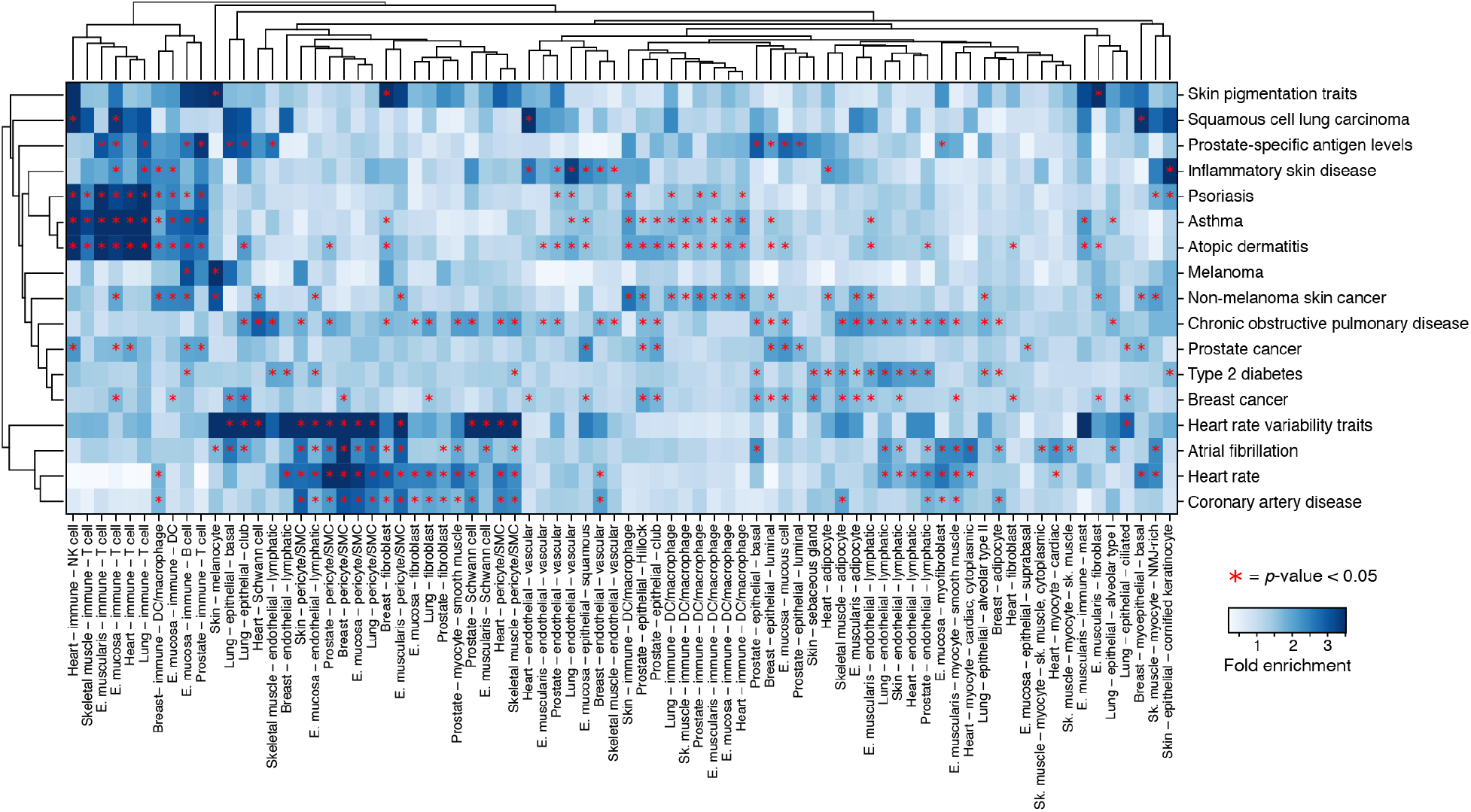
Cell type specificity enrichment of 21 complex traits across broad cell types from 8 tissues. Fold-enrichment (color bar) of each GWAS locus set (rows) for specificity in each broad cell type category (columns). Red stars: nominal significance (*P* < 0.05). Only traits with at least one enrichment at tissue-wide significance (BH FDR < 0.05 correcting for number of cell types tested per tissue per trait) are shown. Rows and columns are hierarchically clustered using the Euclidean distance and average linkage method. Red stars denote nominal significance.

**Fig. S19.**
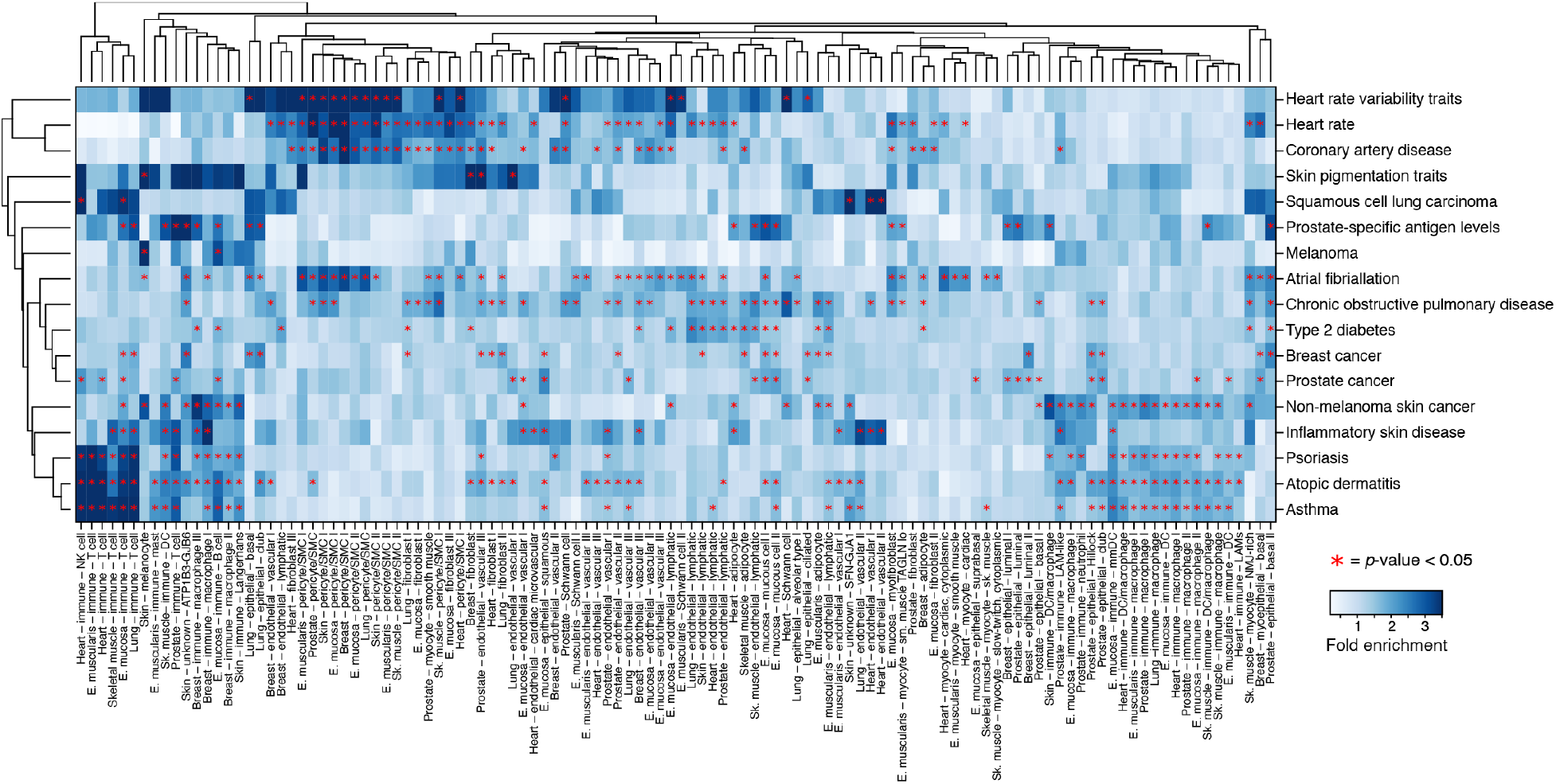
Cell type specificity enrichment of 21 complex traits across granular cell types from 8 tissues. Fold-enrichment (color bar) of each GWAS locus set (rows) for specificity in each granular cell type category (columns). Red stars: nominal significance (*P* < 0.05). Only traits with at least one enrichment at tissue-wide significance (BH FDR < 0.05 correcting for number of cell types tested per tissue per trait) are shown. Rows and columns are hierarchically clustered using the Euclidean distance and average linkage method. Red stars denote nominal significance.

**Fig. S20.**
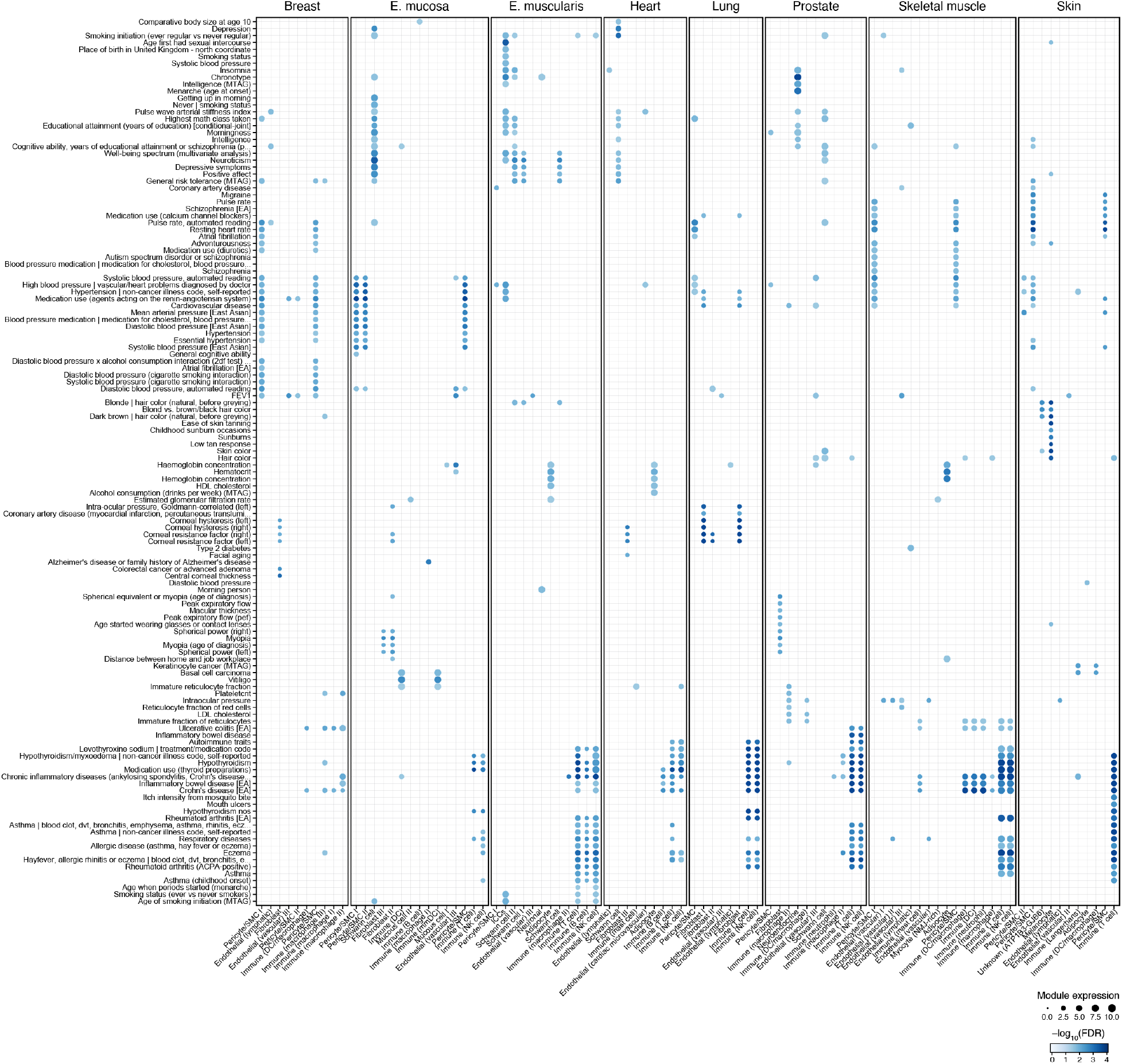
Module-based GWAS gene set enrichment analysis highlights associations between cell types and traits/diseases. Significance (–log_10_(Benjamini–Hochberg FDR), dot size) of enrichment of GWAS module (rows) expression (dot color) in each cell type (columns).

**Fig. S21.**
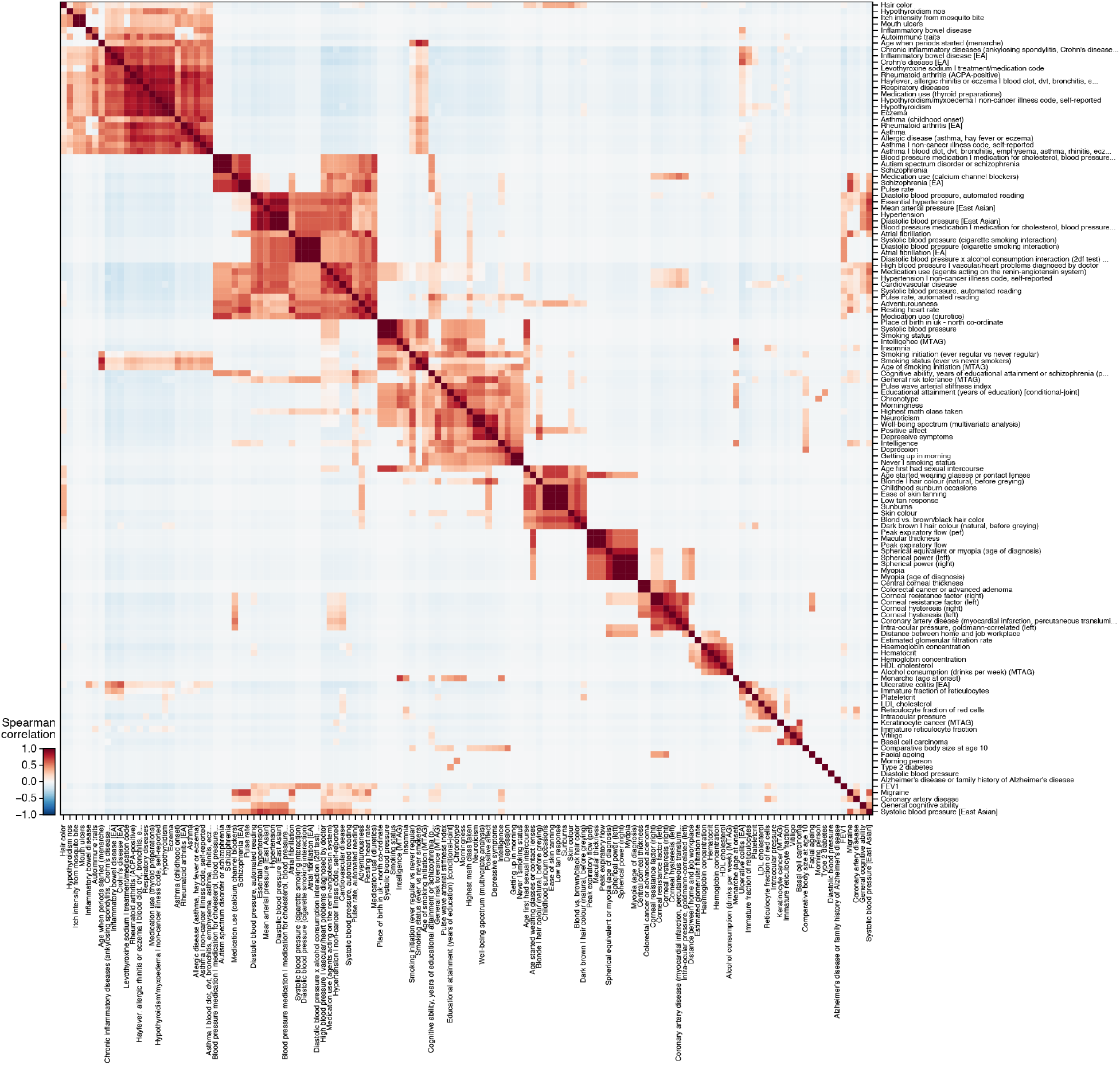
Disease/trait grouping by GWAS-cell type enrichments. Similarity (Spearman correlation coefficient, color bar) between each pair of GWAS traits/diseases (rows, columns) based on their enriched cell type profiles (as in Figure S20).

**Fig. S22.**
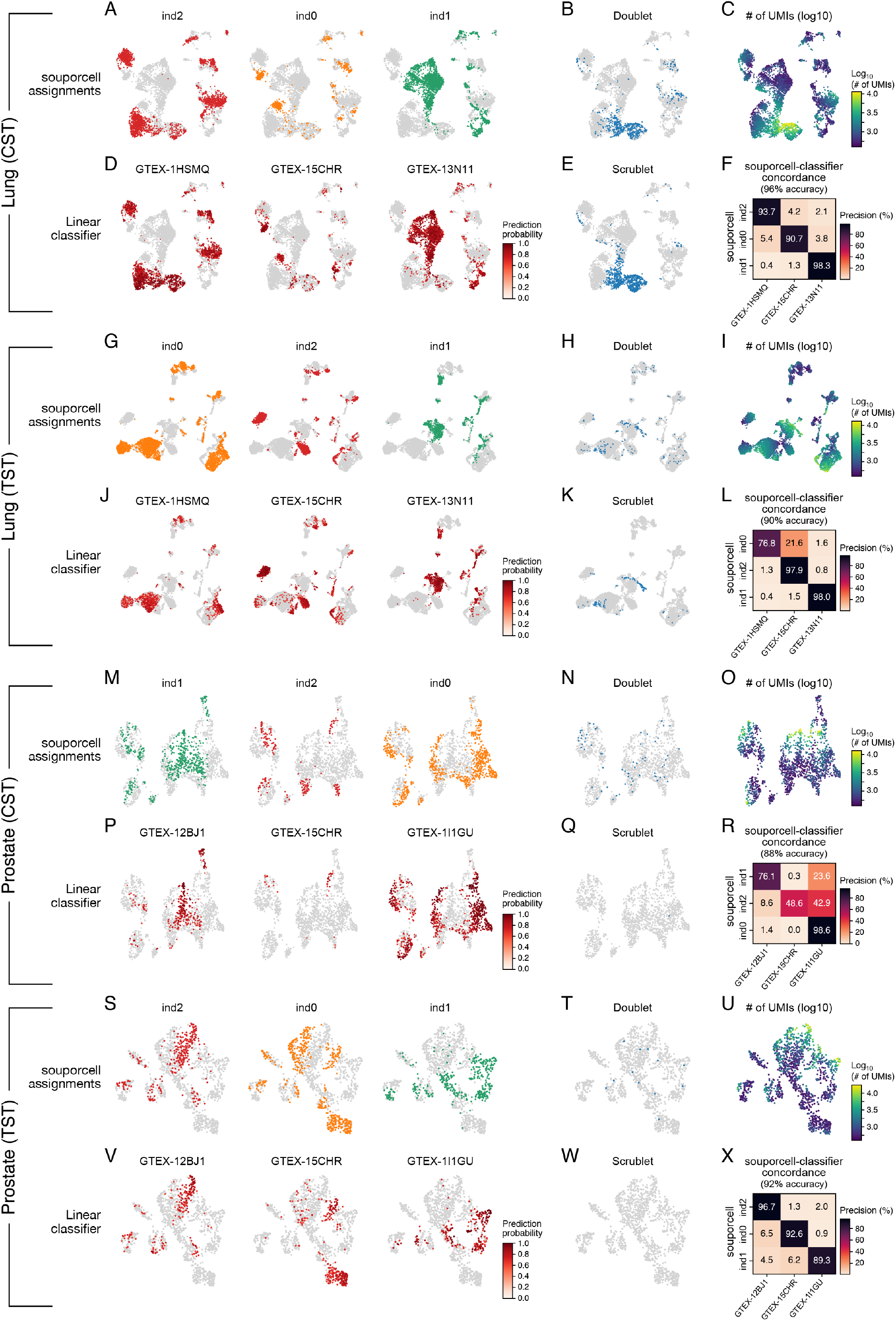
Successful processing of multiplexed samples by snRNA-Seq. Analysis of snRNA-Seq profiles from frozen tissue samples from 3 individuals processed in multiplex for lung (**A–L**) and prostate (**M–X**) by either the CST (**A–F, M-R**) or the TST (**G–L, S–X**) protocols. UMAP visualizations of nuclei profiles colored by the demultiplexing assignments of each nucleus by souporcell to individuals **(A,G,M,S)**, doublets **(B,H,N,T)**, log_10_-transformed total number of UMIs (**C,I,O,U)**, demultiplexing predictions of an expression-based linear classifier **(D,J,P,V)**, and doublets predicted by Scrublet **(E,K,Q,W)**. Heat maps (**F,L,R,X**) show the concordance (% precision, color bar) between souporcell (rows) and the linear classifier (columns) calls. Accuracy values are labeled on top.

